# A non-canonical Hippo pathway represses the expression of ΔNp63

**DOI:** 10.1101/2023.02.13.528336

**Authors:** Ana Maria Low-Calle, Hana Ghoneima, Nicholas Ortega, Adriana M. Cuibus, Chen Katz, David Tong, Carol Prives, Ron Prywes

## Abstract

The p63 transcription factor, a member of the p53 family, plays an oncogenic role in squamous cancers, while in breast cancers its expression is often repressed. In the canonical conserved Hippo pathway, known to play a complex role in regulating growth of cancer cells, the protein kinases MST1/2 and LATS1/2 act sequentially to phosphorylate and inhibit the YAP/TAZ transcription factors. We found that in the MCF10A mammary epithelial cell line as well as in squamous and breast cancer cell lines, expression of ΔNp63 RNA and protein is strongly repressed by inhibition of the Hippo pathway protein kinases in a manner that is independent of p53. While MST1/2 and LATS1 are required for p63 expression, the next step of the pathway, namely phosphorylation and degradation of the YAP/TAZ transcriptional activators is not required for repression of p63. This suggests that regulation of p63 expression occurs by a non-canonical version of the Hippo pathway. We additionally identified additional genes that were similarly regulated suggesting the broader importance of this pathway. Interestingly, we observed that experimentally lowering p63 expression leads to increased YAP protein levels, thereby constituting a feedback loop. These results, which reveal the intersection of the Hippo and p63 pathways, may prove useful for the control of their activities in cancer cells.

**One Sentence Summary:** Regulation of p63 expression occurs by a non-canonical version of the Hippo pathway in mammary epithelial, breast carcinoma and head and neck squamous carcinoma cells

## Introduction

The p53 family member p63 is a transcription factor that was initially described to play a role in the embryonic development of the skin and glandular epithelial tissues such as the breast (*1, 2*). p63 has two major alternative isoforms at the 5’ end of the gene encoding TAp63 and the ΔNp63; and each of these have three additional splice variants at the 3’ end; α, β, and γ (*3*) thereby creating six p63 variants with distinct N-and C-termini. The ΔNp63α isoform is the predominant isoform expressed in normal stratified and glandular epithelial tissues (*4, 5*). p63 regulates the expression of epithelial-specific genes and regulates survival, cell adhesion, and keratin gene expression in breast epithelial cells and squamous cell carcinomas (SCC) (*6, 7*). The p63 locus is frequently amplified in primary lung and head and neck squamous cell carcinomas (*8*) and in a more recent study the expression of the ΔNp63 isoform was enriched across multiple types of SCC (*9*). Copy number gain at the p63 locus at Chromosome 3q28 was found in SCCs and showed a correlation with elevated expression of ΔNp63 (*9*).

In breast cancers, the role of p63 is more complex, suggesting it can possess either tumor suppressor or growth promoting activities. On the one hand p63 expression is often lower in breast tumors than in normal tissue and it has been suggested that p63 can play a role as a suppressor of epithelial to mesenchymal transition (EMT) and metastasis (*10, 11*). Yet p63 can also promote growth of basal-like breast cancer cells (*12, 13*).

Other genes that have been found to be frequently amplified in SCC are the Hippo pathway transcriptional activators YAP and TAZ (*9, 14*). Each of these have been found to induce growth in both breast cancer cells and SCC (*15*–*18*).

The Hippo pathway is a tumor suppressor signaling pathway that negatively regulates YAP and TAZ (*19, 20*). The upstream MST1 and 2 (Mammalian Sterile-like-20) protein kinases serve to phosphorylate and activate the LATS1 and 2 protein kinases (Large Tumor Suppressor Kinase) (*21*). Activated LATS1/2 phosphorylate the transcriptional activators YAP and TAZ which are then sequestered in the cytoplasm and targeted for degradation (*19, 20, 22, 23*). When LATS1/2 are inactivated, YAP and TAZ are translocated to the nucleus where they interact with TEAD transcription factors to induce expression of growth-inducing genes such as Connective Tissue Growth Factor (CTGF) and others (*24, 25*). Additional conserved components of this pathway are MOB1A and MOB1B which bind to LATS1/2 (*26, 27*) and SAV1 which binds to MST1/2 (*28*–*30*).

We previously found that oncogenic transformation of MCF10A cells with H-RAS or PIK3CA leads to the epithelial-mesenchymal transition (EMT) and repression of p63 expression (*10*). Suppression of p63 expression also induces EMT. We were therefore interested in cellular signaling pathways that might control p63 expression. In addition, suppression of p63 expression may be useful therapeutically in SCCs where its expression is elevated. Since both p63 and YAP/TAZ have been shown to play roles in breast and squamous cancers, we considered the possibility of a connection between Hippo pathway components, YAP/TAZ, and p63. Our research has revealed unexpected links between p63 and this pathway which are described in this study.

## Results

### The Hippo pathway kinases MST1 and MST2 regulate the levels of p63

Given the increased expression of p63 in squamous cell carcinomas and its reduced expression in breast carcinomas, we were interested in how p63 might be regulated. The Hippo signaling pathway interacts in multiple ways with members of the p53 tumor suppressor family (*31*), which provided the impetus to evaluate whether pharmacological inhibition of the Hippo pathway had any effect on expression of p63. For this purpose we used the immortalized untransformed mammary epithelial cell line MCF10A (*32*), that we previously showed expresses mainly the ΔNp63α isoform of p63 (*10*). To pharmacologically inhibit the pathway we used XMU-MP1, a small molecule inhibitor of the uppermost protein kinases in the pathway, namely MST1 and MST2 (*33*). After treatment of the MCF10A cells with XMU-MP1 for 24 hours we observed a significant downregulation of p63 expression at the mRNA and protein levels (Fig. 1A, B). As expected (*23, 33*), inhibition of MST1/2 also led to a decrease in phospho-YAP levels and an increase in total YAP levels (Fig.1A). TAZ and phospho-TAZ levels were similarly regulated. The upregulation of YAP and TAZ proteins along with decreased phosphorylation at the LATS target sites on YAP (Ser^127^) and TAZ (Ser^89^) indicated that the kinase activity of LATS was suppressed due to the MST1/2 inhibitor (Fig. 1A). In addition, expression of the well-known YAP/TAZ target gene CTGF was strongly induced (Fig. 1A, B) (*24, 25*). To confirm that the effect of the inhibitor was through MST1/2, we used siRNA to suppress the levels of the kinases MST1 and MST2 individually and together and observed an increase in the protein levels of YAP and TAZ and upregulation of CTGF in all three conditions compared to the control siRNA (Fig. 1C, D). CTGF induction was the highest with MST2 knockdown while p63 suppression was slightly stronger with MST1 siRNA (Fig. 1C, 1D). These results further demonstrate that MST1/2 are required for p63 expression.

**Fig. 1.**
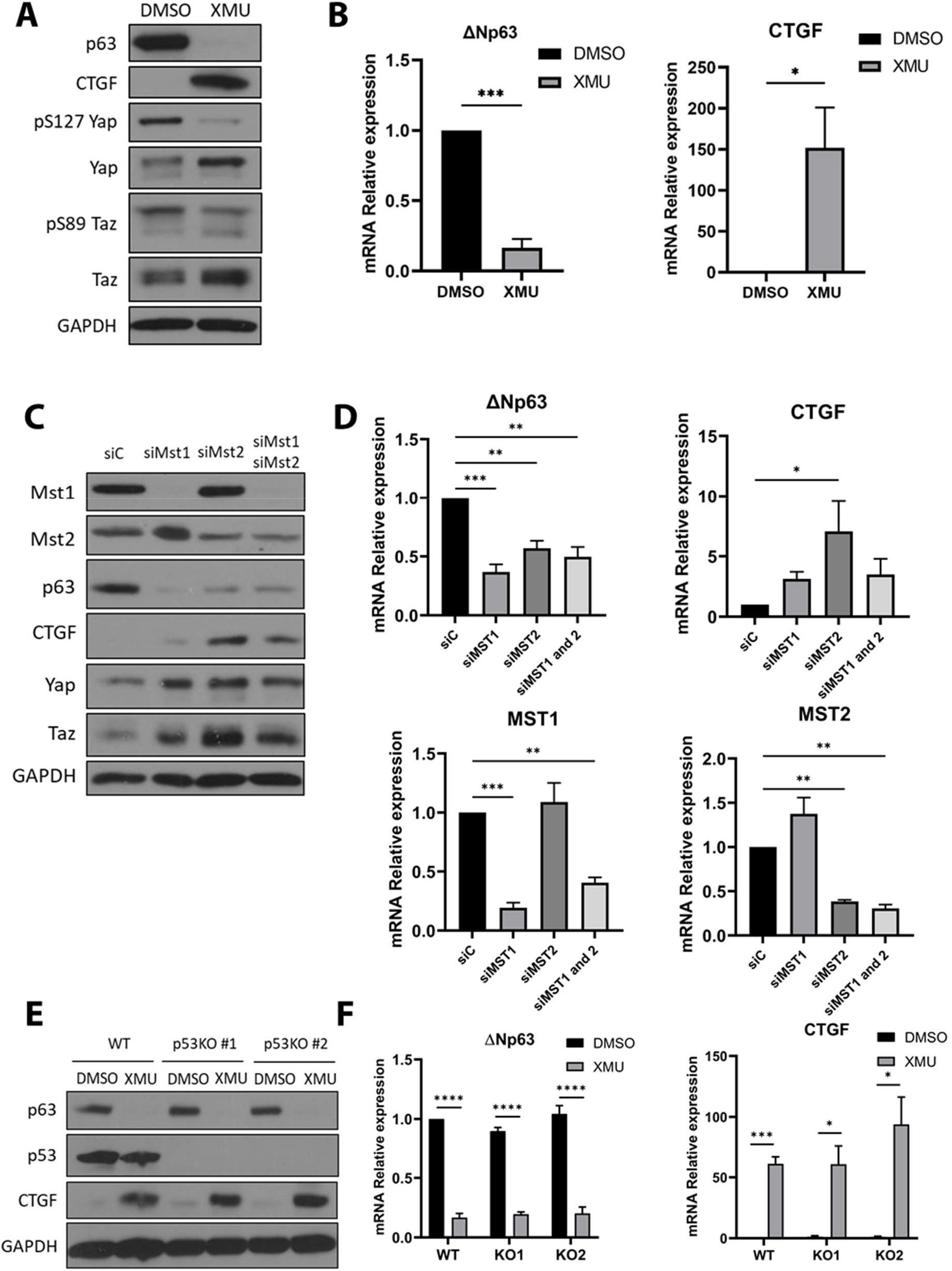
The hippo pathway Kinases MST1 and MST2 regulate expression of p63. **A**. Immunoblot of MCF10A cells treated with 5 µM of the MST1/2 inhibitor XMU-MP1 for 24h. Primary antibodies were used to the indicated proteins or their phosphorylated forms. **B**. RT-qPCR of MCF10A cells treated as in **A**. Bars represent the mean ± SEM of three independent experiments; for CTGF four independent experiments were done. Student’s t-test was performed. **C**. Immunoblot of MCF10A cells transfected with Smartpool siRNA against MST1, MST2 or both for 72h. **D**. RT-qPCR of MCF10A cells treated as in C. ANOVA with pairwise comparisons against the siC samples were performed. **E**. Immunoblot of MCF10A WT parental cells and two different p53 CRISPR null clones treated with 5 µM of XMU-MP1 or DMSO for 24h. **F**. RT-qPCR of the cell lines shown in **E**. Student’s t-tests were performed. Bars represent the mean ± SEM of three independent experiments with three technical replicates. Immunoblots are a representative experiment of three independent experiments. Statistical significance is represented in the graphs as: * p<0.05, ** p<0.01, *** p<0.005, **** p<0.001.

Since MCF10A cells harbor wild-type p53 and p53 is known to be a phosphorylation target of LATS (*34, 35*) we asked whether p53 was involved in the repression of p63 after the inhibition of MST1/2. When parental MCF10A cells and two p53 CRISPR KO clones were treated with XMU-MP1, p63 was downregulated to the same extent in all three cell lines (Fig. 1E, F) indicating that the effect of MST1/2 inhibition on p63 expression is independent of p53.

### LATS1 and MOB1A regulate p63 expression

Since both MST1 and 2 are required for p63 expression in MCF10A cells, we evaluated whether their downstream protein kinases targets, LATS1 and 2, were also required. We first treated MCF10A with the LATS1/2 inhibitor Truli (*36*) for 24 h which led to a 91% downregulation of p63 mRNA and a 7-fold induction of CTGF mRNA (Fig. 2B). p63 reduction and CTGF induction were also seen at the protein level along with decreased phosphorylation of YAP, and upregulation the total levels of YAP and TAZ proteins (Fig. 2A). There was also a mobility shift of phospho-TAZ protein, suggesting a change in its modifications due to LATS inhibition.

**Fig. 2.**
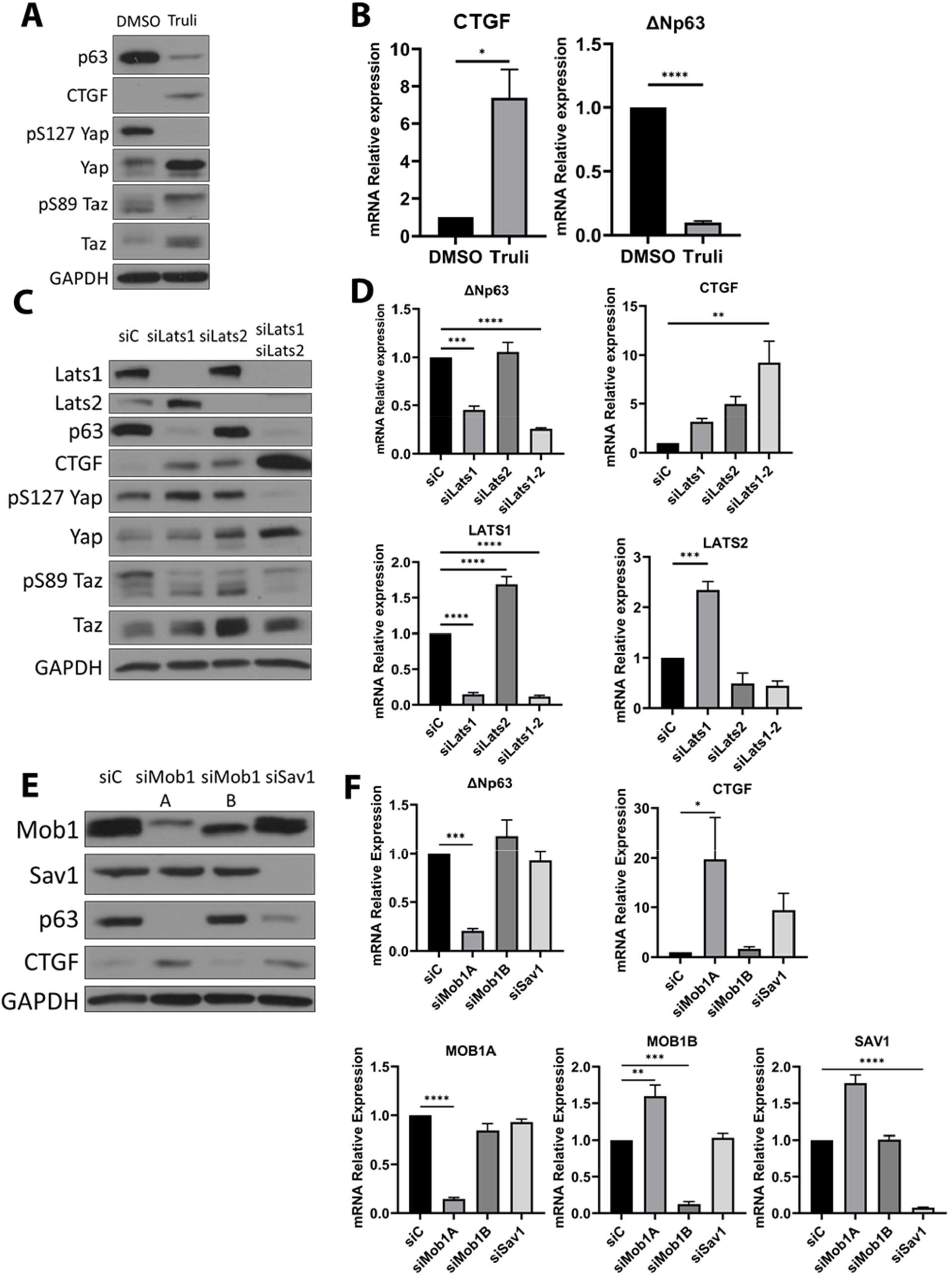
Lats1 and MOB1A regulate p63 expression. **A**. Immunoblot of MCF10A cells treated with DMSO or 5 µM of the LATS1/2 inhibitor Truli for 24 h. **B**. qRT-PCR of MCF10A cells treated as in A. **C**. Immunoblot of MCF10A cells transfected with Smartpool siRNA against LATS1, LATS2 or both, for 72h. **D**. qRT-PCR of MCF10A cells transfected as in C. **E**. Immunoblot of MCF10A cells transfected with Smartpool siRNA against MOB1A, MOB1B or SAV1 for 72h. **F**. qRT-PCR of MCF10A cells transfected as in E. For the immunoblots, one representative of three independent experiments is shown. For qRT-PCR, bars represent the mean ± SEM of three independent experiments with three technical replicates normalized to the siRNA control (siC) or DMSO treatment. Statistical analysis was Student t-tests in B and one-way ANOVA with Dunnett’s multiple comparisons test against the siC treatment were performed in D. and F. Statistical significance is represented in the graphs as: * p<0.05, ** p<0.01, *** p<0.005, **** p<0.001.

To confirm the inhibitor results, we ablated Lats1, Last2 and both Lats1 and 2 with siRNA to reduce their levels. Knockdown of LATS1, but not LATS2, downregulated p63 at the protein and mRNA levels (Fig. 2C, D) even though siRNA knockdown of both Lats1 and Lats2 were required for full activation of CTGF expression and decreased phospho-YAP and phospho-TAZ levels (Fig. 2C, D). These results showing that LATS1 is preferentially required for p63 repression, suggested that the repression of p63 might be through an independent process that is distinct from the activation of YAP and TAZ.

To further evaluate the requirement of the Hippo pathway for p63 expression, we tested cofactors of the MST and LATS kinases, SAV1 and MOB1A/B, respectively. When siRNAs directed against MOB1A, MOB1B or SAV1 were used, only MOB1A was required for p63 expression (Fig. 2E, 2F). We did not observe an effect of MOB1B siRNA on p63 or CTGF expression, but could not confirm this at the protein level due to lack of an antibody that can distinguish between A and B isoforms, although the siRNA for MOB1A abolished the lower band while the MOB1B siRNA abolishes the upper band (Fig. 2E). It is possible that MOB1B may simply be less abundant that MOB1A in MCF10A cells such that it is not required for Lats1/2 activity. The effects of SAV1 siRNA were relatively mild, despite general reports that SAV1 is required for MST activity (*28, 29*). While SAV1 siRNA increased CTGF protein and mRNA levels, there was no effect on p63 mRNA and a modest reduction of p63 protein (Fig. 2E, F). Further studies will be required to understand the role of SAV1 in this system. Nevertheless, the results with MOB1A further show the requirement of the Hippo pathway for p63 expression.

### MST and LATS1 regulate p63 expression in breast and squamous cancer cell lines

Since MCF10A is a non-transformed immortalized mammary epithelial cell line, it was of interest to evaluate whether p63 is regulated by the Hippo pathway in cancer cells as well. We selected previously described ΔNp63 positive breast cancer cell lines HCC1937 (triple negative) and HCC1954 (HER2 positive) (*37*) and the squamous carcinoma cell lines TE5, TE14 and Cal27 with high expression of ΔNp63 (see Supplemental Fig. 1 for relative expression of p63 mRNA). Treatment of HCC1937 and HCC1954 cells with the XMU-MP1 inhibitor of MST1/2 led to strong downregulation of ΔNp63 protein and mRNA (Fig. 3A, B). This correlated with markedly increased expression of the YAP/TAZ target gene CTGF (Fig. 3A, B) even though YAP and TAZ protein levels were only slightly increased by XMU-MP1 in both cell lines (Fig. 3A). In the squamous cancer cell lines, we also observed suppression of ΔNp63 mRNA and protein upon treatment with XMU-MP1, though the extent varied (Fig. 3C, D). CTGF mRNA and protein induction also varied, with the strongest activation in Cal27 cells (Fig. 3C, D). The reduction in phospho-YAP levels by XMU-MP1 treatment was also particularly strong in Cal27 cells (Fig. 3C). These results demonstrate that MST kinase activity is required to maintain p63 expression in multiple cancer cell lines.

**Fig. 3.**
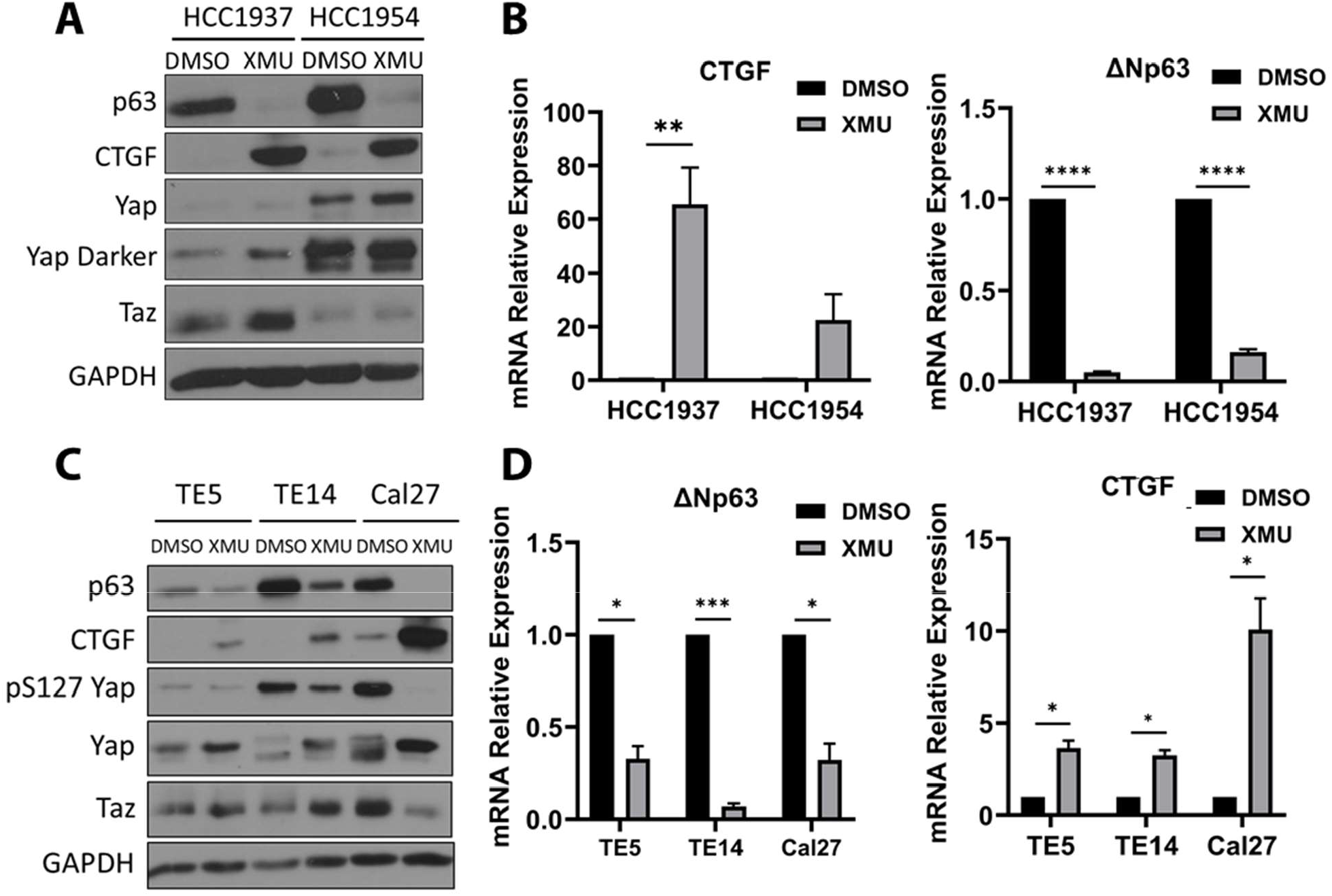
MST regulate p63 expression in Breast and Squamous carcinoma cell lines. **A**. Immunoblot of HCC1937 and HCC1954 breast carcinoma cell lines were treated with DMSO or 5 µM of XMU-MP1 for 24 h. **B**. RT-qPCR of HCC1937 and HCC1954 cells treated same as in A. **C**. Immunoblot of squamous carcinoma cells treated with XMU-MP1 for 24 h. **D**. RT-qPCR of squamous carcinoma cells treated with XMU-MP1 for 6 h. For the immunoblots, one representative of three independent experiments is shown. For qRT-PCR, bars represent the mean ± SEM of three independent experiments with three technical replicates normalized to the DMSO treatment. Statistical analysis was Student’s t-test. Statistical significance is represented in the graphs as: * p<0.05, ** p<0.01, *** p<0.005, **** p<0.001.

We next asked whether the LATS kinases were also involved in regulating p63 in cancer cells. We inhibited LATS1/2 activity with the inhibitor Truli and found that p63 was significantly downregulated in HCC1937 and HCC1954 cells (Supplemental Fig. 2A, B). A similar repression of p63 mRNA was found in the squamous cancer cells TE5, TE14 and Cal27 (Supplemental Fig. 2D) while p63 protein levels were clearly downregulated in the Cal27 cell line, with relatively little change in the other lines. The strongest repression of p63 protein was found in the Cal27 cell line despite being one of the cell lines with the highest relative expression of p63 mRNA (Supplemental Fig. 1). CTGF RNA was only increased about two fold in the breast cancer cell lines (Supplemental Fig. 2B). Increased CTGF protein levels were low in the HCC1937 cell line but clearly induced by Truli; CTGF protein levels were constitutively high in the HCC1954 cell line suggesting other mechanisms of increased expression (Supplemental Fig. 2B). In response to LATS inhibition, we observed a decrease in phospho-YAP and a mild increase of total levels of YAP in both cell lines; TAZ levels increased mostly in HCC1937 cells (Supplemental Fig. 2A). While phospho-YAP inhibition was seen in each of the squamous cell lines, the increase in total YAP levels was seen the clearest in Cal27 cells (Supplemental Fig. 2C). We can conclude from these results that in breast and squamous carcinoma cells MST and LATS activity are required for the expression of p63. However, the effects of the inhibition of these kinases on the levels of YAP, TAZ, and CTGF varied depending on the cell line and suggested to us that the p63 repression and CTGF induction pathways are different. In order to confirm that LATS1 activity but not LATS2 is required for p63 expression in cancer cells, as we showed for MCF10A (Fig. 2C, D). We knockdown LATS1, LATS2 and both kinases in the SCC cell line Cal27 with Smartpool siRNA for 72 h (Supplemental Fig. 2E, F) and as observed in MCF10A, ablation of LATS1 but not LATS2 significantly repressed p63, while the activation of YAP/TAZ and CTGF induction increased significantly with the knocldown of either LATS1 or 2 (Supplemental Fig. 2F).

### Repression of p63 after MST1/2 inhibition is YAP/TAZ independent

YAP and TAZ are inhibited by LATS phosphorylation (*19, 20, 23*) which led us to ask whether they are also involved in the repression of p63 caused by MST inhibition. To address this, we used two approaches. We inhibited YAP transcriptional activity with the drug Verteporfin that impairs YAP interaction with TEAD transcription factors (*38*) and we suppressed YAP and TAZ levels with two different siRNAs. In both cases we observed that neither the inhibitor (Fig. 4A, B) nor the siRNA (Fig. 4C, D) had any effect on the repression of p63 induced by XMU-MP1 inhibition of MST; p63 was still suppressed at the mRNA level despite the lack of YAP/TAZ protein and activity. We confirmed that YAP/TAZ inhibition was effective by observing a significant drop in the induction of CTGF mRNA and protein (Fig. 4A-D). These results indicate that YAP/TAZ activity is not required for the repression of p63 upon MST inhibition and implicate the involvement of other factors downstream of LATS1 that are not part of the canonical pathway.

**Fig. 4.**
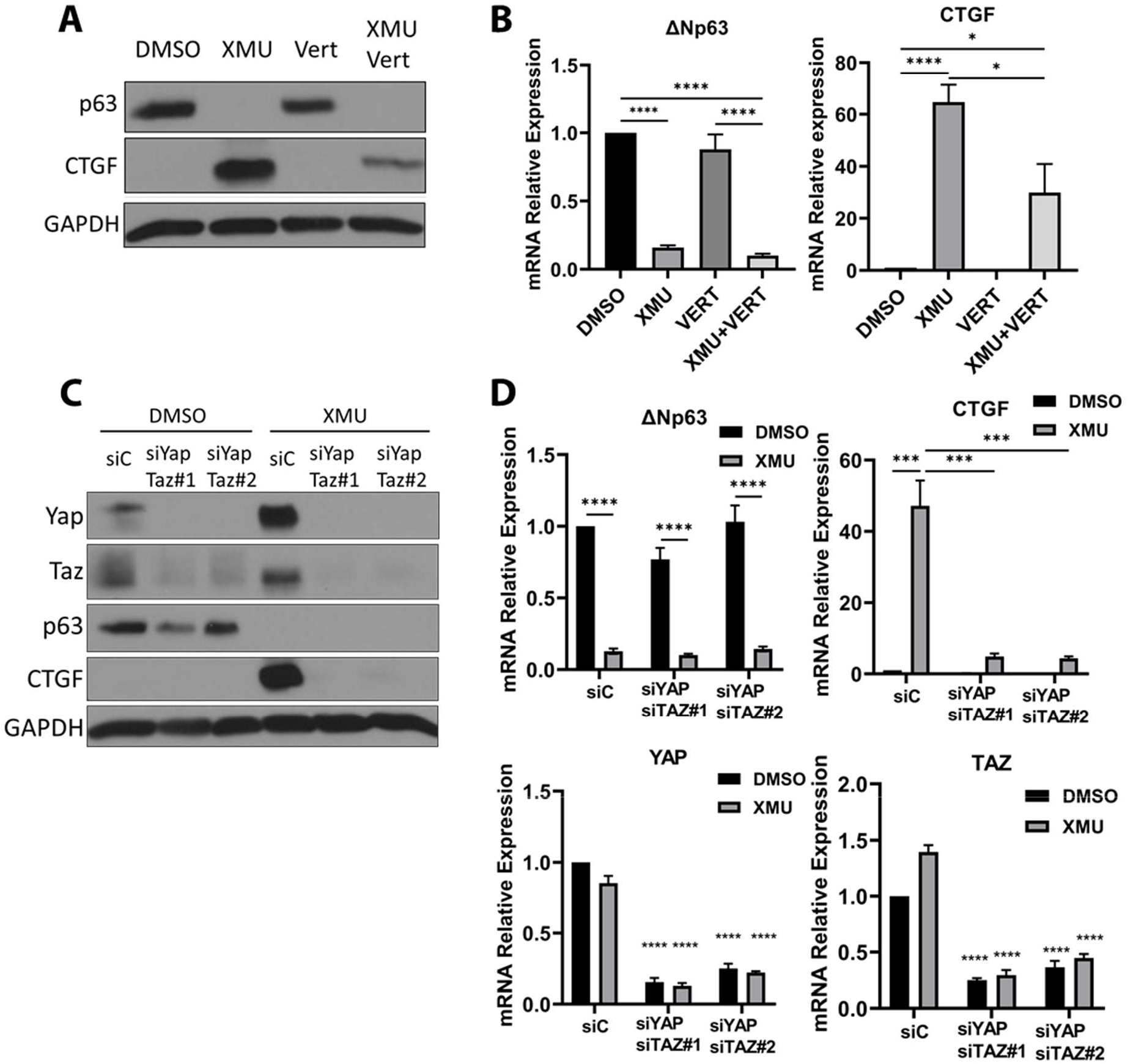
Repression of p63 after MST1/2 inhibition is Yap/Taz independent. **A**. Immunoblot of MCF10A cells treated with DMSO or 5 µM of XMU-MP1, 4 µM of the YAP/TAZ inhibitor Verteporfin, or a combination of both inhibitors for 24 h. **B**. RT-qPCR of MCF10A cells treated as shown in A. Bars represent the mean ± SEM of four independent experiments with three technical replicates, normalized to DMSO **C**. Immunoblot of MCF10A cells transfected with the combination two different siRNAs against YAP and TAZ for 24 h, then treated with 5 µM of XMU-MP1 for 24 h. **D**. RT-qPCR of MCF10A cells transfected with 2 different siRNAs against YAP and TAZ for 24 h, then treated with 5 µM of XMU-MP1 for 24 h. Bars represent the mean ± SEM of three independent experiments with three technical replicates, normalized to DMSO treated siControl (siC). Statistical analysis was ANOVA with Dunnett’s test for multiple comparisons against the siC treated with DMSO. For the immunoblots one representative experiment of three independent experiments is shown. Statistical significance is represented in the graphs as: * p<0.05, ** p<0.01, *** p<0.005, **** p<0.001.

### p63 inhibits YAP activity

Since regulating Hippo pathway had such a strong effect on p63 expression, we hypothesized that there might be reciprocity such that p63 might impose feedback inhibition on the Hippo pathway. When we tested this by treating MCF10A cells with two different siRNAs to p63, and found a particularly strong increase in YAP protein levels and increased mRNA and protein levels of the YAP/TAZ target gene CTGF (Fig. 5A, B). p63 siRNAs increased the total levels of both the YAP and TAZ proteins, although their respective mRNA levels were not changed, suggesting that the regulation is post-transcriptional (Fig. 5A, B). As there was only a slight increase in YAP and TAZ phosphorylation (Fig. 5A), the relative amount of phosphorylated YAP to total YAP went down upon p63 ablation, as shown in the top right panel of Fig. 5A. The increase in TAZ upon p63 knockdown did not change the relative phosphorylation significantly (as shown in Fig. 5A. bottom right panel). We also tested whether p63 loss could also increase YAP/TAZ activity after XMU-MP1 treatment by measuring induction of the CTGF gene and found increased induction of CTGF when p63 levels were suppressed with siRNAs (Fig. 5C). These results show that p63 serves to restrict YAP levels and hence dampen their transcriptional activity in MCF10A cells.

**Fig. 5.**
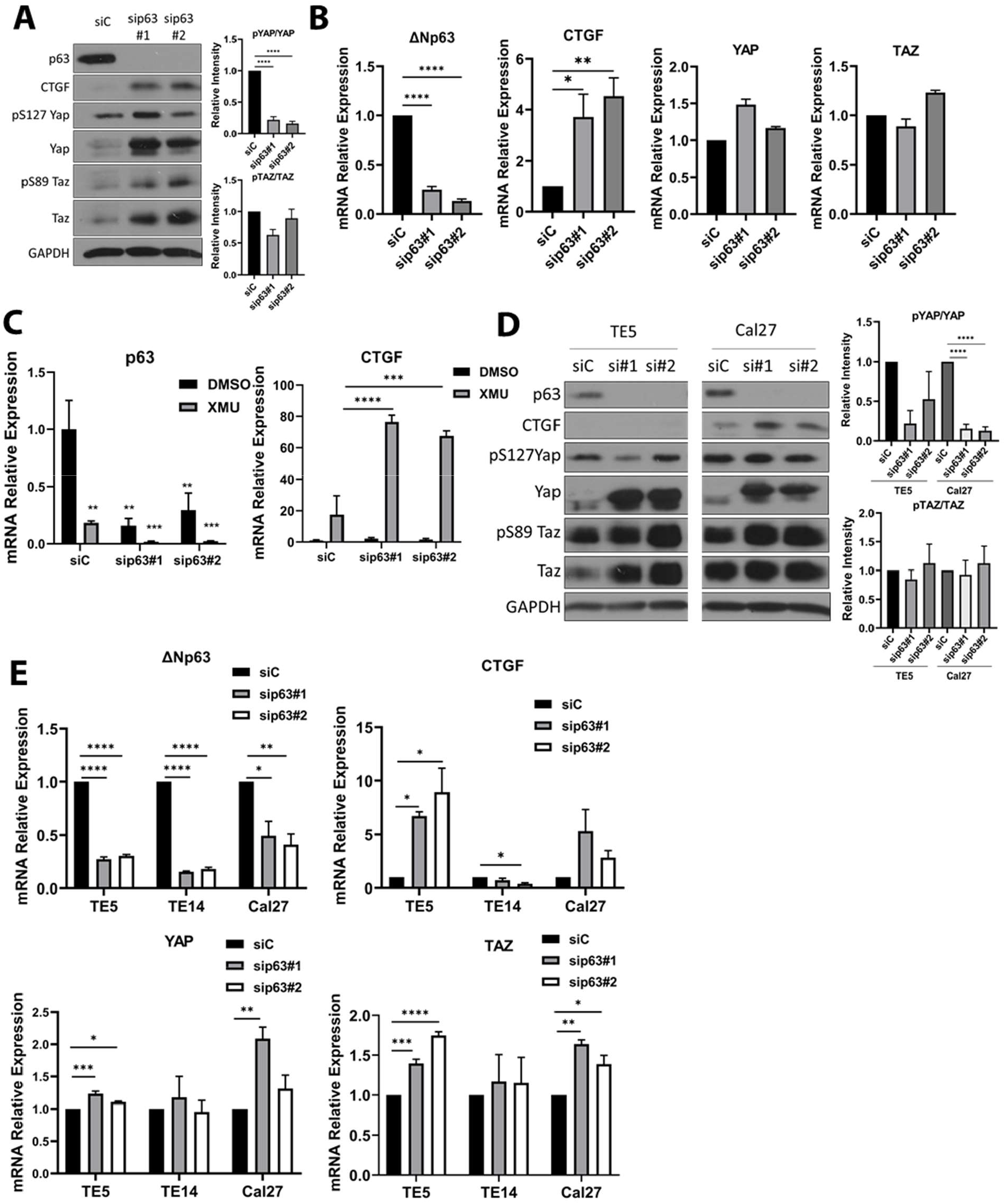
p63 inhibits YAP activity. **A**. Left, Immunoblot of MCF10A cells transfected with siControl or two siRNAs against p63 for 72 h. Top Right, Bars represent mean ± SEM of densitometry quantification of the relative intensity of phospho Ser^127^ YAP over the relative intensity of the total YAP, normalized to the siC treatment. Bottom Right, Bars represent mean ± SEM of densitometry densitometry quantification of the relative intensity of phospho Ser^89^ TAZ over the relative intensity of the total TAZ, normalized to the siC treatment. **B**. RT-qPCR of same cells shown in A. right **C**. RT-qPCR of MCF10A cells transfected with siC or two siRNAs against p63 for 72 h then treated for 6 h with DMSO or 5 µM XMU. **D**. Left, Immunoblot of TE5, TE14 and Cal27 cells transfected with siC or two siRNAs against p63 for 72 h. Top Right, Bars represent mean ± SEM of densitometry quantification of the relative intensity of phospho Ser^127^ YAP over the relative intensity of the total YAP, normalized to the siC treatment. Bottom Right, Bars represent mean ± SEM of densitometry quantification of the relative intensity of phospho Ser^89^ TAZ over the relative intensity of the total TAZ, normalized to the siC treatment. **E**. qRT-PCR of TE5, TE14 and Cal27 cells transfected with siC or two siRNAs against p63 for 72 h. For the immunoblots one representative experiment of three is shown. For RT-qPCR the bars represent the mean ± SEM of three independent experiments with three technical replicates, normalized to siC. For the CTGF RT-qPCR experiment shown in B, four independent experiments were performed. Statistical analysis was One-Way ANOVA with Dunnett’s test for multiple comparisons against the siC treatment or the DMSO siC treatment. Statistical significance is represented in the graphs as: * p<0.05, ** p<0.01, *** p<0.005, **** p<0.001.

We also found that p63 siRNAs increased YAP protein levels in each of the three SCC cell lines used (Fig. 5D and Supplemental Fig. 3), while YAP mRNA was unaffected (Fig. 5E and Supplemental Fig. 3), suggesting the broad presence of this pathway whereby p63 loss stabilizes YAP protein levels. However, the effects on on CTGF and TAZ protein levels varied among the three SCC cell lines. In TE5 cells, p63 knockdown increased YAP and TAZ protein levels and CTGF mRNA, yet CTGF protein was undetectable (Fig. 5D and E). In TE14 cells, knockdown increased YAP levels, but there was little effect on TAZ levels or CTGF mRNA and CTGF protein was undetectable (Fig. 5E and Supplemental Fig. 3). Finally in Cal27 cells, p63 knockdown increase YAP levels and CTGF mRNA, but had little effect on TAZ protein levels (Fig. 5D and E). Despite the variability of the results in these cell lines, our data show clearly that p63 inhibits YAP levels and activity in both MCF10A and SCC cell lines. This suggests that YAP activity to increase CTGF expression is regulated by the Hippo pathway both by inhibition of LATS phosphorylation and by p63 activity.

### Genes repressed by the non-canonical Hippo pathway

We next asked whether the non-canonical pathway is more generally used and whether other genes were repressed by the Hippo pathway independent of YAP and TAZ. We performed RNAseq on MCF10A cells that were treated with XMU-MP1 with control or YAP/TAZ siRNAs. We expected that genes regulated similar to p63 would be repressed by XMU-MP1 treatment both with and without YAP/TAZ siRNAs. We found 333 genes that were repressed more than 3 fold by XMU-MP1 (Supplemental Table 2A). Of these, 162 were also repressed greater than 3 fold upon YAP/TAZ knockdown, suggesting that their repression is independent of YAP and TAZ (Supplemental Table 2B). The p63 gene was found in this group of genes as expected.

We identified 184 genes induced greater than 3 fold (albeit with a relaxed p value of 0.2) (Supplemental Table 2C). Of these genes, 82 were induced less than half in siYAP/TAZ cells, suggesting dependence on YAP and TAZ (Supplemental Table 2D). These include the well known Hippo targets CTGF and CYR61 (a.k.a. CCN2 and CCN1). We sought to confirm the repression of some of the genes identified by RNAseq by RT-qPCR. We picked 8 genes with varied repression levels and found that 7 of the 8 were strongly repressed by XMU-MP1 treatment (Fig. 6A). If these genes are regulated similar to p63 by the Hippo pathway, we would expect that they are also regulated by LATS1/2. We tested this by treating cells with the LATS1/2 inhibitor TRULI. We found that 2 (ELF3 and CYPB1) out of the 7 genes tested repressed by XMU-MP1, were also repressed by TRULI. One gene, PLAU, was modestly reduced, while the others were unaffected or induced (IL1A) by TRULI (Fig. 6B). However, these results show that there is a set of genes including TP63, ELF3 and CYP1B1 that are dependent on the Hippo pathway kinases and independent of YAP and TAZ (Fig. 6C). Using gene ontology, we did not find a significant categorization of the XMU-MP1 regulated genes. Nevertheless, ELF3 is notable as it is an ets-related transcription factor expressed specifically in epithelial cells similar to p63 (*39*) CYP1B1 is a cytochrome 450 related gene involved in detoxification of chemicals, but it is not clear why it would be regulated by the Hippo kinases (*40*).

**Fig. 6.**
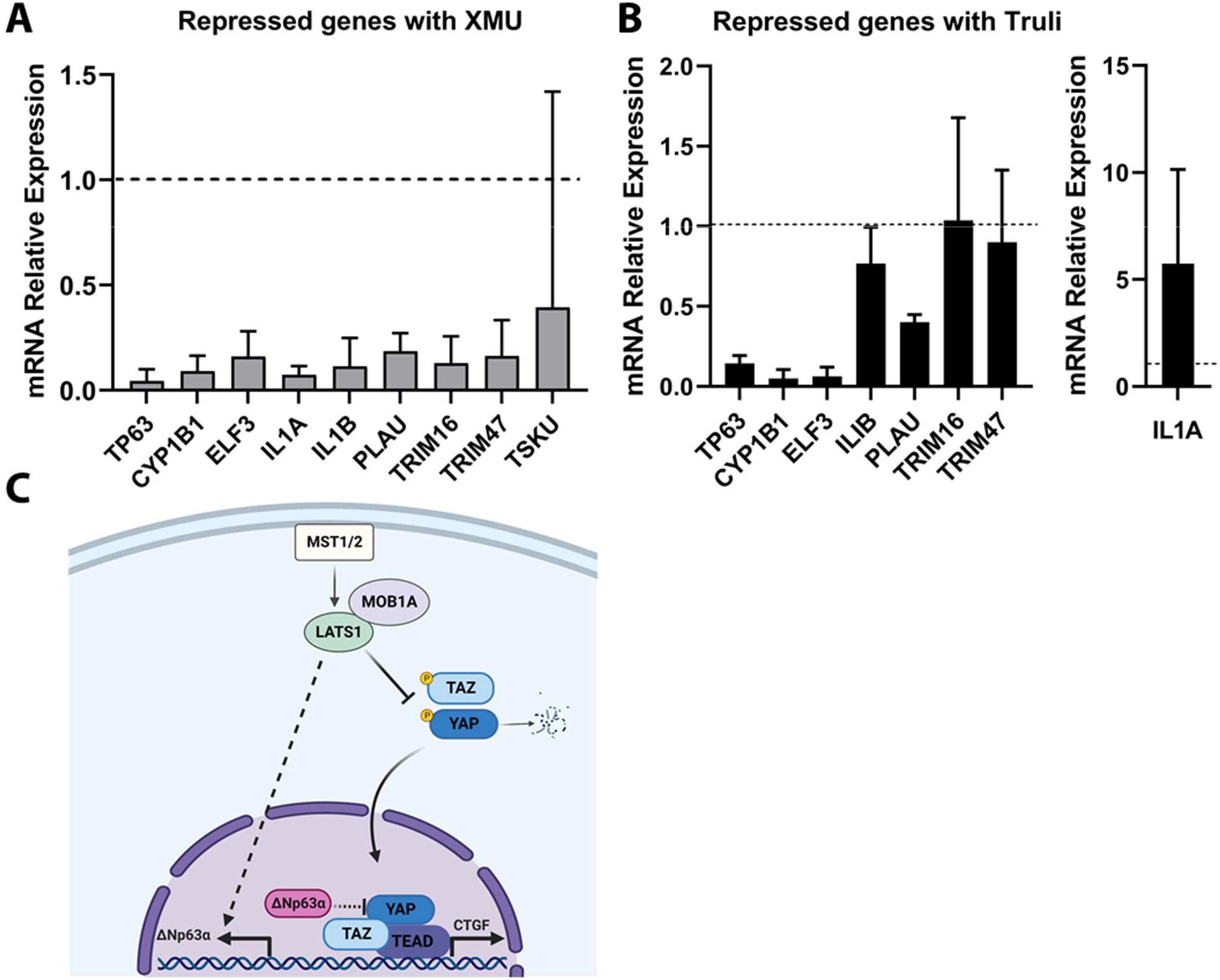
RNAseq validation of Repressed genes by XMU-MP1 and Truli treatment. **A**. RT-qPCR of MCF10A cells treated with 5 µM of XMU-MP1 for 6 h. Bars represent the mean relative expression ± 95% CI of four independent experiments. The horizontal line represents the relative expression of 1 in control DMSO treated cells. **B**. RT-qPCR of MCF10A cells treated with 10 µM of Truli for 24 h. Bars represent the mean relative expresssion ± 95% CI of three independent experiments. Horizontal lines represent the relative expression of 1 in control DMSO treated cells. **C**. Model of a non-canonical Hippo pathway that regulates the expression of p63 and CTGF separately in mammary and cancer cells. In addition, the repression of YAP levels by p63 is shown.

## Discussion

Here we describe that p63 expression is regulated by the Hippo pathway in different types of cells including non-transformed mammary MCF10A cells, breast cancer, and tongue and esophageal squamous carcinoma cells. Further our data indicate that the Hippo pathway regulates the expression of ΔNp63 in a non-canonical manner. While the MST1/2, LATS1 and MOB1A factors are required, regulation of p63 is independent of YAP and TAZ (Fig. 6C). In addition, we found that p63 feeds back into the pathway, causing the inhibition of YAP/TAZ, since p63 loss increased YAP/TAZ protein levels and the induction of their target gene CTGF.

Surprisingly, in our experimental setting, we found that LATS1 but not LATS2 was preferentially required for the expression of p63, even though CTGF induction required the inhibition of both LATS1 and LATS2. LATS1-specific effects have been seen before: LATS1 inhibited autophagy in a kinase-independent manner while LATS2 had no effect (*41*). Moreover, in an affinity proteomics study done by immunoprecipitation with overexpressed MOB1A/B proteins, MOB1A bound LATS1 proteins more abundantly compared to LATS2, suggesting a stronger interaction with LATS1 (*42*). This could explain the difference between LATS1 and LATS2 in their regulation of p63 expression. The preferential inhibition of LATS1 may provide a means to repress p63 expression without the potential side effects or toxicity of inhibiting both LATS1 and LATS2.

MOB1A and MOB1B are scaffold proteins that activate LATS kinases. Phosphorylation of MOB1 by MST1/2 allows them to bind to LATS1/2 (*27*); in turn LATS1/2 can then be phosphorylated and activated by MST1/2 (*26, 27, 43*). We found that MOB1A proteins were required for maintenance of p63 expression while there was little effect of MOB1B depletion, despite the fact that their protein products are 95% identical (*44*). It is possible, however, that this is due to the higher expression of MOB1A compared to MOB1B (Fig. 2E and our previous RNAseq data (*10*)). Alternatively, MOB1A binds LATS1 more tightly compared to MOB1B (*42*). Since p63 repression and CTGF induction were similarly sensitive to MOB1A depletion, this suggests that a common pathway involving MOB1A and LATS1 is involved. The unique requirement of LATS1 in this complex, as opposed to LATS2, may be due to preferential binding of substrates for LATS1.

The protein SAV1 binds to and is required for MST1/2 recruitment to the plasma membrane (*28, 45*) and promotes the auto-phosphorylation and activation of MST1/2 by preventing its interaction with the phosphatase STRIPAK (*29*). In contrast to expectations, SAV1 depletion did not inhibit MST activity enough to cause the downregulation of p63 mRNA, while it only increased CTGF expression moderately (Fig. 2F). Although the extent of depletion of SAV1 protein and mRNA by siRNAs appears strong (Fig. 2E, F), it is possible that residual SAV1 is sufficient to maintain p63 levels. Alternatively, MST1/2 may be able to function independently of SAV1 in a pathway to maintain p63 expression. Surprisingly, SAV1 loss decreased p63 protein levels, but not p63 mRNA (Fig. 2E, 2F) suggesting post-transcriptional regulation. p63 protein stability has been found to be regulated by the E3 ubiquitin ligases Pirh2, WWP1, and ITCH among others (*46*–*48*). Interestingly, ITCH is also an E3 ligase for the Hippo kinase LATS1 (*49*). Further studies will be required to determine whether LATS1 and SAV1 regulate p63 protein stability through any of these E3 ligases.

The most striking deviation from the canonical Hippo pathway that we discovered was that YAP and TAZ are not required to repress p63 levels in multiple cell lines. While many transcription factors have been reported to regulate ΔNp63 expression (*50*) it’s unclear which, if any such factors, may be acting here. We tested FOXO3 as a potential regulator (*11, 51*), but found no role (Supplemental Fig. 4). Further studies will be required to identify the steps in the pathway between LATS1 and ΔNp63. In contrast to our results, overexpression of TAZ was shown to repress ΔNp63 expression (*52*). This difference may be due to differential effects exerted by the overexpression of TAZ alone compared to the inhibition of endogenous MST1/2 that we reported here.

Other non-canonical (YAP/TAZ-independent) pathway functions for LATS kinases have been found. LATS1 can target Estrogen Receptor α for ubiquitin-mediated degradation (*53*) and LATS2 binds to and inhibits MDM2 to prevent p53 degradation (*34*). In addition, LATS2 but not LATS1 inhibits the maturation and subsequent nuclear translocation of the mevalonate pathway transcription factor SREBP2 (*54*). Together with our results, this suggests that there are additional targets of the LATS kinases besides YAP and TAZ that can mediate effects of the Hippo pathway.

This YAP/TAZ-independent Hippo pathway is also supported by our RNAseq data where we found that a set of genes is repressed upon inhibition of MST1/2 (Fig 6A). Similarly, several of those genes were also repressed by LATS1/2 inhibition (Fig 6B) suggesting that a similar downstream factor is responsible for the repression of this subset of genes (p63, ELF3, CYP1B1 and PLAU). However, four of the genes repressed by XMU treatment were not repressed by TRULI (such as IL1A, IL1B, TRIM16 and TRIM47). This suggests that there is a pathway dependent upon MST1/2, but independent of both LATS1/2 and YAP/TAZ. It will be important to determine more factors involved in each of these alternative Hippo pathways and their physiological effects.

In conclusion, we found that it is possible to downregulate the levels of p63 in breast and squamous carcinoma cells as well as immortalized mammary cells by inhibiton of the Hippo pathway kinases. This defines an alternative Hippo pathway, independent of YAP/TAZ, that also regulates a set of other genes. We propose that these kinases and associated factors involved in p63 regulation could be useful targets for further development of therapies for cancers that show a strong dependency on p63 for growth. It will be important to take into account that their targeting has the potential to increase YAP/TAZ levels and their induction of cellular growth. This complication could be accommodated by inhibition of YAP/TAZ with an existing drug like Verteporfin. Further understanding of this pathway to identify factors that are specific to Hippo regulation of p63 levels will also be useful for finding specific inhibitors of this wing of the Hippo pathway.

## Materials and Methods

### Cell culture

The MCF10A immortalized non-transformed mammary epithelial cell line (a gift obtained from David Weber, University of Maryland School of Medicine) were grown in DMEM/F12 (Thermo Fisher Scientific, 11320033) with 5% horse serum (Thermo Fisher Scientific, 16050122), penicillin-streptomycin solution (Thermo Fisher Scientific, 15140122), 10 µg/mL insulin (Sigma-Aldrich, 91077C), 0.5 µg/mL hydrocortisone (Sigma-Aldrich, H0888) and 20 ng/mL Epidermal Growth Factor (PeproTech, AF-100-15). MCF10A CRISPR p53 Knock out (KO) clones were generated using CRISPR/Cas9 genome editing as previously described (*55*). Esophageal squamous cell carcinoma cell lines TE5 and TE14 were a gift from Anil Rustgi, Herbert Irving Comprehensive Cancer Center, Columbia University Irving Medical Center and were grown in RPMI 1640 (Thermo Fisher Scientific, 11875119) supplemented with 10% fetal bovine serum (Gemini Bio-Products, 900108H) and penicillin-streptomycin solution. The Cal27 tongue squamous cell carcinoma cell line was a gift from Alison Taylor (Columbia University Irving Medical Center) and was maintained in DMEM (Thermo Fisher Scientific, 12100061) supplemented with 10% fetal bovine serum and penicillin streptomycin solution. Breast cancer cell lines HCC1937 (Triple negative, primary ductal carcinoma) and HCC1954 (HER2 positive, ductal carcinoma) were obtained from the High-Throughput Screening facility of the Columbia University Genome Center. These cell lines were grown in RPMI 1640 with 10% fetal bovine serum and penicillin-streptomycin solution. Cells were detached with trypsin-EDTA (0.05%) (Thermo Fisher Scientific, 15400054) and counted before seeding for experiments using trypan blue in a Countess 3 Automated Cell Counter (Thermo Fisher Scientific). All cell lines were grown at 37°C with 5% CO_2._

### siRNA transfection

For siRNA transfections, cells (1.5×10^5^) were plated in 6 well plates and 24 h later each siRNA (17 nM) was transfected with Lipofectamine™ RNAiMAX (Thermo Fisher Scientific, 13778150) following manufacturer’s instructions. Cells were harvested 72 h post transfection. The following siRNAs were used siC (Silencer™ Select Negative Control No. 1, Thermo Fisher Scientific, 4390843), sip63#1 (Thermo Fisher Scientific, 4392420, ID s16411), sip63#2 (Thermo Fisher Scientific, 4392420, ID s531582), FoxO3#1 (Thermo Fisher Scientific, 4427037, ID s5260), and FoxO3#2 (Thermo Fisher Scientific, 4427037, ID s5261), siYap#1 (Thermo Fisher Scientific, 4427037, ID s20366), siYap#2 (Thermo Fisher Scientific, ID s20367), siTaz#1 (Thermo Fisher Scientific, 4392420, ID s24787), and siTaz#2 (Thermo Fisher Scientific, 4392420, ID s24788). The following Smartpool siRNAs were used: siRNA negative control (siC) (Dharmacon, D-001810-10-05), siMst1 (Dharmacon, L-004157-00-0005), siMst2 (Dharmacon, L-004874-00-0005), siLats1 (Dharmacon, L-004632-00-0005), siLats2 (Dharmacon, L-003865-00-0005), siMob1a (Dharmacon, L-021097-00-0005), siMob1b (Dharmacon, L-018359-01-0005) and siSav1 (Dharmacon, L-013070-01-0005). To decrease levels of YAP and TAZ proteins, MCF10A (3×10 cells/well) were seeded in 6 wells, 24 h later the siRNAs (11 nM) were transfected into wells and 24 h post transfection fresh media with or without drug (e.g. XMU-MP1) was added and cells were harvested 24 h later.

### Inhibitors

XMU-MP1 (Selleck Chemicals, S8334) is an inhibitor of MST1/2 (*33*). Truli (Gift from Dr. J. Hudspeth and also obtained from CSNPharm, CSN26140) is an inhibitor of LATS1/2 (*36*). Verteporfin (Selleck Chemicals, S1786) is an inhibitor of YAP binding to the TEAD transcription factors (*38*).

### Immunoblot

Total protein lysates were obtained by washing the cells with ice cold phosphate buffered saline (PBS), and cells were lysed with RIPA buffer (50 mM Tris-HCl pH 8.0, 150 mM NaCl, 0.5% Sodium Deoxycholate, 0.1% Sodium Dodecyl Sulfate, and 1% NP40) supplemented with fresh protease inhibitors (3 µg/mL Leupeptin, 1 µg/mL macroglobulin, 1 µM benzamidine, 0.5 µM PMSF) and phosphatase Inhibitors (EMD Millipore, 524625). Cell lysates were incubated 10 minutes on ice and centrifuged at 13000 rpm for 10 minutes. The protein concentrations of sample supernatants were measured with the Bio-Rad protein assay dye reagent (Bio-Rad, 5000006). Equivalent amounts of protein samples were mixed with Laemmli buffer (100 mM Tris-HCl pH 6.8, 2% SDS, 5% 2-mercaptoethanol, 5% glycerol) and incubated at 95°C for 5 min. Lysates (35 µg protein) were run on 10% SDS-polyacrylamide gels and transferred to a nitrocellulose membranes (Bio-Rad, 1620115). After blocking for 10 min with 5% nonfat dried milk in PBS with 0.05% Tween-20 (Sigma-Aldrich, P1379) (PBST), the membrane was incubated with primary antibodies diluted in 0.1% milk in PBST at 4°C overnight. The following primary antibodies from Cell Signaling Technologies were used: GAPDH (1:10000, 5174), MST1 (1:500, 3682), MST2 (1:1000, 3952), LATS2 (1:500, 5888), pS127-YAP (1:8000, 13008), YAP (1:4000, 14074), pS89-TAZ (1:500, 59971), TAZ (1:500, 71192), CTGF (1:300, 86641), MOB1A/B (1:500, 13730), SAV1 (1:1000, 13301), and FOXO3 (1: 1000, 2497). In addition, primary antibodies to LATS1 (1:1000, Millipore, MABS1823) and p63 (1:500, Biocare Medical, CM163A) were used. The secondary antibodies, peroxidase-conjugated Anti-Rabbit IgG (Sigma-Aldrich, A6154) and anti-mouse IgG (Sigma-Aldrich, A4416), were incubated for 1 h at room temperature at 1:5000 dilution in 0.5% milk PBST for 1 h. Membranes were imaged with Pierce™ ECL Western Blotting Substrate (Thermo Fisher Scientific, 322106) or Immobilon Western Chemiluminescent HRP Substrate (Millipore, WBKLS0500). Quantification of changes in phospho YAP and phosphotaz TAZ were calculated as the relative intensity by the densitometry of each protein band, divided by the intensity of GAPDH. The relative intensity of the protein in the sip63 samples was normalized to the relative intensity in the siRNA control sample. The mean of three independent immunoblot experiments was calcuated. The densitometry was analyzed with an ImageJ Gel analyzer plug in.

### Quantitative RT-PCR

Total RNA (500 ng) isolated with the RNAeasy mini kit (Qiagen, 74106), was converted to cDNA with the QuantiTect Reverse Transcription Kit (Qiagen, 205313) following manufacturer’s instructions. Real-time PCR was performed in the StepOne Plus thermal cycler (Applied Biosystems) with Applied Biosystems Power SYBR Green PCR Master Mix (Thermo Fisher Scientific, 4368708) and primers (0.2 µM) following the program: 95°C for 10 minutes followed by 40 cycles of 95°C for 15 seconds, 60°C for one minute. Melting curves were performed between 60°C -95°C with 1°C/sec. The relative expression of each gene was calculated according to the ΔΔCt method expression using the Ct values of the housekeeping gene RPL32 (or ribosomal 18S RNA for Figs. 3D, 5C and 6) and normalized to the siRNA control or DMSO treated condition. For the list of primers used see Supplemental Table 1.

### RNAseq analysis and validation by RT-qPCR

MCF10A cells (3×10^5^ cells per well) were seeded in 6 well plates. Next day cells were transfected with 11 nM of two different siRNAs sets against YAP and TAZ or 22 nM of control siRNA; 48 h post transfection fresh media with DMSO or 5 µM of XMU-MP1 was added and cells were harvested 6 h later. Total RNA was isolated using TRIzol reagent (Thermofisher, 15596026) following the manufacturer’s instructions. Samples from three independent experiments were sent for RNA sequencing and analysis by LC Sciences as described by the company (https://lcsciences.com/services/rna-sequencing-services/transcriptome-sequencing/). Select repressed genes that were differentially expressed and considered statistically significant (p<0.05) were validated by RT-qPCR. Relative expression was normalized to 18S gene. In addition, the same set of repressed genes was validated in MCF10A cells treated with DMSO or 10 µM of the LATS inhibitor Truli for 24 h.

### Statistical analysis

Statistical analysis was performed using GraphPad Prism Software version 9.3.1 for Windows (GraphPad Software, San Diego, California USA, www.graphpad.com). For simple comparisons between two treatments Student’s t-test was used. For analysis between three or more treatments One-way ANOVA with Dunnett’s multiple comparisons test was performed. Statistical significance is represented in the graphs as: * p<0.05, ** p<0.01, *** p<0.005, **** p<0.001.

## Supplementary Materials

Supplemental Fig. 1. Comparison of p63 mRNA levels across cell lines.

Supplemental Fig. 2. LATS regulate p63 expression in breast and squamous cancer cell lines. Supplemental Fig. 3. p63 inhibits YAP activity.

Supplemental Fig. 4. FoxO3 is not required for ΔNp63 expression in MCF10A cells.

Supplemental Table 2A. Genes significantly repressed by XMU.

Supplemental Table 2B. Genes significantly repressed by XMU-MP1 independent of YAP/TAZ. Supplemental Table 2C. Genes induced by XMU-MP1.

Supplemental Table 2D. YAP/TAZ-dependent genes induced by XMU-MP1.

## Acknowledgements

We thank Ella Freulich for her help in tissue culture and technical assistance and Joshua Choe for his helpful advice and suggestions. We thank the current and former members of the Prives Lab, for their advice and useful discussions. We thank Dr. Anil Rustgi, Dr. Alison Taylor and Dr. David Weber for providing the cell lines we used in this study and Dr. James Hudspeth for providing us with the LATS small molecule inhibitor Truli.

## Author contributions

A.M.L.C., C.P. and R.P. conceived the project and designed the experiments; A.M.L.C., R.P., H.G., A.M.C., N.O., D.T. and C.K. performed the experiments, A.M.L.C., H.G., N.O., R.P., A.M.C., C.K., C.P. and R.P. analyzed the data, and A.M.L.C., C.P., and R.P. wrote the manuscript.

## Data and Materials availability

The RNA-seq data will be available in GEO Datasets with accession number to be determined.

## References

1. A. Yang, R. Schweitzer, D. Sun, M. Kaghad, N. Walker, R. T. Bronson, C. Tabin, A. Sharpe, D. Caput, C. Crum, F. McKeon, p63 is essential for regenerative proliferation in limb, craniofacial and epithelial development. Nature. 398, 714–718 (1999).

2. A. A. Mills, B. Zheng, X.-J. Wang, H. Vogel, D. R. Roop, A. Bradley, p63 is a p53 homologue required for limb and epidermal morphogenesis. Nature. 398, 708–713 (1999).

3. A. Yang, M. Kaghad, Y. Wang, E. Gillett, M. D. Fleming, V. Dötsch, N. C. Andrews, D. Caput, F. McKeon, p63, a p53 Homolog at 3q27–29, Encodes Multiple Products with Transactivating, Death-Inducing, and Dominant-Negative Activities. Molecular Cell. 2, 305–316 (1998).

4. C. J. D. Como, M. J. Urist, I. Babayan, M. Drobnjak, C. V. Hedvat, J. Teruya-Feldstein, K. Pohar, A. Hoos, C. Cordon-Cardo, p63 Expression Profiles in Human Normal and Tumor Tissues. Clin Cancer Res. 8, 494–501 (2002).

5. C. B. Marshall, J. S. Beeler, B. D. Lehmann, P. Gonzalez-Ericsson, V. Sanchez, M. E. Sanders, K. L. Boyd, J. A. Pietenpol, Tissue-specific expression of p73 and p63 isoforms in human tissues. Cell Death Dis. 12, 745 (2021).

6. D. K. Carroll, J. S. Carroll, C.-O. Leong, F. Cheng, M. Brown, Alea. A. Mills, J. S. Brugge, L. W. Ellisen, p63 regulates an adhesion programme and cell survival in epithelial cells. Nature Cell Biology. 8, 551–561 (2006).

7. L. Boldrup, P. Coates, X. Gu, K. Nylander, ΔNp63 isoforms regulate CD44 and keratins 4, 6, 14 and 19 in squamous cell carcinoma of head and neck. The Journal of Pathology. 213, 384–391 (2007).

8. K. Hibi, B. Trink, M. Patturajan, W. H. Westra, O. L. Caballero, D. E. Hill, E. A. Ratovitski, J. Jen, D. Sidransky, AIS is an oncogene amplified in squamous cell carcinoma. Proceedings of the National Academy of Sciences. 97, 5462–5467 (2000).

9. J. D. Campbell, C. Yau, R. Bowlby, Y. Liu, K. Brennan, H. Fan, A. M. Taylor, C. Wang, V. Walter, R. Akbani, L. A. Byers, C. J. Creighton, C. Coarfa, J. Shih, A. D. Cherniack, O. Gevaert, M. Prunello, H. Shen, P. Anur, J. Chen, H. Cheng, D. N. Hayes, S. Bullman, C. S. Pedamallu, A. I. Ojesina, S. Sadeghi, K. L. Mungall, A. G. Robertson, C. Benz, A. Schultz, R. S. Kanchi, C. M. Gay, A. Hegde, L. Diao, J. Wang, W. Ma, P. Sumazin, H.-S. Chiu, T.-W. Chen, P. Gunaratne, L. Donehower, J. S. Rader, R. Zuna, H. Al-Ahmadie, A. J. Lazar, E. R. Flores, K. Y. Tsai, J. H. Zhou, A. K. Rustgi, E. Drill, R. Shen, C. K. Wong, S. J. Caesar-Johnson, J. A. Demchok, I. Felau, M. Kasapi, M. L. Ferguson, C. M. Hutter, H. J. Sofia, R. Tarnuzzer, Z. Wang, L. Yang, J. C. Zenklusen, J. (Julia) Zhang, S. Chudamani, J. Liu, L. Lolla, R. Naresh, T. Pihl, Q. Sun, Y. Wan, Y. Wu, J. Cho, T. DeFreitas, S. Frazer, N. Gehlenborg, G. Getz, D. I. Heiman, J. Kim, M. S. Lawrence, P. Lin, S. Meier, M. S. Noble, G. Saksena, D. Voet, H. Zhang, B. Bernard, N. Chambwe, V. Dhankani, T. Knijnenburg, R. Kramer, K. Leinonen, Y. Liu, M. Miller, S. Reynolds, I. Shmulevich, V. Thorsson, W. Zhang, R. Akbani, B. M. Broom, A. M. Hegde, Z. Ju, R. S. Kanchi, A. Korkut, J. Li, H. Liang, S. Ling, W. Liu, Y. Lu, G. B. Mills, K.-S. Ng, A. Rao, M. Ryan, J. Wang, J. N. Weinstein, J. Zhang, A. Abeshouse, J. Armenia, D. Chakravarty, W. K. Chatila, I. de Bruijn, J. Gao, B. E. Gross, Z. J. Heins, R. Kundra, K. La, M. Ladanyi, A. Luna, M. G. Nissan, A. Ochoa, S. M. Phillips, E. Reznik, F. Sanchez-Vega, C. Sander, N. Schultz, R. Sheridan, S. O. Sumer, Y. Sun, B. S. Taylor, J. Wang, H. Zhang, P. Anur, M. Peto, P. Spellman, C. Benz, J. M. Stuart, C. K. Wong, C. Yau, D. N. Hayes, J. S. Parker, M. D. Wilkerson, A. Ally, M. Balasundaram, R. Bowlby, D. Brooks, R. Carlsen, E. Chuah, N. Dhalla, R. Holt, S. J. M. Jones, K. Kasaian, D. Lee, Y. Ma, M. A. Marra, M. Mayo, R. A. Moore, A. J. Mungall, K. Mungall, A. G. Robertson, S. Sadeghi, J. E. Schein, P. Sipahimalani, A. Tam, N. Thiessen, K. Tse, T. Wong, A. C. Berger, R. Beroukhim, A. D. Cherniack, C. Cibulskis, S. B. Gabriel, G. F. Gao, G. Ha, M. Meyerson, S. E. Schumacher, J. Shih, M. H. Kucherlapati, R. S. Kucherlapati, S. Baylin, L. Cope, L. Danilova, M. S. Bootwalla, P. H. Lai, D. T. Maglinte, D. J. Van Den Berg, D. J. Weisenberger, J. T. Auman, S. Balu, T. Bodenheimer, C. Fan, K. A. Hoadley, A. P. Hoyle, S. R. Jefferys, C. D. Jones, S. Meng, P. A. Mieczkowski, L. E. Mose, A. H. Perou, C. M. Perou, J. Roach, Y. Shi, J. V. Simons, T. Skelly, M. G. Soloway, D. Tan, U. Veluvolu, H. Fan, T. Hinoue, P. W. Laird, H. Shen, W. Zhou, M. Bellair, K. Chang, K. Covington, C. J. Creighton, H. Dinh, H. Doddapaneni, L. A. Donehower, J. Drummond, R. A. Gibbs, R. Glenn, W. Hale, Y. Han, J. Hu, V. Korchina, S. Lee, L. Lewis, W. Li, X. Liu, M. Morgan, D. Morton, D. Muzny, J. Santibanez, M. Sheth, E. Shinbrot, L. Wang, M. Wang, D. A. Wheeler, L. Xi, F. Zhao, J. Hess, E. L. Appelbaum, M. Bailey, M. G. Cordes, L. Ding, C. C. Fronick, L. A. Fulton, R. S. Fulton, C. Kandoth, E. R. Mardis, M. D. McLellan, C. A. Miller, H. K. Schmidt, R. K. Wilson, D. Crain, E. Curley, J. Gardner, K. Lau, D. Mallery, S. Morris, J. Paulauskis, R. Penny, C. Shelton, T. Shelton, M. Sherman, E. Thompson, P. Yena, J. Bowen, J. M. Gastier-Foster, M. Gerken, K. M. Leraas, T. M. Lichtenberg, N. C. Ramirez, L. Wise, E. Zmuda, N. Corcoran, T. Costello, C. Hovens, A. L. Carvalho, A. C. de Carvalho, J. H. Fregnani, A. Longatto-Filho, R. M. Reis, C. Scapulatempo-Neto, H. C. S. Silveira, D. O. Vidal, A. Burnette, J. Eschbacher, B. Hermes, A. Noss, R. Singh, M. L. Anderson, P. D. Castro, M. Ittmann, D. Huntsman, B. Kohl, X. Le, R. Thorp, C. Andry, E. R. Duffy, V. Lyadov, O. Paklina, G. Setdikova, A. Shabunin, M. Tavobilov, C. McPherson, R. Warnick, R. Berkowitz, D. Cramer, C. Feltmate, N. Horowitz, A. Kibel, M. Muto, C. P. Raut, A. Malykh, J. S. Barnholtz-Sloan, W. Barrett, K. Devine, J. Fulop, Q. T. Ostrom, K. Shimmel, Y. Wolinsky, A. E. Sloan, A. De Rose, F. Giuliante, M. Goodman, B. Y. Karlan, C. H. Hagedorn, J. Eckman, J. Harr, J. Myers, K. Tucker, L. A. Zach, B. Deyarmin, H. Hu, L. Kvecher, C. Larson, R. J. Mural, S. Somiari, A. Vicha, T. Zelinka, J. Bennett, M. Iacocca, B. Rabeno, P. Swanson, M. Latour, L. Lacombe, B. Têtu, A. Bergeron, M. McGraw, S. M. Staugaitis, J. Chabot, H. Hibshoosh, A. Sepulveda, T. Su, T. Wang, O. Potapova, O. Voronina, L. Desjardins, O. Mariani, S. Roman-Roman, X. Sastre, M.-H. Stern, F. Cheng, S. Signoretti, A. Berchuck, D. Bigner, E. Lipp, J. Marks, S. McCall, R. McLendon, A. Secord, A. Sharp, M. Behera, D. J. Brat, A. Chen, K. Delman, S. Force, F. Khuri, K. Magliocca, S. Maithel, J. J. Olson, T. Owonikoko, A. Pickens, S. Ramalingam, D. M. Shin, G. Sica, E. G. Van Meir, H. Zhang, W. Eijckenboom, A. Gillis, E. Korpershoek, L. Looijenga, W. Oosterhuis, H. Stoop, K. E. van Kessel, E. C. Zwarthoff, C. Calatozzolo, L. Cuppini, S. Cuzzubbo, F. DiMeco, G. Finocchiaro, L. Mattei, A. Perin, B. Pollo, C. Chen, J. Houck, P. Lohavanichbutr, A. Hartmann, C. Stoehr, R. Stoehr, H. Taubert, S. Wach, B. Wullich, W. Kycler, D. Murawa, M. Wiznerowicz, K. Chung, W. J. Edenfield, J. Martin, E. Baudin, G. Bubley, R. Bueno, A. De Rienzo, W. G. Richards, S. Kalkanis, T. Mikkelsen, H. Noushmehr, L. Scarpace, N. Girard, M. Aymerich, E. Campo, E. Giné, A. L. Guillermo, N. Van Bang, P. T. Hanh, B. D. Phu, Y. Tang, H. Colman, K. Evason, P. R. Dottino, J. A. Martignetti, H. Gabra, H. Juhl, T. Akeredolu, S. Stepa, D. Hoon, K. Ahn, K. J. Kang, F. Beuschlein, A. Breggia, M. Birrer, D. Bell, M. Borad, A. H. Bryce, E. Castle, V. Chandan, J. Cheville, J. A. Copland, M. Farnell, T. Flotte, N. Giama, T. Ho, M. Kendrick, J.-P. Kocher, K. Kopp, C. Moser, D. Nagorney, D. O’Brien, B. P. O’Neill, T. Patel, G. Petersen, F. Que, M. Rivera, L. Roberts, R. Smallridge, T. Smyrk, M. Stanton, R. H. Thompson, M. Torbenson, J. D. Yang, L. Zhang, F. Brimo, J. A. Ajani, A. M. A. Gonzalez, C. Behrens, J. Bondaruk, R. Broaddus, B. Czerniak, B. Esmaeli, J. Fujimoto, J. Gershenwald, C. Guo, A. J. Lazar, C. Logothetis, F. Meric-Bernstam, C. Moran, L. Ramondetta, D. Rice, A. Sood, P. Tamboli, T. Thompson, P. Troncoso, A. Tsao, I. Wistuba, C. Carter, L. Haydu, P. Hersey, V. Jakrot, H. Kakavand, R. Kefford, K. Lee, G. Long, G. Mann, M. Quinn, R. Saw, R. Scolyer, K. Shannon, A. Spillane, onathan Stretch, M. Synott, J. Thompson, J. Wilmott, H. Al-Ahmadie, T. A. Chan, R. Ghossein, A. Gopalan, D. A. Levine, V. Reuter, S. Singer, B. Singh, N. V. Tien, T. Broudy, C. Mirsaidi, P. Nair, P. Drwiega, J. Miller, J. Smith, H. Zaren, J.-W. Park, N. P. Hung, E. Kebebew, W. M. Linehan, A. R. Metwalli, K. Pacak, P. A. Pinto, M. Schiffman, L. S. Schmidt, C. D. Vocke, N. Wentzensen, R. Worrell, H. Yang, M. Moncrieff, C. Goparaju, J. Melamed, H. Pass, N. Botnariuc, I. Caraman, M. Cernat, I. Chemencedji, A. Clipca, S. Doruc, G. Gorincioi, S. Mura, M. Pirtac, I. Stancul, D. Tcaciuc, M. Albert, I. Alexopoulou, A. Arnaout, J. Bartlett, J. Engel, S. Gilbert, J. Parfitt, H. Sekhon, G. Thomas, D. M. Rassl, R. C. Rintoul, C. Bifulco, R. Tamakawa, W. Urba, N. Hayward, H. Timmers, A. Antenucci, F. Facciolo, G. Grazi, M. Marino, R. Merola, R. de Krijger, A.-P. Gimenez-Roqueplo, A. Piché, S. Chevalier, G. McKercher, K. Birsoy, G. Barnett, C. Brewer, C. Farver, T. Naska, N. A. Pennell, D. Raymond, C. Schilero, K. Smolenski, F. Williams, C. Morrison, J. A. Borgia, M. J. Liptay, M. Pool, C. W. Seder, K. Junker, L. Omberg, M. Dinkin, G. Manikhas, D. Alvaro, M. C. Bragazzi, V. Cardinale, G. Carpino, E. Gaudio, D. Chesla, S. Cottingham, M. Dubina, F. Moiseenko, R. Dhanasekaran, K.-F. Becker, K.-P. Janssen, J. Slotta-Huspenina, M. H. Abdel-Rahman, D. Aziz, S. Bell, C. M. Cebulla, A. Davis, R. Duell, J. B. Elder, J. Hilty, B. Kumar, J. Lang, N. L. Lehman, R. Mandt, P. Nguyen, R. Pilarski, K. Rai, L. Schoenfield, K. Senecal, P. Wakely, P. Hansen, R. Lechan, J. Powers, A. Tischler, W. E. Grizzle, K. C. Sexton, A. Kastl, J. Henderson, S. Porten, J. Waldmann, M. Fassnacht, S. L. Asa, D. Schadendorf, M. Couce, M. Graefen, H. Huland, G. Sauter, T. Schlomm, R. Simon, P. Tennstedt, O. Olabode, M. Nelson, O. Bathe, P. R. Carroll, J. M. Chan, P. Disaia, P. Glenn, R. K. Kelley, C. N. Landen, J. Phillips, M. Prados, J. Simko, K. Smith-McCune, S. VandenBerg, K. Roggin, A. Fehrenbach, A. Kendler, S. Sifri, R. Steele, A. Jimeno, F. Carey, I. Forgie, M. Mannelli, M. Carney, B. Hernandez, B. Campos, C. Herold-Mende, C. Jungk, A. Unterberg, A. von Deimling, A. Bossler, J. Galbraith, L. Jacobus, M. Knudson, T. Knutson, D. Ma, M. Milhem, R. Sigmund, A. K. Godwin, R. Madan, H. G. Rosenthal, C. Adebamowo, S. N. Adebamowo, A. Boussioutas, D. Beer, T. Giordano, A.-M. Mes-Masson, F. Saad, T. Bocklage, L. Landrum, R. Mannel, K. Moore, K. Moxley, R. Postier, J. Walker, R. Zuna, M. Feldman, F. Valdivieso, R. Dhir, J. Luketich, E. M. M. Pinero, M. Quintero-Aguilo, C. G. Carlotti, J. S. Dos Santos, R. Kemp, A. Sankarankuty, D. Tirapelli, J. Catto, K. Agnew, E. Swisher, J. Creaney, B. Robinson, C. S. Shelley, E. M. Godwin, S. Kendall, C. Shipman, C. Bradford, T. Carey, A. Haddad, J. Moyer, L. Peterson, M. Prince, L. Rozek, G. Wolf, R. Bowman, K. M. Fong, I. Yang, R. Korst, W. K. Rathmell, J. L. Fantacone-Campbell, J. A. Hooke, A. J. Kovatich, C. D. Shriver, J. DiPersio, B. Drake, R. Govindan, S. Heath, T. Ley, B. Van Tine, P. Westervelt, M. A. Rubin, J. I. Lee, N. D. Aredes, A. Mariamidze, J. M. Stuart, P. W. Laird, K. A. Hoadley, J. N. Weinstein, M. Peto, C. R. Pickering, Z. Chen, C. Van Waes, Genomic, Pathway Network, and Immunologic Features Distinguishing Squamous Carcinomas. Cell Reports. 23, 194-212.e6 (2018).

10. K. E. Yoh, K. Regunath, A. Guzman, S.-M. Lee, N. T. Pfister, O. Akanni, L. J. Kaufman, C. Prives, R. Prywes, Repression of p63 and induction of EMT by mutant Ras in mammary epithelial cells. Proc. Natl. Acad. Sci. U.S.A. 113, E6107–E6116 (2016).

11. L. Hu, S. Liang, H. Chen, T. Lv, J. Wu, D. Chen, M. Wu, S. Sun, H. Zhang, H. You, H. Ji, Y. Zhang, J. Bergholz, Z.-X. J. Xiao, ΔNp63α is a common inhibitory target in oncogenic PI3K/Ras/Her2-induced cell motility and tumor metastasis. Proc Natl Acad Sci USA. 114, E3964–E3973 (2017).

12. R. Chakrabarti, Y. Wei, J. Hwang, X. Hang, M. Andres Blanco, A. Choudhury, B. Tiede, R.-A. Romano, C. DeCoste, L. Mercatali, T. Ibrahim, D. Amadori, N. Kannan, C. J. Eaves, S. Sinha, Y. Kang, ΔNp63 promotes stem cell activity in mammary gland development and basal-like breast cancer by enhancing Fzd7 expression and Wnt signalling. Nature Cell Biology. 16, 1004–1015 (2014).

13. P. Orzol, M. Nekulova, J. Holcakova, P. Muller, B. Votesek, P. J. Coates, ΔNp63 regulates cell proliferation, differentiation, adhesion, and migration in the BL2 subtype of basal-like breast cancer. Tumor Biol. 37, 10133–10140 (2016).

14. L. Wang, Z. Zhang, X. Yu, X. Huang, Z. Liu, Y. Chai, L. Yang, Q. Wang, M. Li, J. Zhao, J. Hou, F. Li, Unbalanced YAP-SOX9 circuit drives stemness and malignant progression in esophageal squamous cell carcinoma. Oncogene. 38, 2042–2055 (2019).

15. S. W. Chan, C. J. Lim, K. Guo, C. P. Ng, I. Lee, W. Hunziker, Q. Zeng, W. Hong, A Role for TAZ in Migration, Invasion, and Tumorigenesis of Breast Cancer Cells. Cancer Res. 68, 2592–2598 (2008).

16. M. Cordenonsi, F. Zanconato, L. Azzolin, M. Forcato, A. Rosato, C. Frasson, M. Inui, M. Montagner, A. R. Parenti, A. Poletti, M. G. Daidone, S. Dupont, G. Basso, S. Bicciato, S. Piccolo, The Hippo Transducer TAZ Confers Cancer Stem Cell-Related Traits on Breast Cancer Cells. Cell. 147, 759–772 (2011).

17. F. Zanconato, M. Forcato, G. Battilana, L. Azzolin, E. Quaranta, B. Bodega, A. Rosato, S. Bicciato, M. Cordenonsi, S. Piccolo, Genome-wide association between YAP/TAZ/TEAD and AP-1 at enhancers drives oncogenic growth. Nat Cell Biol. 17, 1218–1227 (2015).

18. J. Li, Z. Li, Y. Wu, Y. Wang, D. Wang, W. Zhang, H. Yuan, J. Ye, X. Song, J. Yang, H. Jiang, J. Cheng, The Hippo effector TAZ promotes cancer stemness by transcriptional activation of SOX2 in head neck squamous cell carcinoma. Cell Death Dis. 10, 1–15 (2019).

19. B. Zhao, X. Wei, W. Li, R. S. Udan, Q. Yang, J. Kim, J. Xie, T. Ikenoue, J. Yu, L. Li, P. Zheng, K. Ye, A. Chinnaiyan, G. Halder, Z.-C. Lai, K.-L. Guan, Inactivation of YAP oncoprotein by the Hippo pathway is involved in cell contact inhibition and tissue growth control. Genes & Development. 21, 2747–2761 (2007).

20. Q.-Y. Lei, H. Zhang, B. Zhao, Z.-Y. Zha, F. Bai, X.-H. Pei, S. Zhao, Y. Xiong, K.-L. Guan, TAZ promotes cell proliferation and epithelial-mesenchymal transition and is inhibited by the hippo pathway. Mol Cell Biol. 28, 2426–2436 (2008).

21. E. H. Y. Chan, M. Nousiainen, R. B. Chalamalasetty, A. Schäfer, E. A. Nigg, H. H. W. Silljé, The Ste20-like kinase Mst2 activates the human large tumor suppressor kinase Lats1. Oncogene. 24, 2076–2086 (2005).

22. Y. Hao, A. Chun, K. Cheung, B. Rashidi, X. Yang, Tumor Suppressor LATS1 Is a Negative Regulator of Oncogene YAP. Journal of Biological Chemistry. 283, 5496– 5509 (2008).

23. B. Zhao, L. Li, K. Tumaneng, C.-Y. Wang, K.-L. Guan, A coordinated phosphorylation by Lats and CK1 regulates YAP stability through SCF -TRCP. Genes & Development. 24, 72–85 (2010).

24. B. Zhao, X. Ye, J. Yu, L. Li, W. Li, S. Li, J. Yu, J. D. Lin, C.-Y. Wang, A. M. Chinnaiyan, Z.-C. Lai, K.-L. Guan, TEAD mediates YAP-dependent gene induction and growth control. Genes Dev. 22, 1962–1971 (2008).

25. H. Zhang, C.-Y. Liu, Z.-Y. Zha, B. Zhao, J. Yao, S. Zhao, Y. Xiong, Q.-Y. Lei, K.-L. Guan, TEAD transcription factors mediate the function of TAZ in cell growth and epithelial-mesenchymal transition. J. Biol. Chem. 284, 13355–13362 (2009).

26. A. Hergovich, D. Schmitz, B. A. Hemmings, The human tumour suppressor LATS1 is activated by human MOB1 at the membrane. Biochemical and Biophysical Research Communications. 345, 50–58 (2006).

27. M. Praskova, F. Xia, J. Avruch, MOBKL1A/MOBKL1B phosphorylation by MST1 and MST2 inhibits cell proliferation. Curr Biol. 18, 311–321 (2008).

28. B. A. Callus, A. M. Verhagen, D. L. Vaux, Association of mammalian sterile twenty kinases, Mst1 and Mst2, with hSalvador via C-terminal coiled-coil domains, leads to its stabilization and phosphorylation. The FEBS Journal. 273, 4264–4276 (2006).

29. S. J. Bae, L. Ni, A. Osinski, D. R. Tomchick, C. A. Brautigam, X. Luo, SAV1 promotes Hippo kinase activation through antagonizing the PP2A phosphatase STRIPAK. Elife. 6 (2017), doi:10.7554/eLife.30278.

30. Z. Lin, R. Xie, K. Guan, M. Zhang, A WW Tandem-Mediated Dimerization Mode of SAV1 Essential for Hippo Signaling. Cell Reports. 32, 108118 (2020).

31. N. Furth, Y. Aylon, M. Oren, p53 shades of Hippo. Cell Death & Differentiation. 25, 81–92 (2018).

32. H. D. Soule, T. M. Maloney, S. R. Wolman, W. D. Peterson, R. Brenz, C. M. McGrath, J. Russo, R. J. Pauley, R. F. Jones, S. C. Brooks, Isolation and characterization of a spontaneously immortalized human breast epithelial cell line, MCF-10. Cancer research. 50, 6075–6086 (1990).

33. F. Fan, Z. He, L.-L. Kong, Q. Chen, Q. Yuan, S. Zhang, J. Ye, H. Liu, X. Sun, J. Geng, L. Yuan, L. Hong, C. Xiao, W. Zhang, X. Sun, Y. Li, P. Wang, L. Huang, X. Wu, Z. Ji, Q. Wu, N.-S. Xia, N. S. Gray, L. Chen, C.-H. Yun, X. Deng, D. Zhou, Pharmacological targeting of kinases MST1 and MST2 augments tissue repair and regeneration. Science Translational Medicine. 8, 352ra108–352ra108 (2016).

34. Y. Aylon, D. Michael, A. Shmueli, N. Yabuta, H. Nojima, M. Oren, A positive feedback loop between the p53 and Lats2 tumor suppressors prevents tetraploidization. Genes Dev. 20, 2687–2700 (2006).

35. N. Furth, N. Bossel Ben-Moshe, Y. Pozniak, Z. Porat, T. Geiger, E. Domany, Y. Aylon, M. Oren, Down-regulation of LATS kinases alters p53 to promote cell migration. Genes Dev. 29, 2325–2330 (2015).

36. N. Kastan, K. Gnedeva, T. Alisch, A. A. Petelski, D. J. Huggins, J. Chiaravalli, A. Aharanov, A. Shakked, E. Tzahor, A. Nagiel, N. Segil, A. J. Hudspeth, Small-molecule inhibition of Lats kinases may promote Yap-dependent proliferation in postmitotic mammalian tissues. Nat Commun. 12, 3100 (2021).

37. V. Gatti, C. Fierro, M. Compagnone, F. Giangrazi, E. K. Markert, L. Bongiorno-Borbone, G. Melino, A. Peschiaroli, ΔNp63 regulates the expression of hyaluronic acid-related genes in breast cancer cells. Oncogenesis. 7, 65 (2018).

38. Y. Liu-Chittenden, B. Huang, J. S. Shim, Q. Chen, S.-J. Lee, R. A. Anders, J. O. Liu, D. Pan, Genetic and pharmacological disruption of the TEAD–YAP complex suppresses the oncogenic activity of YAP. Genes Dev. 26, 1300–1305 (2012).

39. I. Y. Luk, C. M. Reehorst, J. M. Mariadason, ELF3, ELF5, EHF and SPDEF Transcription Factors in Tissue Homeostasis and Cancer. Molecules. 23, 2191 (2018).

40. F. Li, W. Zhu, F. J. Gonzalez, Potential role of CYP1B1 in the development and treatment of metabolic diseases. Pharmacology & Therapeutics. 178, 18–30 (2017).

41. F. Tang, R. Gao, B. Jeevan-Raj, C. B. Wyss, R. K. R. Kalathur, S. Piscuoglio, C. K. Y. Ng, S. K. Hindupur, S. Nuciforo, E. Dazert, T. Bock, S. Song, D. Buechel, M. F. Morini, A. Hergovich, P. Matthias, D.-S. Lim, L. M. Terracciano, M. H. Heim, M. N. Hall, G. Christofori, LATS1 but not LATS2 represses autophagy by a kinase-independent scaffold function. Nat Commun. 10, 5755 (2019).

42. S. Xiong, A. L. Couzens, M. J. Kean, D. Y. Mao, S. Guettler, I. Kurinov, A.-C. Gingras, F. Sicheri, Regulation of Protein Interactions by Mps One Binder (MOB1) Phosphorylation. Mol Cell Proteomics. 16, 1111–1125 (2017).

43. L. Ni, Y. Zheng, M. Hara, D. Pan, X. Luo, Structural basis for Mob1-dependent activation of the core Mst-Lats kinase cascade in Hippo signaling. Genes Dev. 29, 1416–1431 (2015).

44. E. S. Stavridi, K. G. Harris, Y. Huyen, J. Bothos, P.-M. Verwoerd, S. E. Stayrook, N. P. Pavletich, P. D. Jeffrey, F. C. Luca, Crystal Structure of a Human Mob1 Protein: Toward Understanding Mob-Regulated Cell Cycle Pathways. Structure. 11, 1163–1170 (2003).

45. F. Yin, J. Yu, Y. Zheng, Q. Chen, N. Zhang, D. Pan, Spatial organization of Hippo signaling at the plasma membrane mediated by the tumor suppressor Merlin/NF2. Cell. 154, 1342–1355 (2013).

46. Y.-S. Jung, Y. Qian, W. Yan, X. Chen, Pirh2 E3 ubiquitin ligase modulates keratinocyte differentiation through p63. J Invest Dermatol. 133, 1178–1187 (2013).

47. Y. Li, Z. Zhou, C. Chen, WW domain-containing E3 ubiquitin protein ligase 1 targets p63 transcription factor for ubiquitin-mediated proteasomal degradation and regulates apoptosis. Cell Death & Differentiation. 15, 1941–1951 (2008).

48. M. Rossi, R. I. Aqeilan, M. Neale, E. Candi, P. Salomoni, R. A. Knight, C. M. Croce, G. Melino, The E3 ubiquitin ligase Itch controls the protein stability of p63. Proc Natl Acad Sci U S A. 103, 12753–12758 (2006).

49. K. C. Ho, Z. Zhou, Y.-M. She, A. Chun, T. D. Cyr, X. Yang, Itch E3 ubiquitin ligase regulates large tumor suppressor 1 stability [corrected]. Proc Natl Acad Sci U S A. 108, 4870–4875 (2011).

50. M. L. Fisher, S. Balinth, A. A. Mills, p63-related signaling at a glance. Journal of Cell Science. 133, jcs228015 (2020).

51. Y. Wang, J. Li, Y. Gao, Y. Luo, H. Luo, L. Wang, Y. Yi, Z. Yuan, Z.-X. Jim Xiao, Hippo kinases regulate cell junctions to inhibit tumor metastasis in response to oxidative stress. Redox Biol. 26 (2019), doi:10.1016/j.redox.2019.101233.

52. I. Valencia-Sama, Y. Zhao, D. Lai, H. J. Janse van Rensburg, Y. Hao, X. Yang, Hippo Component TAZ Functions as a Co-repressor and Negatively Regulates ΔNp63 Transcription through TEA Domain (TEAD) Transcription Factor. J Biol Chem. 290, 16906–16917 (2015).

53. A. Britschgi, S. Duss, S. Kim, J. P. Couto, H. Brinkhaus, S. Koren, D. De Silva, K. D. Mertz, D. Kaup, Z. Varga, H. Voshol, A. Vissieres, C. Leroy, T. Roloff, M. B. Stadler, C. H. Scheel, L. J. Miraglia, A. P. Orth, G. M. C. Bonamy, V. A. Reddy, M. Bentires-Alj, The Hippo kinases LATS1 and 2 control human breast cell fate via crosstalk with ERα. Nature. 541, 541–545 (2017).

54. Y. Aylon, A. Gershoni, R. Rotkopf, I. E. Biton, Z. Porat, A. P. Koh, X. Sun, Y. Lee, M.-I. Fiel, Y. Hoshida, S. L. Friedman, R. L. Johnson, M. Oren, The LATS2 tumor suppressor inhibits SREBP and suppresses hepatic cholesterol accumulation. Genes Dev. 30, 786–797 (2016).

55. D. Venkatesh, N. A. O’Brien, F. Zandkarimi, D. R. Tong, M. E. Stokes, D. E. Dunn, E. S. Kengmana, A. T. Aron, A. M. Klein, J. M. Csuka, S.-H. Moon, M. Conrad, C. J. Chang, D. C. Lo, A. D’Alessandro, C. Prives, B. R. Stockwell, MDM2 and MDMX promote ferroptosis by PPARα-mediated lipid remodeling. Genes Dev. 34, 526– 543 (2020).

