## Supplemental files for "A non-canonical Hippo pathway represses the expression of ΔNp63"

**Supplementary Figures**

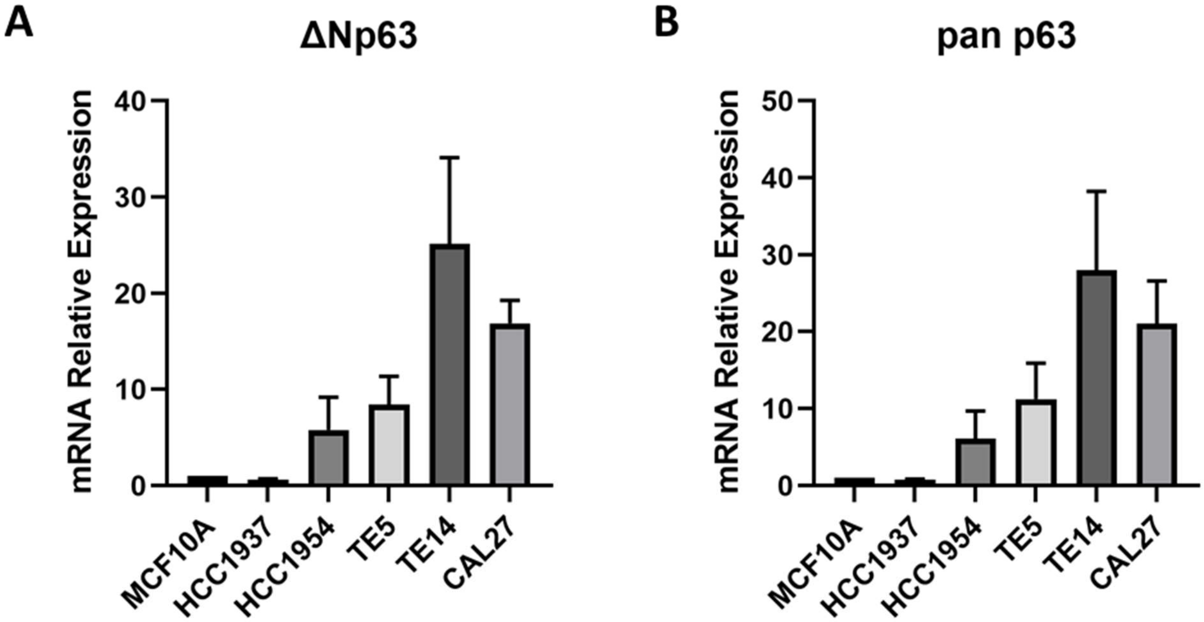


**Supplemental Fig. 1. Comparison of p63 mRNA levels across cell lines**

**A.** RT-qPCR with ΔNp63 specific isoform primers. Expression was normalized to MCF10A levels. **B.** RT-qPCR for p63 with pan isoform primers. Expression was normalized to MCF10A levels. Bars represent the mean ± SEM of four independent experiments.

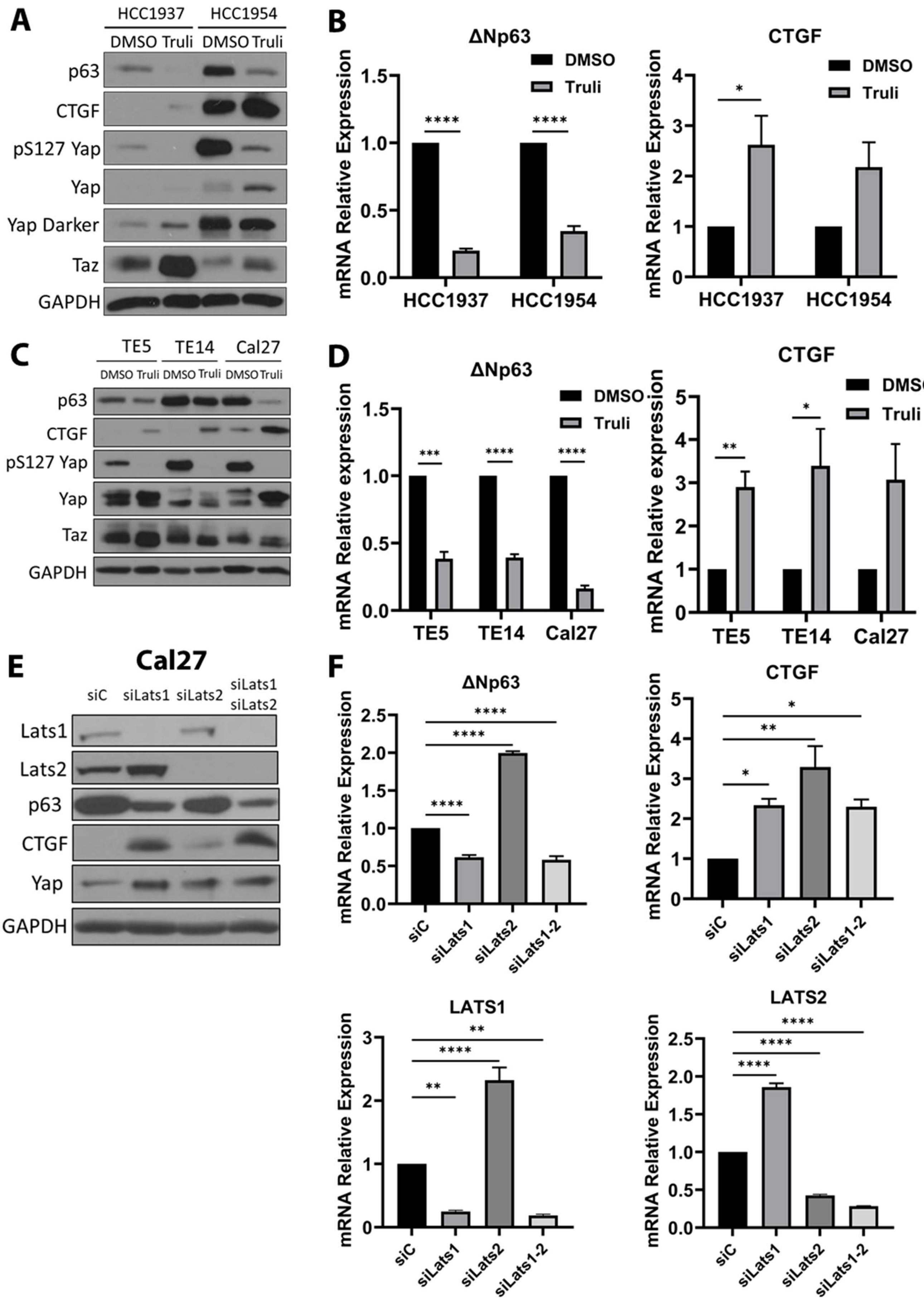

A
HCC1937
HCC1954
DMSO Truli DMSO Truli
p63
CTGF
pS127 Yap
Yap
Yap Darker
Taz
GAPDH
B
ΔNp63
CTGF
DMSO
Truli
mRNA Relative Expression
HCC1937
HCC1954
C
TE5
TE14
Cal27
DMSO Truli DMSO Truli DMSO Truli
p63
CTGF
pS127 Yap
Yap
Taz
GAPDH
D
ΔNp63
CTGF
mRNA Relative expression
TE5
TE14
Cal27
E
Cal27
siC
siLats1
siLats2
siLats1 siLats2
Lats1
Lats2
p63
CTGF
Yap
GAPDH
F
ΔNp63
CTGF
LATS1
LATS2
mRNA Relative Expression
siC
siLats1
siLats2
siLats1-2

**Supplemental Fig. 2. LATS regulate p63 expression in breast and squamous cancer cell lines**

**A.** Immunoblot of breast cancer cells treated with 20 uM Truli for 24 h. **B.** RT-qPCR of breast cancer cells treated as shown in A. **C.** Immunoblot of squamous carcinoma cells treated with Truli for 24 h. **D.** RT-qPCR of squamous carcinoma cells treated as shown in B. **E.** Immunoblot of Cal27 squamous carcinoma cell line transfected with siControl, siLATS, siLATS2, or both siLATS1-2 smartpool siRNA for 72 h. **F.** RT-qPCR of Cal27 squamous carcinoma cells treated as shown in E. For the immunoblots, one representative of three independent experiments is shown. For RT-qPCR, bars represent the mean ± SEM of three independent experiments with three technical replicates normalized to the DMSO treatment. Statistical analysis was Student t-test between DMSO and treatment. Statistical significance is represented in the graphs as: * $p<0.05$, ** $p<0.01$, *** $p<0.005$, **** $p<0.001$.

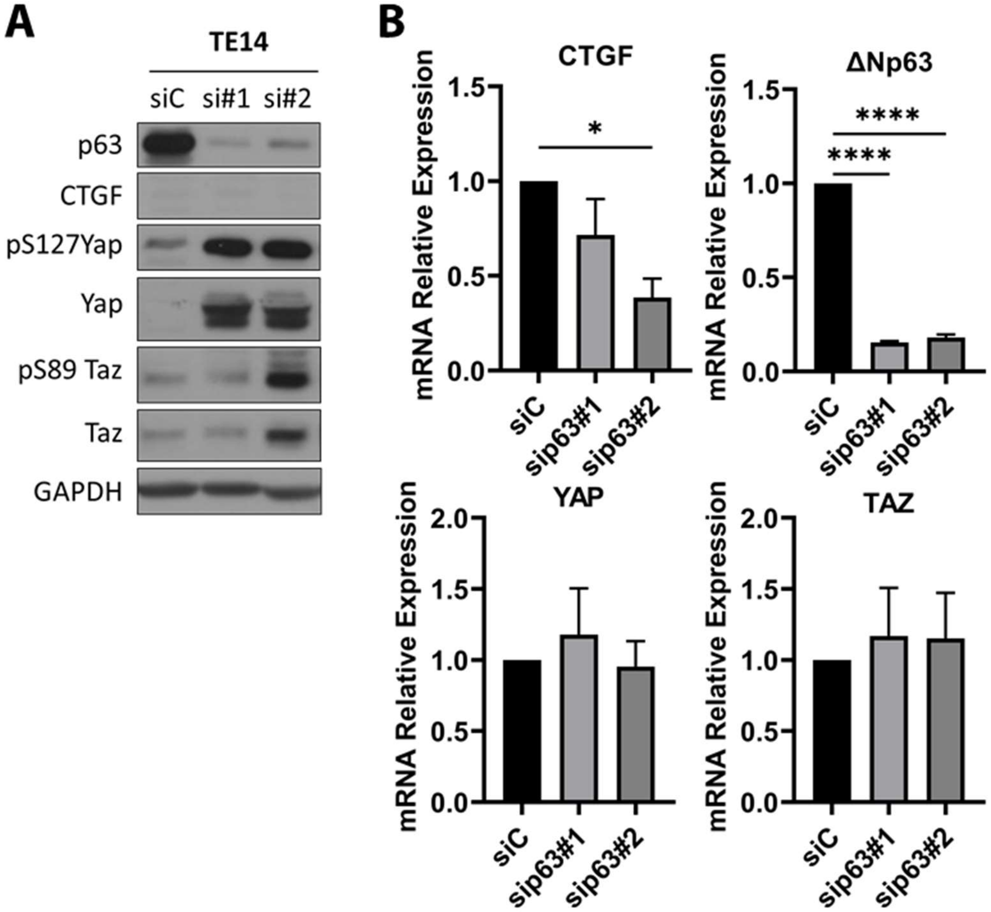


**Supplemental Fig. 3. p63 inhibits YAP activity.**

**A.** Immunoblot of TE14 cells transfected with control siRNA (siC) or two siRNAs against p63 for 72h. **B.** RT-qPCR of same cells in A. Immunoblots is a representative experiment of three independent experiments. For RT-qPCR, bars represent the mean or dNp63 expression the statistical analysis was Stechnical replicates, normalized to siC. Statistical analysis was One-Way ANOVA with Dunnett's multiple comparisons test against the siC. Statistical significance is represented in the graphs as: * p<0.05, ** p<0.01, *** p<0.005, **** p<0.001.

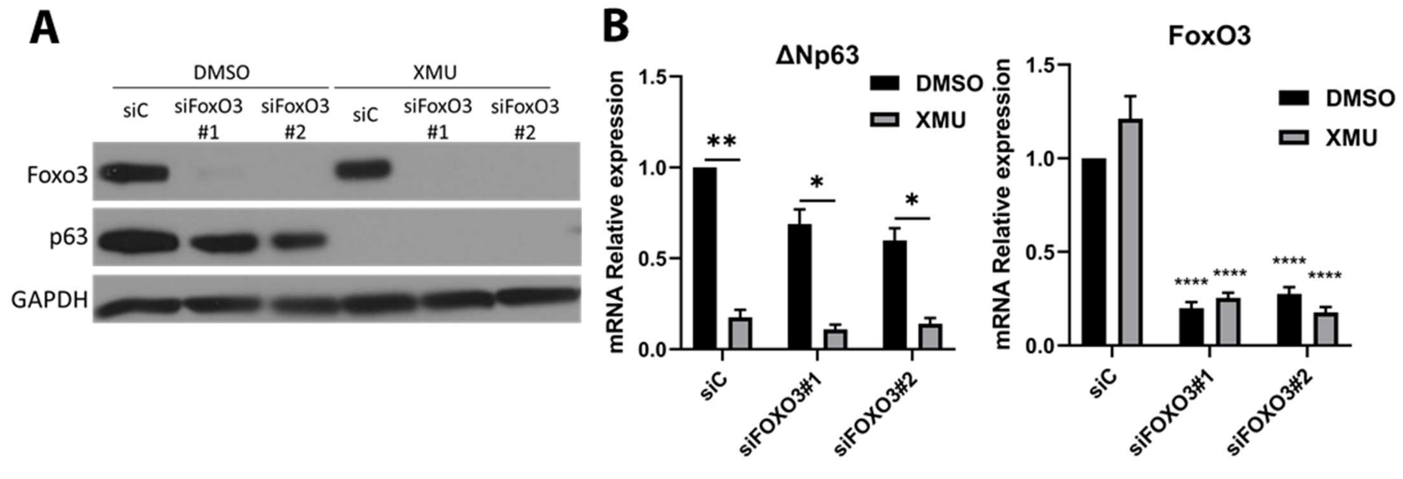


**Supplemental Fig. 4. FoxO3 is not required for ΔNp63 expression in MCF10A cells.**

**A.** Immunoblot of MCF10A cells that were transfected with control or two different FoxO3a-specific siRNAs for 48h; cells were treated for 24h with 5 μM of XMU-MP1. The immunoblot is of one representative experiment of three independendent. **B.** qRT-PCR for ΔNp63 and FoxO3 as MCF10A cells treated as in A. Bars represent the mean relative expression ± SEM of three independent experiments with three technical replicates. For ∆Np63 expression the statistical analysis was Student's t-test comparing DMSO vs XMU treatment. For FOXO3 expression the statistical analysis was ANOVA with Dunett's test of multiple comparisons against the siC DMSO treatment. Statistical significance is represented in the graphs as: * $p<0.05$, ** $p<0.01$, *** $p<0.005$, **** $p<0.001$.

**Supplemental Table 1. Primer sequences used for qRT-PCR**

| Gene | Forward sequence | Reverse sequence |
|---|---|---|
| RPL32 | TTCCTGGTCCACAACGTCAAG | TGTGAGCGATCTCGGCAC |
| CTGF | GGAGTGGGTGTGTGACGAG | CTTCCAGTCGGTAAGCCGC |
| ΔNp63 | AAACAATGCCCAGACTCAATTT | GTGGTCTGTGTTATAGGGACTG |
| Panp63 | AAACCAGAGATGGGCAAGTC | TCGTACCATCACCGTTCTTTG |
| MST1 | CAATGCAGAGGATGAGGAAGAG | TCTTGCCAAAGCTGTTGATCT |
| MST2 | GCATGAACCCTTCCCTATGTC | AACCGCATCTGTAGTTCTTCTAAA |
| LATS1 | AAATCTCATCAGCAGCGTCTAC | CTCATTTGATCCTGGGCATCTT |
| LATS2 | TGGTCACATTAAACTCACAGATTTC | GTTAGACACATCATCCCAGAGG |
| SAV1 | TCTGGTACCTGCAAATCCATATC | GGAAGAGTTCCCACTTCAGAAT |
| MOB1A | CCCGTGAGCTAAGGACGGT | AGGAAGCTCATCTTCGGTCC |
| MOB1B | GTCTCCTGTTCCATTCGCCT | AAGAAGCTCATGTTGGCCGC |
| YAP | TTTTGGCTGCCACCAAGCTA | AATTCAGTCTGCCTGAGGGC |
| WWTR1 (Taz) | GGAGCTCATGAGGCAGGAAG | CCTGAACTGGGGCAAGAGTC |
| FOXO3 | GGGCAAAGCAGACCCTCAA | GTCCACTTGCTGAGAGCAGA |
| 18S | | |
| CYP1B1 | TCCTCTTCACCAGGTATCC | CCACGACCTGATCCAATTC |
| ELF3 | CTCTGCAATTGTGCCCTTGA | CATCAGAAGAGCTGGAAGTGAG |
| IL1A | GTGCTGCTGAAGGAGATG | GTGAGTTTCCCAGAAGAAGAG |
| IL1B | GCCTGAAGCCCTTGCTGTGT | GCGGCATCCAGCTACGAAT |
| PLAU | GATTCCTGCCAGGGAGA | GTGTGAGACTCTCGTGTAGA |
| TRIM16 | GTCCAGTTCTTGGAGGAGTA | CCAGCAACTGGATTAAGTGTA |
| TRIM47 | AGCTCAGCTTCACCAAATC | GCTCAGCATCAGCTTCC |
| TSKU | GGACCCAATGCACTTTCT | CTGCTACTCCTGTGCTAATG |

**Supplemental Table 2A. Genes significantly repressed by XMU**

Repressed genes table sorted for XMU fold change over DMSO treated siC and $p<0.05$.

**Supplemental Table 2B. Genes significantly repressed by XMU independent of YAP/TAZ**

Repressed genes independent of YAP/TAZ. Genes were sorted based on XMU fold change expression of 0.5 over the siC DMSO treated and siYAP/TAZ XMU treated of least 0.33-fold change and $p<0.05$.

**Supplemental Table 2C. Genes induced by XMU**

Induced genes table sorted based on XMU treated siC 3-fold change over DMSO treated siC and $p<0.2$.

**Supplemental Table 2D. Genes significantly induced by XMU dependent upon YAP/TAZ.**

Genes were sorted based on XMU fold change expression of 3-fold over the siC DMSO treated condition (with $p<0.2$) and whose expression in siYAP/TAZ XMU treated cells was less than 50% of that in siC XMU treated cells.

**Supplemental Table 2A: Genes significantly repressed by XMU.**

Repressed genes table sorted for XMU fold change over DMSO treated siC and p<0.05.

| Gene Name | FPKM siC | FPKM siC + XMU | FPKM siYap/ siTaz | FPKM siYap/ siTaz + XMU | XMU Fold Change in siC cells | p value | XMU Fold change in siYap/siTaz cells |
|---|---|---|---|---|---|---|---|
| BDKRB1 | 2.36 | 0.00 | 3.25 | 0.71 | 0.000 | 0.001 | 0.217 |
| TICAM2 | 1.21 | 0.00 | 0.40 | 0.00 | 0.000 | 0.008 | 0.000 |
| LOC112268278 | 1.09 | 0.00 | 0.20 | 0.11 | 0.000 | 0.009 | 0.569 |
| MIR6510 | 2.36 | 0.02 | 5.42 | 0.28 | 0.006 | 0.008 | 0.052 |
| LOC91370 | 2.45 | 0.07 | 0.72 | 0.56 | 0.027 | 0.028 | 0.779 |
| BDKRB2 | 3.47 | 0.10 | 5.99 | 1.35 | 0.029 | 0.000 | 0.225 |
| KCTD4 | 2.39 | 0.08 | 0.19 | 0.06 | 0.035 | 0.013 | 0.300 |
| LOC105377276 | 1.19 | 0.05 | 0.14 | 0.29 | 0.040 | 0.016 | 2.124 |
| SERTAD4 | 2.10 | 0.09 | 0.50 | 0.08 | 0.042 | 0.010 | 0.157 |
| IL7R | 3.80 | 0.17 | 0.43 | 0.20 | 0.044 | 0.049 | 0.455 |
| ZBED2 | 10.68 | 0.50 | 1.24 | 0.09 | 0.046 | 0.015 | 0.070 |
| MIG7 | 19.87 | 0.93 | 1.52 | 0.40 | 0.047 | 0.006 | 0.265 |
| HCAR2 | 5.07 | 0.25 | 31.22 | 3.65 | 0.050 | 0.046 | 0.117 |
| PDGFB | 6.65 | 0.35 | 3.30 | 0.12 | 0.053 | 0.005 | 0.038 |
| CYP1B1 | 10.59 | 0.58 | 20.66 | 4.43 | 0.054 | 0.028 | 0.214 |
| ADGRF1 | 5.82 | 0.32 | 4.86 | 2.68 | 0.055 | 0.003 | 0.550 |
| SH3TC1 | 5.47 | 0.34 | 12.53 | 1.54 | 0.062 | 0.002 | 0.123 |
| GRHL3 | 6.69 | 0.46 | 68.45 | 2.07 | 0.068 | 0.040 | 0.030 |
| HRCT1 | 7.95 | 0.56 | 3.16 | 0.15 | 0.070 | 0.002 | 0.048 |
| KRT80 | 33.02 | 2.41 | 78.06 | 62.03 | 0.073 | 0.008 | 0.795 |
| LINC00899 | 2.28 | 0.17 | 0.24 | 0.01 | 0.075 | 0.040 | 0.049 |
| JADE2 | 15.96 | 1.29 | 5.86 | 1.87 | 0.081 | 0.007 | 0.319 |
| CYP1A1 | 2.71 | 0.22 | 4.55 | 1.54 | 0.082 | 0.006 | 0.338 |
| ADGRF4 | 3.98 | 0.33 | 17.67 | 5.68 | 0.082 | 0.014 | 0.321 |
| TSKU | 46.32 | 3.87 | 39.06 | 2.39 | 0.084 | 0.002 | 0.061 |
| MANSC1 | 1.00 | 0.09 | 3.28 | 2.22 | 0.088 | 0.002 | 0.678 |
| GLIS2 | 2.63 | 0.23 | 0.65 | 0.10 | 0.089 | 0.002 | 0.147 |
| NIPAL4 | 6.50 | 0.58 | 22.07 | 0.53 | 0.089 | 0.000 | 0.024 |
| KCNN4 | 37.92 | 3.40 | 37.44 | 18.67 | 0.090 | 0.002 | 0.499 |
| RAB7B | 2.13 | 0.19 | 13.53 | 2.08 | 0.091 | 0.049 | 0.154 |
| TNS3 | 4.79 | 0.44 | 0.64 | 0.16 | 0.092 | 0.012 | 0.253 |
| KRT7-AS | 16.36 | 1.53 | 13.05 | 1.28 | 0.093 | 0.005 | 0.098 |
| LINC00963 | 1.01 | 0.10 | 0.79 | 0.20 | 0.094 | 0.011 | 0.251 |
| CCDC9B | 15.68 | 1.48 | 17.53 | 5.70 | 0.095 | 0.029 | 0.325 |
| NKX1-2 | 3.81 | 0.36 | 0.06 | 0.00 | 0.095 | 0.036 | 0.002 |
| PLAU | 154.48 | 14.84 | 99.22 | 7.21 | 0.096 | 0.021 | 0.073 |
| TFAP4 | 6.35 | 0.63 | 0.26 | 0.09 | 0.099 | 0.023 | 0.351 |
| GNAO1 | 1.00 | 0.10 | 0.47 | 0.36 | 0.101 | 0.021 | 0.773 |
| TNIP3 | 1.47 | 0.15 | 0.19 | 0.06 | 0.102 | 0.043 | 0.325 |
| IL1B | 17.63 | 1.85 | 45.05 | 11.81 | 0.105 | 0.022 | 0.262 |
| BEND3 | 1.30 | 0.14 | 0.21 | 0.10 | 0.108 | 0.014 | 0.474 |
| GRAMD2A | 4.67 | 0.51 | 1.56 | 0.31 | 0.109 | 0.046 | 0.197 |

| | | | | | | | |
|---|---|---|---|---|---|---|---|
| P2RY6 | 1.18 | 0.13 | 0.06 | 0.02 | 0.109 | 0.004 | 0.326 |
| BHLHE41 | 1.66 | 0.18 | 2.30 | 0.22 | 0.110 | 0.008 | 0.098 |
| MIR6835 | 4.50 | 0.50 | 3.75 | 1.01 | 0.110 | 0.003 | 0.271 |
| NOG | 1.79 | 0.20 | 0.09 | 0.06 | 0.111 | 0.002 | 0.621 |
| TP63 | 4.17 | 0.46 | 2.46 | 0.59 | 0.111 | 0.005 | 0.239 |
| IL1A | 57.41 | 6.41 | 77.51 | 2.27 | 0.112 | 0.024 | 0.029 |
| FJX1 | 22.69 | 2.62 | 1.93 | 0.20 | 0.115 | 0.000 | 0.105 |
| C1QTNF6 | 2.09 | 0.24 | 0.28 | 0.24 | 0.115 | 0.018 | 0.858 |
| LOC101928837 | 1.61 | 0.19 | 4.89 | 0.11 | 0.116 | 0.009 | 0.023 |
| IFIT3 | 13.48 | 1.61 | 1.96 | 0.28 | 0.119 | 0.002 | 0.144 |
| HOXC8 | 2.13 | 0.27 | 3.50 | 0.71 | 0.127 | 0.001 | 0.202 |
| TRN-GTT11-1 | 1.45 | 0.19 | 0.64 | 0.00 | 0.128 | 0.010 | 0.000 |
| ZHX2 | 3.18 | 0.41 | 1.44 | 0.60 | 0.128 | 0.010 | 0.416 |
| MDFI | 9.60 | 1.24 | 10.42 | 4.24 | 0.129 | 0.000 | 0.407 |
| KLF11 | 1.29 | 0.17 | 2.10 | 0.70 | 0.129 | 0.002 | 0.335 |
| ADORA2B | 11.57 | 1.49 | 2.10 | 0.46 | 0.129 | 0.003 | 0.218 |
| BACE1-AS | 1.37 | 0.18 | 0.67 | 0.31 | 0.129 | 0.012 | 0.464 |
| ETV7 | 2.24 | 0.29 | 0.75 | 0.10 | 0.130 | 0.002 | 0.139 |
| IQANK1 | 3.94 | 0.52 | 12.55 | 5.79 | 0.131 | 0.002 | 0.461 |
| NNT-AS1 | 1.22 | 0.16 | 0.34 | 0.11 | 0.131 | 0.011 | 0.316 |
| TNS4 | 56.98 | 7.49 | 43.13 | 4.71 | 0.131 | 0.014 | 0.109 |
| C10orf55 | 1.61 | 0.21 | 2.24 | 0.35 | 0.132 | 0.001 | 0.155 |
| CSF1 | 14.83 | 1.97 | 4.89 | 0.84 | 0.133 | 0.029 | 0.172 |
| JPH2 | 2.48 | 0.33 | 0.05 | 0.05 | 0.134 | 0.005 | 0.917 |
| EPN3 | 13.42 | 1.80 | 64.00 | 18.88 | 0.134 | 0.014 | 0.295 |
| STRA6 | 1.49 | 0.20 | 5.81 | 2.36 | 0.135 | 0.023 | 0.407 |
| TRIM47 | 26.01 | 3.50 | 18.60 | 1.83 | 0.135 | 0.020 | 0.099 |
| AMIGO2 | 6.19 | 0.85 | 1.98 | 0.31 | 0.138 | 0.000 | 0.156 |
| HNRNPA3P6 | 2.28 | 0.31 | 0.25 | 1.34 | 0.138 | 0.001 | 5.454 |
| VNN1 | 4.21 | 0.58 | 10.20 | 3.29 | 0.139 | 0.006 | 0.322 |
| LINC01186 | 1.21 | 0.17 | 0.00 | 0.00 | 0.141 | 0.034 | 1.000 |
| MLKL | 12.90 | 1.86 | 4.91 | 2.21 | 0.144 | 0.017 | 0.450 |
| PHC1P1 | 2.94 | 0.43 | 1.16 | 1.38 | 0.145 | 0.028 | 1.184 |
| BHLHE40 | 40.71 | 6.15 | 50.03 | 27.53 | 0.151 | 0.000 | 0.550 |
| EPB41L1 | 1.50 | 0.23 | 1.75 | 1.22 | 0.152 | 0.006 | 0.700 |
| TRIM16L | 13.12 | 2.00 | 11.97 | 3.72 | 0.152 | 0.010 | 0.311 |
| BIRC3 | 39.86 | 6.08 | 11.10 | 1.84 | 0.153 | 0.033 | 0.166 |
| LOC100505549 | 1.17 | 0.18 | 0.44 | 0.26 | 0.153 | 0.005 | 0.588 |
| SOGA1 | 10.09 | 1.54 | 0.94 | 0.53 | 0.153 | 0.004 | 0.564 |
| EFNA1 | 48.14 | 7.36 | 38.38 | 11.36 | 0.153 | 0.026 | 0.296 |
| SLC2A12 | 1.21 | 0.19 | 0.23 | 0.13 | 0.153 | 0.033 | 0.542 |
| CRACDL | 1.10 | 0.17 | 1.55 | 0.36 | 0.153 | 0.002 | 0.229 |
| NECTIN4 | 6.72 | 1.03 | 68.85 | 4.53 | 0.154 | 0.039 | 0.066 |
| MIR1915HG | 1.31 | 0.21 | 0.69 | 0.13 | 0.159 | 0.001 | 0.193 |
| C1orf116 | 8.44 | 1.34 | 16.05 | 1.53 | 0.159 | 0.014 | 0.095 |
| PARTICL | 1.25 | 0.20 | 0.82 | 0.13 | 0.159 | 0.018 | 0.153 |
| P2RY2 | 1.93 | 0.31 | 4.24 | 1.03 | 0.160 | 0.003 | 0.244 |

| | | | | | | | |
|---|---|---|---|---|---|---|---|
| TGFB2 | 1.68 | 0.27 | 0.07 | 0.04 | 0.160 | 0.024 | 0.608 |
| IFIT2 | 3.83 | 0.62 | 0.46 | 0.11 | 0.161 | 0.011 | 0.234 |
| APOL3 | 2.60 | 0.42 | 0.75 | 0.22 | 0.162 | 0.014 | 0.296 |
| RIN2 | 1.62 | 0.26 | 0.97 | 0.04 | 0.162 | 0.007 | 0.041 |
| PWWP2B | 5.30 | 0.86 | 8.09 | 0.74 | 0.163 | 0.003 | 0.091 |
| ARHGEF10L | 1.30 | 0.21 | 5.77 | 1.00 | 0.165 | 0.001 | 0.173 |
| NCR3LG1 | 1.17 | 0.19 | 0.24 | 0.15 | 0.166 | 0.028 | 0.629 |
| CYSRT1 | 23.11 | 3.88 | 122.91 | 41.93 | 0.168 | 0.011 | 0.341 |
| GJB4 | 9.91 | 1.67 | 83.42 | 23.48 | 0.168 | 0.022 | 0.282 |
| ZMYND8 | 8.85 | 1.50 | 11.74 | 1.48 | 0.169 | 0.000 | 0.126 |
| THBD | 2.53 | 0.43 | 8.90 | 0.17 | 0.172 | 0.025 | 0.019 |
| ITPKB | 1.16 | 0.20 | 2.51 | 0.45 | 0.172 | 0.033 | 0.178 |
| CXCL2 | 14.01 | 2.42 | 1.38 | 1.25 | 0.173 | 0.007 | 0.908 |
| TRIM16 | 37.82 | 6.55 | 71.94 | 14.87 | 0.173 | 0.002 | 0.207 |
| GPRC5C | 2.95 | 0.51 | 2.96 | 0.60 | 0.173 | 0.010 | 0.203 |
| BCL3 | 10.83 | 1.88 | 13.14 | 0.90 | 0.173 | 0.025 | 0.069 |
| SH3PXD2A-AS1 | 5.47 | 0.96 | 42.91 | 17.80 | 0.175 | 0.000 | 0.415 |
| DHRS3 | 51.55 | 9.03 | 13.39 | 5.46 | 0.175 | 0.000 | 0.407 |
| IL6R | 2.35 | 0.41 | 10.71 | 3.27 | 0.175 | 0.030 | 0.306 |
| FAT2 | 6.50 | 1.15 | 5.45 | 2.80 | 0.177 | 0.001 | 0.515 |
| EEF1AKMT3 | 2.64 | 0.47 | 0.72 | 0.24 | 0.177 | 0.021 | 0.335 |
| KCTD15 | 3.28 | 0.58 | 2.00 | 1.77 | 0.178 | 0.041 | 0.881 |
| LRRC61 | 3.13 | 0.56 | 0.73 | 0.38 | 0.179 | 0.025 | 0.522 |
| LOC105372578 | 1.92 | 0.35 | 6.07 | 1.49 | 0.180 | 0.050 | 0.245 |
| DLK2 | 3.82 | 0.69 | 0.43 | 0.18 | 0.181 | 0.001 | 0.418 |
| INHBA | 3.68 | 0.69 | 2.41 | 1.35 | 0.186 | 0.002 | 0.560 |
| PAK6 | 6.40 | 1.20 | 14.58 | 4.91 | 0.188 | 0.003 | 0.336 |
| RIN3 | 6.11 | 1.15 | 3.10 | 2.35 | 0.189 | 0.002 | 0.756 |
| AMER1 | 1.25 | 0.24 | 0.76 | 0.16 | 0.189 | 0.004 | 0.209 |
| LZTS2 | 7.40 | 1.42 | 8.69 | 1.31 | 0.191 | 0.009 | 0.151 |
| ZBTB16 | 1.12 | 0.22 | 1.48 | 0.43 | 0.192 | 0.000 | 0.288 |
| DTX4 | 11.43 | 2.20 | 3.52 | 0.54 | 0.193 | 0.002 | 0.154 |
| MYEOV | 10.10 | 1.95 | 10.99 | 3.26 | 0.194 | 0.028 | 0.297 |
| MCTS1 | 2.03 | 0.39 | 0.34 | 0.96 | 0.194 | 0.026 | 2.800 |
| DKK1 | 17.38 | 3.41 | 1.63 | 0.15 | 0.196 | 0.018 | 0.091 |
| LRFN4 | 15.14 | 2.98 | 10.49 | 5.78 | 0.197 | 0.001 | 0.551 |
| TMC5 | 2.48 | 0.49 | 0.95 | 0.58 | 0.197 | 0.008 | 0.607 |
| ARHGEF16 | 10.71 | 2.11 | 8.14 | 3.39 | 0.197 | 0.003 | 0.416 |
| EVI2B | 2.40 | 0.48 | 0.12 | 0.11 | 0.198 | 0.015 | 0.871 |
| LINC00920 | 1.20 | 0.24 | 0.36 | 0.08 | 0.199 | 0.032 | 0.236 |
| CXCL1 | 102.08 | 20.66 | 20.58 | 18.98 | 0.202 | 0.007 | 0.922 |
| LINC00857 | 5.40 | 1.11 | 1.18 | 0.41 | 0.205 | 0.001 | 0.345 |
| DIXDC1 | 1.33 | 0.27 | 0.56 | 0.25 | 0.205 | 0.000 | 0.449 |
| B3GNT3 | 23.01 | 4.75 | 25.08 | 11.26 | 0.206 | 0.000 | 0.449 |
| ZNF532 | 2.22 | 0.46 | 1.71 | 0.85 | 0.207 | 0.003 | 0.496 |
| SYT12 | 2.49 | 0.51 | 12.72 | 5.94 | 0.207 | 0.004 | 0.467 |
| PLEKHA7 | 2.98 | 0.62 | 2.09 | 0.46 | 0.207 | 0.031 | 0.218 |

| | | | | | | | |
|---|---|---|---|---|---|---|---|
| RARG | 21.22 | 4.42 | 80.94 | 9.86 | 0.209 | 0.004 | 0.122 |
| LINC01704 | 1.41 | 0.29 | 0.10 | 0.04 | 0.209 | 0.037 | 0.394 |
| TGIF2 | 8.69 | 1.83 | 0.95 | 0.30 | 0.210 | 0.001 | 0.319 |
| E2F5 | 2.45 | 0.52 | 1.36 | 0.81 | 0.211 | 0.007 | 0.594 |
| SLC12A8 | 1.82 | 0.39 | 0.52 | 0.23 | 0.212 | 0.002 | 0.447 |
| OTUB2 | 1.68 | 0.36 | 21.24 | 3.20 | 0.212 | 0.013 | 0.151 |
| TRAF5 | 1.46 | 0.31 | 0.63 | 0.21 | 0.212 | 0.001 | 0.337 |
| LOC257396 | 1.15 | 0.25 | 0.10 | 0.05 | 0.213 | 0.016 | 0.494 |
| CABLES1 | 3.05 | 0.66 | 1.21 | 0.48 | 0.217 | 0.004 | 0.398 |
| ACTRT3 | 1.43 | 0.31 | 0.37 | 0.06 | 0.217 | 0.008 | 0.169 |
| TFEB | 1.78 | 0.39 | 4.94 | 0.41 | 0.218 | 0.005 | 0.082 |
| IRF1-AS1 | 3.56 | 0.78 | 0.13 | 0.76 | 0.220 | 0.002 | 5.629 |
| GJB3 | 55.64 | 12.28 | 146.95 | 26.65 | 0.221 | 0.006 | 0.181 |
| MEAK7 | 17.26 | 3.81 | 20.14 | 5.68 | 0.221 | 0.001 | 0.282 |
| SAMD10 | 5.86 | 1.29 | 7.36 | 4.50 | 0.221 | 0.003 | 0.612 |
| MYC | 71.15 | 15.76 | 23.85 | 2.28 | 0.222 | 0.003 | 0.096 |
| DENND2D | 2.82 | 0.63 | 2.73 | 1.08 | 0.222 | 0.007 | 0.395 |
| PDZD2 | 1.67 | 0.37 | 2.06 | 1.04 | 0.222 | 0.003 | 0.505 |
| CD14 | 28.89 | 6.44 | 18.21 | 15.45 | 0.223 | 0.011 | 0.849 |
| ADGRG1 | 21.58 | 4.82 | 40.45 | 26.17 | 0.223 | 0.012 | 0.647 |
| FOSL1 | 100.37 | 22.44 | 43.04 | 37.55 | 0.224 | 0.046 | 0.873 |
| RUNX2 | 1.23 | 0.28 | 0.16 | 0.09 | 0.224 | 0.032 | 0.564 |
| CAVIN2 | 35.51 | 8.00 | 5.39 | 2.05 | 0.225 | 0.003 | 0.381 |
| LOC105371795 | 1.66 | 0.37 | 1.04 | 0.45 | 0.225 | 0.004 | 0.431 |
| SLC16A13 | 5.95 | 1.35 | 8.64 | 3.24 | 0.226 | 0.036 | 0.375 |
| KIFC3 | 5.03 | 1.15 | 2.74 | 1.38 | 0.228 | 0.011 | 0.505 |
| TRIM14 | 4.58 | 1.05 | 1.94 | 0.93 | 0.229 | 0.003 | 0.482 |
| AMPD3 | 2.35 | 0.54 | 5.85 | 2.22 | 0.229 | 0.001 | 0.380 |
| MIR100HG | 2.38 | 0.55 | 1.70 | 0.51 | 0.230 | 0.013 | 0.299 |
| GJB5 | 24.02 | 5.52 | 63.28 | 23.61 | 0.230 | 0.020 | 0.373 |
| F3 | 72.96 | 16.81 | 19.91 | 13.66 | 0.230 | 0.011 | 0.686 |
| PPP1R13L | 25.74 | 5.94 | 44.48 | 14.17 | 0.231 | 0.011 | 0.318 |
| TRIB2 | 4.03 | 0.93 | 15.98 | 3.27 | 0.232 | 0.000 | 0.204 |
| CCND1 | 51.55 | 11.98 | 21.51 | 20.56 | 0.232 | 0.008 | 0.956 |
| ZNF618 | 1.09 | 0.25 | 0.23 | 0.07 | 0.232 | 0.001 | 0.316 |
| ZNF747 | 2.39 | 0.56 | 2.65 | 0.21 | 0.233 | 0.009 | 0.079 |
| GNB1L | 1.41 | 0.33 | 0.47 | 0.21 | 0.233 | 0.006 | 0.441 |
| FBXO32 | 6.60 | 1.55 | 23.13 | 10.98 | 0.235 | 0.035 | 0.475 |
| SOX12 | 1.79 | 0.42 | 0.85 | 0.31 | 0.235 | 0.003 | 0.369 |
| MACC1 | 2.03 | 0.48 | 0.41 | 0.21 | 0.235 | 0.027 | 0.518 |
| FLT3LG | 1.78 | 0.42 | 0.38 | 0.11 | 0.235 | 0.001 | 0.290 |
| SVIL | 5.42 | 1.28 | 4.72 | 1.89 | 0.236 | 0.000 | 0.400 |
| NAV1 | 1.81 | 0.43 | 1.59 | 0.26 | 0.237 | 0.003 | 0.165 |
| HNRNPA1P33 | 4.18 | 0.99 | 0.91 | 0.38 | 0.237 | 0.006 | 0.421 |
| RGS3 | 6.42 | 1.53 | 8.38 | 0.92 | 0.238 | 0.024 | 0.109 |
| THRA | 4.27 | 1.02 | 6.40 | 3.21 | 0.239 | 0.029 | 0.501 |
| PIGC | 16.18 | 3.87 | 12.39 | 0.94 | 0.239 | 0.001 | 0.076 |

| | | | | | | | |
|---|---|---|---|---|---|---|---|
| SMAD3 | 15.00 | 3.59 | 10.06 | 2.01 | 0.239 | 0.002 | 0.200 |
| ITPRIPL2 | 6.37 | 1.53 | 2.81 | 0.62 | 0.240 | 0.002 | 0.221 |
| HR | 9.32 | 2.26 | 81.79 | 29.94 | 0.242 | 0.005 | 0.366 |
| DTX3L | 14.23 | 3.46 | 4.94 | 1.42 | 0.243 | 0.000 | 0.288 |
| LNCTAM34A | 2.12 | 0.52 | 0.72 | 0.26 | 0.244 | 0.002 | 0.364 |
| TLR3 | 3.86 | 0.95 | 0.70 | 0.19 | 0.245 | 0.003 | 0.267 |
| MLPH | 1.84 | 0.45 | 3.00 | 0.82 | 0.245 | 0.000 | 0.274 |
| CAMKK1 | 1.31 | 0.32 | 1.55 | 0.54 | 0.246 | 0.015 | 0.348 |
| INAVA | 20.09 | 4.94 | 28.76 | 2.81 | 0.246 | 0.003 | 0.098 |
| CLMN | 3.06 | 0.75 | 0.45 | 0.28 | 0.246 | 0.027 | 0.622 |
| CLCF1 | 3.25 | 0.80 | 7.83 | 0.28 | 0.246 | 0.036 | 0.036 |
| IL31RA | 1.07 | 0.26 | 0.20 | 0.40 | 0.247 | 0.003 | 1.973 |
| UTP18 | 17.64 | 4.37 | 7.01 | 2.60 | 0.248 | 0.000 | 0.371 |
| LINP1 | 11.52 | 2.86 | 0.67 | 0.16 | 0.248 | 0.000 | 0.235 |
| TRIP6 | 95.65 | 23.74 | 27.88 | 15.66 | 0.248 | 0.000 | 0.562 |
| ALKBH8 | 3.88 | 0.97 | 1.46 | 0.32 | 0.249 | 0.001 | 0.216 |
| SBNO2 | 10.83 | 2.72 | 11.67 | 2.50 | 0.251 | 0.003 | 0.214 |
| SLC37A1 | 2.46 | 0.62 | 1.79 | 0.71 | 0.251 | 0.000 | 0.399 |
| LRRC8D | 12.30 | 3.11 | 3.35 | 1.66 | 0.253 | 0.025 | 0.496 |
| IL17D | 1.09 | 0.28 | 0.48 | 0.24 | 0.253 | 0.037 | 0.507 |
| CLCN4 | 1.41 | 0.36 | 1.60 | 0.68 | 0.253 | 0.014 | 0.428 |
| TRIM6 | 9.45 | 2.40 | 2.07 | 0.19 | 0.254 | 0.000 | 0.091 |
| EVA1C | 3.99 | 1.02 | 1.98 | 0.69 | 0.254 | 0.002 | 0.350 |
| TRERF1 | 3.32 | 0.85 | 1.27 | 0.20 | 0.255 | 0.000 | 0.158 |
| ARHGEF37 | 2.28 | 0.58 | 4.73 | 3.72 | 0.255 | 0.034 | 0.788 |
| GPR39 | 10.55 | 2.72 | 7.80 | 1.24 | 0.258 | 0.001 | 0.159 |
| RTN4IP1 | 2.51 | 0.66 | 1.35 | 0.42 | 0.261 | 0.001 | 0.310 |
| MORN4 | 4.97 | 1.30 | 2.86 | 0.94 | 0.261 | 0.022 | 0.328 |
| GTF2IRD1 | 6.62 | 1.73 | 6.97 | 1.34 | 0.261 | 0.000 | 0.192 |
| PPARGC1B | 1.38 | 0.36 | 0.38 | 0.20 | 0.261 | 0.010 | 0.530 |
| ZBED8 | 2.16 | 0.56 | 0.56 | 0.17 | 0.261 | 0.014 | 0.310 |
| ARHGAP45 | 9.28 | 2.43 | 2.28 | 1.90 | 0.262 | 0.024 | 0.832 |
| ELF3 | 59.48 | 15.58 | 60.38 | 12.09 | 0.262 | 0.028 | 0.200 |
| DLGAP1-AS1 | 1.97 | 0.51 | 1.47 | 0.15 | 0.262 | 0.001 | 0.104 |
| MCTS2P | 7.94 | 2.08 | 2.30 | 1.09 | 0.262 | 0.000 | 0.475 |
| SCNN1G | 1.92 | 0.51 | 7.17 | 0.91 | 0.263 | 0.027 | 0.127 |
| CTBP1-DT | 1.86 | 0.49 | 1.08 | 0.36 | 0.263 | 0.034 | 0.333 |
| ANKRD22 | 9.12 | 2.41 | 35.59 | 17.12 | 0.264 | 0.036 | 0.481 |
| RNASE7 | 1.46 | 0.39 | 14.93 | 8.07 | 0.264 | 0.002 | 0.541 |
| FAM83F | 2.89 | 0.77 | 4.10 | 1.60 | 0.265 | 0.000 | 0.390 |
| IKBKE | 12.12 | 3.21 | 9.64 | 3.87 | 0.265 | 0.010 | 0.402 |
| NNMT | 115.77 | 30.71 | 5.36 | 2.70 | 0.265 | 0.000 | 0.503 |
| PITPNM2 | 3.29 | 0.87 | 2.09 | 0.90 | 0.266 | 0.026 | 0.431 |
| EHF | 22.23 | 5.95 | 52.97 | 8.72 | 0.268 | 0.010 | 0.165 |
| GPR87 | 53.01 | 14.20 | 47.17 | 4.89 | 0.268 | 0.012 | 0.104 |
| PRKCQ-AS1 | 2.43 | 0.65 | 0.17 | 0.04 | 0.268 | 0.022 | 0.250 |
| SNORD15B | 3.89 | 1.05 | 0.36 | 2.04 | 0.269 | 0.010 | 5.656 |

| | | | | | | | |
|---|---|---|---|---|---|---|---|
| NFIC | 16.03 | 4.33 | 6.03 | 3.35 | 0.270 | 0.001 | 0.556 |
| ZBED6CL | 1.92 | 0.52 | 2.27 | 1.11 | 0.270 | 0.000 | 0.489 |
| GBP3 | 9.53 | 2.58 | 7.30 | 3.18 | 0.271 | 0.001 | 0.435 |
| L3MBTL3 | 3.86 | 1.04 | 1.16 | 0.22 | 0.271 | 0.001 | 0.192 |
| ALDH1B1 | 19.13 | 5.21 | 3.54 | 2.33 | 0.272 | 0.002 | 0.658 |
| ADAMTS9 | 2.01 | 0.55 | 0.36 | 0.10 | 0.274 | 0.001 | 0.270 |
| KLF9 | 4.04 | 1.11 | 2.88 | 0.84 | 0.275 | 0.013 | 0.293 |
| ELF3-AS1 | 2.96 | 0.81 | 3.64 | 1.10 | 0.275 | 0.026 | 0.302 |
| WDR81 | 1.91 | 0.53 | 2.43 | 0.61 | 0.276 | 0.003 | 0.251 |
| ZFP36L2 | 17.35 | 4.80 | 34.76 | 6.19 | 0.277 | 0.040 | 0.178 |
| RRS1 | 18.15 | 5.02 | 9.32 | 1.48 | 0.277 | 0.006 | 0.159 |
| FAM171A1 | 4.81 | 1.33 | 2.04 | 1.75 | 0.277 | 0.020 | 0.859 |
| INTS3 | 8.10 | 2.24 | 7.59 | 4.67 | 0.277 | 0.042 | 0.616 |
| SH2D4A | 17.16 | 4.79 | 11.65 | 3.41 | 0.279 | 0.007 | 0.293 |
| RAET1E | 1.93 | 0.54 | 4.27 | 3.05 | 0.279 | 0.008 | 0.715 |
| RIN1 | 16.63 | 4.67 | 14.83 | 3.12 | 0.281 | 0.016 | 0.210 |
| AGAP2-AS1 | 7.15 | 2.01 | 4.44 | 3.56 | 0.281 | 0.018 | 0.801 |
| ZNF608 | 3.44 | 0.97 | 0.65 | 0.06 | 0.281 | 0.000 | 0.089 |
| MGA | 1.30 | 0.37 | 0.48 | 0.29 | 0.282 | 0.005 | 0.596 |
| LBHD1 | 4.32 | 1.22 | 0.99 | 2.33 | 0.283 | 0.016 | 2.362 |
| GMPR | 3.76 | 1.08 | 0.77 | 0.64 | 0.286 | 0.006 | 0.839 |
| F2R | 4.69 | 1.34 | 1.53 | 1.19 | 0.286 | 0.002 | 0.780 |
| EPB41L4B | 4.71 | 1.35 | 6.61 | 3.78 | 0.286 | 0.000 | 0.572 |
| IGSF9 | 2.86 | 0.82 | 9.89 | 2.47 | 0.288 | 0.007 | 0.249 |
| NEK3 | 2.25 | 0.65 | 0.92 | 0.41 | 0.289 | 0.000 | 0.446 |
| METTL21A | 2.99 | 0.87 | 1.11 | 0.57 | 0.290 | 0.001 | 0.510 |
| SLC1A3 | 14.72 | 4.27 | 1.27 | 1.28 | 0.290 | 0.019 | 1.005 |
| ACOT11 | 2.64 | 0.77 | 8.06 | 5.28 | 0.290 | 0.001 | 0.656 |
| PARP9 | 3.96 | 1.15 | 1.94 | 0.60 | 0.291 | 0.040 | 0.308 |
| CARD10 | 28.82 | 8.38 | 4.45 | 0.92 | 0.291 | 0.000 | 0.206 |
| PNPLA3 | 4.11 | 1.19 | 5.22 | 2.97 | 0.291 | 0.001 | 0.568 |
| VSTM2L | 8.91 | 2.60 | 1.59 | 1.09 | 0.291 | 0.045 | 0.688 |
| DNER | 2.80 | 0.82 | 14.87 | 13.83 | 0.292 | 0.040 | 0.930 |
| SLC46A1 | 3.90 | 1.14 | 0.71 | 0.51 | 0.292 | 0.002 | 0.722 |
| IL4R | 23.13 | 6.77 | 35.91 | 7.58 | 0.293 | 0.001 | 0.211 |
| CUEDC1 | 4.83 | 1.42 | 1.12 | 0.83 | 0.295 | 0.000 | 0.746 |
| ERVMER34-1 | 14.13 | 4.18 | 0.60 | 0.83 | 0.296 | 0.000 | 1.383 |
| ENC1 | 6.18 | 1.84 | 1.73 | 0.27 | 0.298 | 0.012 | 0.158 |
| C11orf45 | 1.49 | 0.44 | 0.25 | 0.14 | 0.298 | 0.012 | 0.552 |
| CHST14 | 5.22 | 1.56 | 2.10 | 0.54 | 0.298 | 0.002 | 0.256 |
| VSIR | 7.05 | 2.11 | 13.47 | 12.85 | 0.299 | 0.013 | 0.954 |
| NMI | 19.96 | 5.99 | 8.41 | 3.87 | 0.300 | 0.000 | 0.460 |
| MMACHC | 4.15 | 1.25 | 0.46 | 0.60 | 0.301 | 0.002 | 1.307 |
| HOXA3 | 1.86 | 0.56 | 0.53 | 0.06 | 0.301 | 0.007 | 0.121 |
| GBP1 | 8.39 | 2.53 | 4.57 | 3.07 | 0.301 | 0.028 | 0.671 |
| CTU1 | 2.45 | 0.74 | 2.58 | 0.46 | 0.303 | 0.001 | 0.179 |
| PLEKHA2 | 10.33 | 3.14 | 6.06 | 0.85 | 0.304 | 0.005 | 0.140 |

| | | | | | | | |
|---|---|---|---|---|---|---|---|
| LRG1 | 2.64 | 0.80 | 9.21 | 4.72 | 0.304 | 0.032 | 0.513 |
| LASTR | 4.87 | 1.48 | 9.18 | 4.76 | 0.305 | 0.024 | 0.518 |
| CEP41 | 1.90 | 0.58 | 0.44 | 0.16 | 0.305 | 0.001 | 0.357 |
| PDLIM4 | 33.33 | 10.22 | 51.06 | 20.30 | 0.307 | 0.006 | 0.398 |
| ARHGAP23 | 12.10 | 3.71 | 11.16 | 5.67 | 0.307 | 0.041 | 0.508 |
| AFAP1L2 | 11.56 | 3.55 | 3.76 | 1.25 | 0.307 | 0.030 | 0.332 |
| LOC100507002 | 5.17 | 1.59 | 0.49 | 0.00 | 0.309 | 0.002 | 0.000 |
| FAM83A-AS1 | 2.65 | 0.82 | 8.90 | 4.50 | 0.309 | 0.001 | 0.506 |
| SAMD12 | 1.40 | 0.43 | 0.53 | 0.30 | 0.309 | 0.015 | 0.566 |
| CALHM2 | 3.47 | 1.07 | 7.34 | 0.54 | 0.309 | 0.048 | 0.073 |
| SRC | 13.25 | 4.09 | 17.35 | 4.13 | 0.309 | 0.036 | 0.238 |
| SPART | 15.36 | 4.75 | 6.49 | 1.33 | 0.309 | 0.001 | 0.205 |
| STK10 | 29.09 | 8.99 | 31.84 | 7.31 | 0.309 | 0.001 | 0.230 |
| RAP1GAP2 | 4.95 | 1.53 | 5.90 | 4.41 | 0.309 | 0.014 | 0.748 |
| MTA3 | 1.77 | 0.55 | 2.74 | 0.68 | 0.309 | 0.004 | 0.249 |
| MAST4-AS1 | 1.34 | 0.42 | 0.70 | 0.12 | 0.310 | 0.003 | 0.167 |
| SMYD5 | 18.97 | 5.89 | 6.92 | 5.96 | 0.310 | 0.001 | 0.862 |
| TRMT61B | 4.89 | 1.52 | 4.28 | 1.54 | 0.310 | 0.003 | 0.360 |
| CPNE8 | 2.31 | 0.72 | 0.64 | 0.12 | 0.311 | 0.036 | 0.188 |
| LOC102723999 | 1.13 | 0.35 | 1.23 | 0.66 | 0.311 | 0.044 | 0.537 |
| SLC19A1 | 1.63 | 0.51 | 0.51 | 0.45 | 0.311 | 0.002 | 0.873 |
| SECTM1 | 25.66 | 8.02 | 15.17 | 10.53 | 0.312 | 0.009 | 0.694 |
| CRYBG2 | 3.82 | 1.19 | 24.98 | 6.26 | 0.313 | 0.024 | 0.251 |
| FOXQ1 | 4.09 | 1.28 | 2.05 | 1.92 | 0.313 | 0.017 | 0.935 |
| ICAM1 | 5.51 | 1.73 | 3.82 | 6.70 | 0.314 | 0.025 | 1.755 |
| NDST1 | 14.81 | 4.65 | 17.45 | 12.75 | 0.314 | 0.050 | 0.731 |
| SLC43A3 | 161.16 | 50.68 | 11.33 | 11.53 | 0.314 | 0.002 | 1.017 |
| CFLAR | 1.55 | 0.49 | 3.87 | 1.25 | 0.315 | 0.029 | 0.322 |
| COG7 | 7.09 | 2.23 | 7.73 | 2.96 | 0.315 | 0.000 | 0.383 |
| MLLT6 | 5.28 | 1.67 | 2.51 | 2.20 | 0.316 | 0.004 | 0.877 |
| RAI1 | 11.06 | 3.50 | 12.82 | 2.53 | 0.316 | 0.000 | 0.198 |
| TSEN2 | 2.82 | 0.89 | 0.69 | 0.33 | 0.317 | 0.000 | 0.486 |
| NLRX1 | 7.40 | 2.34 | 42.36 | 38.11 | 0.317 | 0.002 | 0.899 |
| PHOSPHO2 | 1.49 | 0.47 | 0.80 | 0.13 | 0.317 | 0.003 | 0.166 |
| SCHIP1 | 8.84 | 2.80 | 2.48 | 0.98 | 0.317 | 0.003 | 0.397 |
| PLEKHA4 | 2.68 | 0.85 | 3.21 | 0.73 | 0.317 | 0.026 | 0.227 |
| PARP12 | 10.45 | 3.31 | 9.09 | 3.96 | 0.317 | 0.042 | 0.436 |
| TRIM22 | 47.40 | 15.04 | 9.13 | 4.65 | 0.317 | 0.001 | 0.509 |
| TOMM40L | 8.60 | 2.73 | 6.37 | 3.80 | 0.317 | 0.008 | 0.597 |
| FHL2 | 14.34 | 4.55 | 8.05 | 1.67 | 0.317 | 0.001 | 0.207 |
| PSTPIP2 | 20.01 | 6.38 | 11.08 | 3.98 | 0.319 | 0.008 | 0.359 |
| CASTOR1 | 4.53 | 1.45 | 10.82 | 7.05 | 0.319 | 0.001 | 0.652 |
| PDGFC | 5.53 | 1.78 | 2.01 | 0.47 | 0.322 | 0.001 | 0.235 |
| IL22RA1 | 1.76 | 0.57 | 6.06 | 3.16 | 0.322 | 0.007 | 0.520 |
| NIPAL2 | 1.41 | 0.45 | 4.49 | 3.54 | 0.322 | 0.005 | 0.788 |
| LOC107986852 | 2.78 | 0.90 | 1.11 | 0.59 | 0.322 | 0.017 | 0.535 |
| KCTD14 | 23.22 | 7.49 | 0.76 | 0.00 | 0.323 | 0.004 | 0.000 |

| | | | | | | | |
|---|---|---|---|---|---|---|---|
| CHST15 | 4.27 | 1.38 | 4.25 | 1.18 | 0.323 | 0.009 | 0.277 |
| CMTM3 | 2.13 | 0.69 | 0.65 | 0.18 | 0.323 | 0.001 | 0.276 |
| LINC02381 | 2.84 | 0.93 | 1.80 | 0.49 | 0.326 | 0.004 | 0.271 |
| EPHX4 | 7.14 | 2.33 | 1.58 | 0.96 | 0.327 | 0.000 | 0.606 |
| NMRAL1 | 7.62 | 2.49 | 1.80 | 4.05 | 0.327 | 0.049 | 2.256 |
| IRAK1BP1 | 1.44 | 0.47 | 0.37 | 0.15 | 0.328 | 0.002 | 0.407 |
| HES1 | 12.29 | 4.05 | 15.24 | 12.21 | 0.329 | 0.003 | 0.801 |
| IRF2BPL | 14.67 | 4.84 | 17.72 | 3.36 | 0.330 | 0.002 | 0.189 |
| DDR1 | 10.69 | 3.52 | 16.39 | 11.31 | 0.330 | 0.005 | 0.690 |

**Supplemental Table 2B. Genes significantly repressed by XMU independent of YAP/TAZ.**
Repressed genes independent of YAP/TAZ. Genes were sorted based on XMU fold change expression of 0.5 over the siC DMSO treated and siYAP/TAZ XMU treated of least 0.33-fold change and p<0.05.

| Gene Name | FPKM siC | FPKM siC + XMU | FPKM siYap/siTaz | FPKM siYap/siTaz + XMU | XMU Fold Change in siC cells | p value | XMU Fold change in siYap/siTaz cells |
|---|---|---|---|---|---|---|---|
| BDKRB1 | 2.36 | 0.00 | 3.25 | 0.71 | 0.000 | 0.001 | 0.217 |
| TICAM2 | 1.21 | 0.00 | 0.40 | 0.00 | 0.000 | 0.008 | 0.000 |
| MIR6510 | 2.36 | 0.02 | 5.42 | 0.28 | 0.006 | 0.008 | 0.052 |
| BDKRB2 | 3.47 | 0.10 | 5.99 | 1.35 | 0.029 | 0.000 | 0.225 |
| KCTD4 | 2.39 | 0.08 | 0.19 | 0.06 | 0.035 | 0.013 | 0.300 |
| SERTAD4 | 2.10 | 0.09 | 0.50 | 0.08 | 0.042 | 0.010 | 0.157 |
| ZBED2 | 10.68 | 0.50 | 1.24 | 0.09 | 0.046 | 0.015 | 0.070 |
| MIG7 | 19.87 | 0.93 | 1.52 | 0.40 | 0.047 | 0.006 | 0.265 |
| HCAR2 | 5.07 | 0.25 | 31.22 | 3.65 | 0.050 | 0.046 | 0.117 |
| PDGFB | 6.65 | 0.35 | 3.30 | 0.12 | 0.053 | 0.005 | 0.038 |
| CYP1B1 | 10.59 | 0.58 | 20.66 | 4.43 | 0.054 | 0.028 | 0.214 |
| SH3TC1 | 5.47 | 0.34 | 12.53 | 1.54 | 0.062 | 0.002 | 0.123 |
| GRHL3 | 6.69 | 0.46 | 68.45 | 2.07 | 0.068 | 0.040 | 0.030 |
| HRCT1 | 7.95 | 0.56 | 3.16 | 0.15 | 0.070 | 0.002 | 0.048 |
| LINC008 | 2.28 | 0.17 | 0.24 | 0.01 | 0.075 | 0.040 | 0.049 |
| JADE2 | 15.96 | 1.29 | 5.86 | 1.87 | 0.081 | 0.007 | 0.319 |
| ADGRF4 | 3.98 | 0.33 | 17.67 | 5.68 | 0.082 | 0.014 | 0.321 |
| TSKU | 46.32 | 3.87 | 39.06 | 2.39 | 0.084 | 0.002 | 0.061 |
| GLIS2 | 2.63 | 0.23 | 0.65 | 0.10 | 0.089 | 0.002 | 0.147 |
| NIPAL4 | 6.50 | 0.58 | 22.07 | 0.53 | 0.089 | 0.000 | 0.024 |
| RAB7B | 2.13 | 0.19 | 13.53 | 2.08 | 0.091 | 0.049 | 0.154 |
| TNS3 | 4.79 | 0.44 | 0.64 | 0.16 | 0.092 | 0.012 | 0.253 |
| KRT7-AS | 16.36 | 1.53 | 13.05 | 1.28 | 0.093 | 0.005 | 0.098 |
| LINC009 | 1.01 | 0.10 | 0.79 | 0.20 | 0.094 | 0.011 | 0.251 |
| CCDC9B | 15.68 | 1.48 | 17.53 | 5.70 | 0.095 | 0.029 | 0.325 |
| NKX1-2 | 3.81 | 0.36 | 0.06 | 0.00 | 0.095 | 0.036 | 0.002 |
| PLAU | 154.48 | 14.84 | 99.22 | 7.21 | 0.096 | 0.021 | 0.073 |
| TNIP3 | 1.47 | 0.15 | 0.19 | 0.06 | 0.102 | 0.043 | 0.325 |
| IL1B | 17.63 | 1.85 | 45.05 | 11.81 | 0.105 | 0.022 | 0.262 |
| GRAMD2A | 4.67 | 0.51 | 1.56 | 0.31 | 0.109 | 0.046 | 0.197 |
| P2RY6 | 1.18 | 0.13 | 0.06 | 0.02 | 0.109 | 0.004 | 0.326 |
| BHLHE41 | 1.66 | 0.18 | 2.30 | 0.22 | 0.110 | 0.008 | 0.098 |
| MIR6835 | 4.50 | 0.50 | 3.75 | 1.01 | 0.110 | 0.003 | 0.271 |
| TP63 | 4.17 | 0.46 | 2.46 | 0.59 | 0.111 | 0.005 | 0.239 |
| IL1A | 57.41 | 6.41 | 77.51 | 2.27 | 0.112 | 0.024 | 0.029 |
| FJX1 | 22.69 | 2.62 | 1.93 | 0.20 | 0.115 | 0.000 | 0.105 |
| LOC1019 | 1.61 | 0.19 | 4.89 | 0.11 | 0.116 | 0.009 | 0.023 |
| IFIT3 | 13.48 | 1.61 | 1.96 | 0.28 | 0.119 | 0.002 | 0.144 |
| HOXC8 | 2.13 | 0.27 | 3.50 | 0.71 | 0.127 | 0.001 | 0.202 |
| TRN-GTT | 1.45 | 0.19 | 0.64 | 0.00 | 0.128 | 0.010 | 0.000 |
| ADORA2B | 11.57 | 1.49 | 2.10 | 0.46 | 0.129 | 0.003 | 0.218 |
| ETV7 | 2.24 | 0.29 | 0.75 | 0.10 | 0.130 | 0.002 | 0.139 |

| | | | | | | | |
|---|---|---|---|---|---|---|---|
| NNT-AS1 | 1.22 | 0.16 | 0.34 | 0.11 | 0.131 | 0.011 | 0.316 |
| TNS4 | 56.98 | 7.49 | 43.13 | 4.71 | 0.131 | 0.014 | 0.109 |
| C10orf5 | 1.61 | 0.21 | 2.24 | 0.35 | 0.132 | 0.001 | 0.155 |
| CSF1 | 14.83 | 1.97 | 4.89 | 0.84 | 0.133 | 0.029 | 0.172 |
| EPN3 | 13.42 | 1.80 | 64.00 | 18.88 | 0.134 | 0.014 | 0.295 |
| TRIM47 | 26.01 | 3.50 | 18.60 | 1.83 | 0.135 | 0.020 | 0.099 |
| AMIGO2 | 6.19 | 0.85 | 1.98 | 0.31 | 0.138 | 0.000 | 0.156 |
| VNN1 | 4.21 | 0.58 | 10.20 | 3.29 | 0.139 | 0.006 | 0.322 |
| TRIM16L | 13.12 | 2.00 | 11.97 | 3.72 | 0.152 | 0.010 | 0.311 |
| BIRC3 | 39.86 | 6.08 | 11.10 | 1.84 | 0.153 | 0.033 | 0.166 |
| EFNA1 | 48.14 | 7.36 | 38.38 | 11.36 | 0.153 | 0.026 | 0.296 |
| CRACDL | 1.10 | 0.17 | 1.55 | 0.36 | 0.153 | 0.002 | 0.229 |
| NECTIN4 | 6.72 | 1.03 | 68.85 | 4.53 | 0.154 | 0.039 | 0.066 |
| MIR1915 | 1.31 | 0.21 | 0.69 | 0.13 | 0.159 | 0.001 | 0.193 |
| C1orf11 | 8.44 | 1.34 | 16.05 | 1.53 | 0.159 | 0.014 | 0.095 |
| PARTICL | 1.25 | 0.20 | 0.82 | 0.13 | 0.159 | 0.018 | 0.153 |
| P2RY2 | 1.93 | 0.31 | 4.24 | 1.03 | 0.160 | 0.003 | 0.244 |
| IFIT2 | 3.83 | 0.62 | 0.46 | 0.11 | 0.161 | 0.011 | 0.234 |
| APOL3 | 2.60 | 0.42 | 0.75 | 0.22 | 0.162 | 0.014 | 0.296 |
| RIN2 | 1.62 | 0.26 | 0.97 | 0.04 | 0.162 | 0.007 | 0.041 |
| PWWP2B | 5.30 | 0.86 | 8.09 | 0.74 | 0.163 | 0.003 | 0.091 |
| ARHGEF1 | 1.30 | 0.21 | 5.77 | 1.00 | 0.165 | 0.001 | 0.173 |
| GJB4 | 9.91 | 1.67 | 83.42 | 23.48 | 0.168 | 0.022 | 0.282 |
| ZMYND8 | 8.85 | 1.50 | 11.74 | 1.48 | 0.169 | 0.000 | 0.126 |
| THBD | 2.53 | 0.43 | 8.90 | 0.17 | 0.172 | 0.025 | 0.019 |
| ITPKB | 1.16 | 0.20 | 2.51 | 0.45 | 0.172 | 0.033 | 0.178 |
| TRIM16 | 37.82 | 6.55 | 71.94 | 14.87 | 0.173 | 0.002 | 0.207 |
| GPRC5C | 2.95 | 0.51 | 2.96 | 0.60 | 0.173 | 0.010 | 0.203 |
| BCL3 | 10.83 | 1.88 | 13.14 | 0.90 | 0.173 | 0.025 | 0.069 |
| IL6R | 2.35 | 0.41 | 10.71 | 3.27 | 0.175 | 0.030 | 0.306 |
| LOC1053 | 1.92 | 0.35 | 6.07 | 1.49 | 0.180 | 0.050 | 0.245 |
| AMER1 | 1.25 | 0.24 | 0.76 | 0.16 | 0.189 | 0.004 | 0.209 |
| LZTS2 | 7.40 | 1.42 | 8.69 | 1.31 | 0.191 | 0.009 | 0.151 |
| ZBTB16 | 1.12 | 0.22 | 1.48 | 0.43 | 0.192 | 0.000 | 0.288 |
| DTX4 | 11.43 | 2.20 | 3.52 | 0.54 | 0.193 | 0.002 | 0.154 |
| MYEOV | 10.10 | 1.95 | 10.99 | 3.26 | 0.194 | 0.028 | 0.297 |
| DKK1 | 17.38 | 3.41 | 1.63 | 0.15 | 0.196 | 0.018 | 0.091 |
| LINC009 | 1.20 | 0.24 | 0.36 | 0.08 | 0.199 | 0.032 | 0.236 |
| PLEKHA7 | 2.98 | 0.62 | 2.09 | 0.46 | 0.207 | 0.031 | 0.218 |
| RARG | 21.22 | 4.42 | 80.94 | 9.86 | 0.209 | 0.004 | 0.122 |
| TGIF2 | 8.69 | 1.83 | 0.95 | 0.30 | 0.210 | 0.001 | 0.319 |
| OTUB2 | 1.68 | 0.36 | 21.24 | 3.20 | 0.212 | 0.013 | 0.151 |
| ACTRT3 | 1.43 | 0.31 | 0.37 | 0.06 | 0.217 | 0.008 | 0.169 |
| TFEB | 1.78 | 0.39 | 4.94 | 0.41 | 0.218 | 0.005 | 0.082 |
| GJB3 | 55.64 | 12.28 | 146.95 | 26.65 | 0.221 | 0.006 | 0.181 |
| MEAK7 | 17.26 | 3.81 | 20.14 | 5.68 | 0.221 | 0.001 | 0.282 |
| MYC | 71.15 | 15.76 | 23.85 | 2.28 | 0.222 | 0.003 | 0.096 |

| | | | | | | | |
|---|---|---|---|---|---|---|---|
| MIR100H | 2.38 | 0.55 | 1.70 | 0.51 | 0.230 | 0.013 | 0.299 |
| PPP1R13 | 25.74 | 5.94 | 44.48 | 14.17 | 0.231 | 0.011 | 0.318 |
| TRIB2 | 4.03 | 0.93 | 15.98 | 3.27 | 0.232 | 0.000 | 0.204 |
| ZNF618 | 1.09 | 0.25 | 0.23 | 0.07 | 0.232 | 0.001 | 0.316 |
| ZNF747 | 2.39 | 0.56 | 2.65 | 0.21 | 0.233 | 0.009 | 0.079 |
| FLT3LG | 1.78 | 0.42 | 0.38 | 0.11 | 0.235 | 0.001 | 0.290 |
| NAV1 | 1.81 | 0.43 | 1.59 | 0.26 | 0.237 | 0.003 | 0.165 |
| RGS3 | 6.42 | 1.53 | 8.38 | 0.92 | 0.238 | 0.024 | 0.109 |
| PIGC | 16.18 | 3.87 | 12.39 | 0.94 | 0.239 | 0.001 | 0.076 |
| SMAD3 | 15.00 | 3.59 | 10.06 | 2.01 | 0.239 | 0.002 | 0.200 |
| ITPRIPL | 6.37 | 1.53 | 2.81 | 0.62 | 0.240 | 0.002 | 0.221 |
| DTX3L | 14.23 | 3.46 | 4.94 | 1.42 | 0.243 | 0.000 | 0.288 |
| TLR3 | 3.86 | 0.95 | 0.70 | 0.19 | 0.245 | 0.003 | 0.267 |
| MLPH | 1.84 | 0.45 | 3.00 | 0.82 | 0.245 | 0.000 | 0.274 |
| INAVA | 20.09 | 4.94 | 28.76 | 2.81 | 0.246 | 0.003 | 0.098 |
| CLCF1 | 3.25 | 0.80 | 7.83 | 0.28 | 0.246 | 0.036 | 0.036 |
| LINP1 | 11.52 | 2.86 | 0.67 | 0.16 | 0.248 | 0.000 | 0.235 |
| ALKBH8 | 3.88 | 0.97 | 1.46 | 0.32 | 0.249 | 0.001 | 0.216 |
| SBNO2 | 10.83 | 2.72 | 11.67 | 2.50 | 0.251 | 0.003 | 0.214 |
| TRIM6 | 9.45 | 2.40 | 2.07 | 0.19 | 0.254 | 0.000 | 0.091 |
| TRERF1 | 3.32 | 0.85 | 1.27 | 0.20 | 0.255 | 0.000 | 0.158 |
| GPR39 | 10.55 | 2.72 | 7.80 | 1.24 | 0.258 | 0.001 | 0.159 |
| RTN4IP1 | 2.51 | 0.66 | 1.35 | 0.42 | 0.261 | 0.001 | 0.310 |
| MORN4 | 4.97 | 1.30 | 2.86 | 0.94 | 0.261 | 0.022 | 0.328 |
| GTF2IRD | 6.62 | 1.73 | 6.97 | 1.34 | 0.261 | 0.000 | 0.192 |
| ZBED8 | 2.16 | 0.56 | 0.56 | 0.17 | 0.261 | 0.014 | 0.310 |
| ELF3 | 59.48 | 15.58 | 60.38 | 12.09 | 0.262 | 0.028 | 0.200 |
| DLGAP1-A | 1.97 | 0.51 | 1.47 | 0.15 | 0.262 | 0.001 | 0.104 |
| SCNN1G | 1.92 | 0.51 | 7.17 | 0.91 | 0.263 | 0.027 | 0.127 |
| CTBP1-D | 1.86 | 0.49 | 1.08 | 0.36 | 0.263 | 0.034 | 0.333 |
| EHF | 22.23 | 5.95 | 52.97 | 8.72 | 0.268 | 0.010 | 0.165 |
| GPR87 | 53.01 | 14.20 | 47.17 | 4.89 | 0.268 | 0.012 | 0.104 |
| PRKCQ-A | 2.43 | 0.65 | 0.17 | 0.04 | 0.268 | 0.022 | 0.250 |
| L3MBTL3 | 3.86 | 1.04 | 1.16 | 0.22 | 0.271 | 0.001 | 0.192 |
| ADAMTS9 | 2.01 | 0.55 | 0.36 | 0.10 | 0.274 | 0.001 | 0.270 |
| KLF9 | 4.04 | 1.11 | 2.88 | 0.84 | 0.275 | 0.013 | 0.293 |
| ELF3-AS | 2.96 | 0.81 | 3.64 | 1.10 | 0.275 | 0.026 | 0.302 |
| WDR81 | 1.91 | 0.53 | 2.43 | 0.61 | 0.276 | 0.003 | 0.251 |
| ZFP36L2 | 17.35 | 4.80 | 34.76 | 6.19 | 0.277 | 0.040 | 0.178 |
| RRS1 | 18.15 | 5.02 | 9.32 | 1.48 | 0.277 | 0.006 | 0.159 |
| SH2D4A | 17.16 | 4.79 | 11.65 | 3.41 | 0.279 | 0.007 | 0.293 |
| RIN1 | 16.63 | 4.67 | 14.83 | 3.12 | 0.281 | 0.016 | 0.210 |
| ZNF608 | 3.44 | 0.97 | 0.65 | 0.06 | 0.281 | 0.000 | 0.089 |
| IGSF9 | 2.86 | 0.82 | 9.89 | 2.47 | 0.288 | 0.007 | 0.249 |
| PARP9 | 3.96 | 1.15 | 1.94 | 0.60 | 0.291 | 0.040 | 0.308 |
| CARD10 | 28.82 | 8.38 | 4.45 | 0.92 | 0.291 | 0.000 | 0.206 |
| IL4R | 23.13 | 6.77 | 35.91 | 7.58 | 0.293 | 0.001 | 0.211 |

| | | | | | | | |
|---|---|---|---|---|---|---|---|
| ENC1 | 6.18 | 1.84 | 1.73 | 0.27 | 0.298 | 0.012 | 0.158 |
| CHST14 | 5.22 | 1.56 | 2.10 | 0.54 | 0.298 | 0.002 | 0.256 |
| HOXA3 | 1.86 | 0.56 | 0.53 | 0.06 | 0.301 | 0.007 | 0.121 |
| CTU1 | 2.45 | 0.74 | 2.58 | 0.46 | 0.303 | 0.001 | 0.179 |
| PLEKHA2 | 10.33 | 3.14 | 6.06 | 0.85 | 0.304 | 0.005 | 0.140 |
| AFAP1L2 | 11.56 | 3.55 | 3.76 | 1.25 | 0.307 | 0.030 | 0.332 |
| LOC1005 | 5.17 | 1.59 | 0.49 | 0.00 | 0.309 | 0.002 | 0.000 |
| CALHM2 | 3.47 | 1.07 | 7.34 | 0.54 | 0.309 | 0.048 | 0.073 |
| SRC | 13.25 | 4.09 | 17.35 | 4.13 | 0.309 | 0.036 | 0.238 |
| SPART | 15.36 | 4.75 | 6.49 | 1.33 | 0.309 | 0.001 | 0.205 |
| STK10 | 29.09 | 8.99 | 31.84 | 7.31 | 0.309 | 0.001 | 0.230 |
| MTA3 | 1.77 | 0.55 | 2.74 | 0.68 | 0.309 | 0.004 | 0.249 |
| MAST4-A | 1.34 | 0.42 | 0.70 | 0.12 | 0.310 | 0.003 | 0.167 |
| CPNE8 | 2.31 | 0.72 | 0.64 | 0.12 | 0.311 | 0.036 | 0.188 |
| CRYBG2 | 3.82 | 1.19 | 24.98 | 6.26 | 0.313 | 0.024 | 0.251 |
| CFLAR | 1.55 | 0.49 | 3.87 | 1.25 | 0.315 | 0.029 | 0.322 |
| RAI1 | 11.06 | 3.50 | 12.82 | 2.53 | 0.316 | 0.000 | 0.198 |
| PHOSPHO | 1.49 | 0.47 | 0.80 | 0.13 | 0.317 | 0.003 | 0.166 |
| PLEKHA4 | 2.68 | 0.85 | 3.21 | 0.73 | 0.317 | 0.026 | 0.227 |
| FHL2 | 14.34 | 4.55 | 8.05 | 1.67 | 0.317 | 0.001 | 0.207 |
| PDGFC | 5.53 | 1.78 | 2.01 | 0.47 | 0.322 | 0.001 | 0.235 |
| KCTD14 | 23.22 | 7.49 | 0.76 | 0.00 | 0.323 | 0.004 | 0.000 |
| CHST15 | 4.27 | 1.38 | 4.25 | 1.18 | 0.323 | 0.009 | 0.277 |
| CMTM3 | 2.13 | 0.69 | 0.65 | 0.18 | 0.323 | 0.001 | 0.276 |
| LINC023 | 2.84 | 0.93 | 1.80 | 0.49 | 0.326 | 0.004 | 0.271 |
| IRF2BPL | 14.67 | 4.84 | 17.72 | 3.36 | 0.330 | 0.002 | 0.189 |

**Supplemental Table 2C. Genes induced by XMU.**

Induced genes table sorted based on XMU treated siC 3-fold change over DMSO treated siC and p<0.2.

| Gene Name | FPKM siC | FPKM siC + XMU | FPKM siYap/siTaz | FPKM siYap/siTaz + XMU | XMU Fold Change in siC cells | p value | XMU Fold change in siYap/siTaz cells |
|---|---|---|---|---|---|---|---|
| ZNF887P | 0.48 | 2.58 | 0.32 | 0.69 | 5.315 | 0.000 | 2.176 |
| SMIM14 | 1.09 | 3.56 | 4.55 | 3.45 | 3.257 | 0.000 | 0.758 |
| GADD45B | 22.08 | 66.85 | 33.23 | 167.84 | 3.028 | 0.000 | 5.051 |
| SNHG9 | 2.52 | 8.52 | 4.50 | 6.16 | 3.382 | 0.000 | 1.369 |
| WBP1LP2 | 0.81 | 4.88 | 0.75 | 3.30 | 6.007 | 0.001 | 4.381 |
| EMC1-AS | 0.13 | 1.56 | 0.58 | 3.30 | 12.274 | 0.001 | 5.691 |
| LMO2 | 1.52 | 8.75 | 0.36 | 0.15 | 5.761 | 0.001 | 0.409 |
| SNORD14 | 8.74 | 27.14 | 2.93 | 4.26 | 3.105 | 0.001 | 1.454 |
| ARL6IP1 | 126.31 | 432.75 | 72.51 | 149.68 | 3.426 | 0.001 | 2.064 |
| ALKBH5 | 39.59 | 136.18 | 22.33 | 24.96 | 3.440 | 0.001 | 1.118 |
| PKD1P3-[illegible] | 0.56 | 1.70 | 0.41 | 0.47 | 3.032 | 0.001 | 1.140 |
| OCEL1 | 4.45 | 14.64 | 6.14 | 4.95 | 3.289 | 0.002 | 0.807 |
| RBM15 | 7.58 | 23.72 | 2.18 | 6.20 | 3.127 | 0.002 | 2.848 |
| TUBB4B | 795.59 | 3015.83 | 522.85 | 1221.87 | 3.791 | 0.002 | 2.337 |
| POLR2A | 28.04 | 89.86 | 26.21 | 49.90 | 3.205 | 0.002 | 1.904 |
| ZRSR2P1 | 0.52 | 4.17 | 1.66 | 1.30 | 8.009 | 0.002 | 0.780 |
| H2BC4 | 1.51 | 9.40 | 65.06 | 37.99 | 6.208 | 0.003 | 0.584 |
| BCL2L11 | 2.41 | 8.11 | 4.57 | 6.08 | 3.358 | 0.003 | 1.330 |
| CRY2 | 2.51 | 12.71 | 3.59 | 6.08 | 5.059 | 0.003 | 1.694 |
| SNORD14 | 12.88 | 38.94 | 7.03 | 9.99 | 3.024 | 0.004 | 1.422 |
| TRIM36 | 1.26 | 5.73 | 1.30 | 2.14 | 4.565 | 0.004 | 1.646 |
| H2AX | 106.17 | 398.51 | 36.61 | 92.10 | 3.754 | 0.004 | 2.516 |
| SERTAD1 | 6.83 | 36.69 | 32.82 | 135.05 | 5.375 | 0.004 | 4.115 |
| SYNJ1 | 1.31 | 4.13 | 2.61 | 2.79 | 3.143 | 0.004 | 1.068 |
| RHOB | 2.59 | 10.57 | 26.72 | 84.95 | 4.075 | 0.004 | 3.180 |
| SNORA31 | 0.58 | 2.00 | 1.19 | 1.47 | 3.480 | 0.005 | 1.240 |
| ASB16 | 0.33 | 1.20 | 0.34 | 0.63 | 3.592 | 0.005 | 1.812 |
| LOC1079 | 0.20 | 1.14 | 2.22 | 1.96 | 5.592 | 0.005 | 0.887 |
| BRD2 | 1.54 | 5.14 | 1.88 | 5.45 | 3.336 | 0.005 | 2.899 |
| TMED7-T | 0.13 | 2.14 | 0.79 | 3.12 | 16.515 | 0.005 | 3.961 |
| H1-10 | 28.64 | 103.92 | 64.23 | 95.61 | 3.629 | 0.005 | 1.489 |
| LOC1053 | 1.41 | 5.29 | 1.87 | 4.30 | 3.744 | 0.006 | 2.306 |
| ASNS | 30.18 | 186.48 | 3.50 | 8.47 | 6.178 | 0.007 | 2.423 |
| PPM1L | 0.47 | 1.66 | 0.53 | 0.77 | 3.561 | 0.007 | 1.432 |
| SORBS1 | 0.43 | 1.98 | 0.09 | 0.61 | 4.651 | 0.007 | 6.561 |
| TMEM91 | 0.25 | 1.00 | 0.30 | 0.27 | 4.053 | 0.007 | 0.887 |
| RNVU1-1 | 0.52 | 3.54 | 1.24 | 13.29 | 6.828 | 0.007 | 10.698 |
| PKMP3 | 0.00 | 1.85 | 0.13 | 2.87 | 18519.197 | 0.008 | 21.954 |
| ARHGEF2 | 0.28 | 1.09 | 0.03 | 0.06 | 3.903 | 0.008 | 1.840 |
| WDR47 | 4.59 | 14.91 | 6.76 | 7.60 | 3.246 | 0.008 | 1.124 |
| HAP1 | 0.13 | 1.01 | 0.30 | 1.03 | 7.593 | 0.008 | 3.443 |
| LOC1027 | 0.20 | 1.47 | 0.38 | 0.48 | 7.450 | 0.008 | 1.271 |

| | | | | | | | |
|---|---|---|---|---|---|---|---|
| H2BC12 | 39.77 | 372.07 | 186.78 | 351.21 | 9.356 | 0.008 | 1.880 |
| ZCCHC24 | 0.71 | 3.70 | 1.32 | 3.81 | 5.249 | 0.009 | 2.878 |
| PKIA | 0.35 | 1.38 | 0.11 | 0.30 | 3.898 | 0.009 | 2.793 |
| CASTOR2 | 0.37 | 2.48 | 4.66 | 4.65 | 6.723 | 0.010 | 0.998 |
| SAT1 | 67.71 | 326.78 | 134.48 | 190.24 | 4.826 | 0.010 | 1.415 |
| SNORD14 | 5.69 | 24.08 | 3.43 | 9.73 | 4.229 | 0.010 | 2.832 |
| TESK2 | 1.26 | 5.69 | 1.35 | 1.87 | 4.526 | 0.011 | 1.380 |
| DNHD1 | 0.44 | 1.64 | 0.45 | 1.43 | 3.709 | 0.011 | 3.207 |
| TAF9 | 37.49 | 188.54 | 24.16 | 23.65 | 5.029 | 0.011 | 0.979 |
| CNNM4 | 6.07 | 24.14 | 9.33 | 18.69 | 3.977 | 0.011 | 2.002 |
| BCORL1 | 1.02 | 3.91 | 1.22 | 2.15 | 3.831 | 0.012 | 1.757 |
| LFNG | 0.94 | 3.59 | 0.77 | 0.93 | 3.839 | 0.013 | 1.199 |
| NXF1 | 34.29 | 167.80 | 46.34 | 161.12 | 4.894 | 0.013 | 3.477 |
| LOC1053 | 0.08 | 1.33 | 0.66 | 4.36 | 16.336 | 0.014 | 6.623 |
| HEXIM1 | 25.08 | 89.28 | 35.80 | 78.84 | 3.560 | 0.014 | 2.202 |
| HSPA1B | 7.08 | 21.61 | 8.82 | 12.17 | 3.054 | 0.015 | 1.380 |
| LOC1053 | 0.27 | 1.08 | 1.52 | 2.53 | 4.068 | 0.015 | 1.663 |
| NOS1AP | 0.30 | 1.33 | 0.15 | 0.26 | 4.414 | 0.015 | 1.774 |
| HES6 | 3.41 | 14.84 | 1.98 | 4.08 | 4.353 | 0.016 | 2.062 |
| RBM14 | 19.85 | 88.34 | 15.93 | 41.60 | 4.451 | 0.016 | 2.612 |
| FBXO10 | 0.39 | 1.17 | 0.38 | 1.24 | 3.006 | 0.017 | 3.291 |
| SERPINI | 0.78 | 8.34 | 1.83 | 5.58 | 10.733 | 0.018 | 3.059 |
| H1-2 | 12.18 | 40.08 | 40.06 | 48.58 | 3.291 | 0.018 | 1.213 |
| IRF7 | 4.81 | 30.19 | 10.48 | 14.54 | 6.277 | 0.018 | 1.387 |
| MAP2 | 0.49 | 1.87 | 0.51 | 0.66 | 3.860 | 0.018 | 1.286 |
| FBXO33 | 3.07 | 9.26 | 8.32 | 9.33 | 3.016 | 0.020 | 1.122 |
| NFYA | 4.67 | 14.30 | 3.61 | 6.87 | 3.061 | 0.021 | 1.905 |
| H4C9 | 0.23 | 1.83 | 0.61 | 5.97 | 8.111 | 0.021 | 9.706 |
| H2AC6 | 5.47 | 30.49 | 97.67 | 80.71 | 5.575 | 0.021 | 0.826 |
| H2BC11 | 4.17 | 15.42 | 23.24 | 23.02 | 3.695 | 0.021 | 0.991 |
| SKI | 6.09 | 25.73 | 6.96 | 11.81 | 4.222 | 0.022 | 1.697 |
| TRIM62 | 2.42 | 7.84 | 5.73 | 2.63 | 3.248 | 0.022 | 0.458 |
| RSRP1 | 4.16 | 13.45 | 5.78 | 14.30 | 3.233 | 0.025 | 2.472 |
| KHDRBS3 | 0.51 | 2.99 | 0.09 | 0.82 | 5.898 | 0.025 | 9.048 |
| RPL17P5 | 0.39 | 1.58 | 0.17 | 2.56 | 4.038 | 0.026 | 14.650 |
| MEX3B | 0.27 | 1.65 | 0.21 | 0.55 | 6.211 | 0.026 | 2.584 |
| GUCA1B | 0.54 | 3.78 | 0.73 | 3.37 | 6.955 | 0.026 | 4.647 |
| LOC1079 | 0.25 | 1.35 | 0.43 | 0.73 | 5.508 | 0.027 | 1.722 |
| VAMP1 | 2.40 | 8.26 | 2.35 | 4.41 | 3.448 | 0.027 | 1.873 |
| TMEM52 | 0.72 | 2.42 | 0.30 | 0.39 | 3.371 | 0.027 | 1.282 |
| FAM222A | 0.32 | 2.94 | 0.86 | 3.86 | 9.101 | 0.028 | 4.508 |
| TUBB3 | 42.93 | 140.28 | 75.05 | 142.36 | 3.268 | 0.028 | 1.897 |
| NRARP | 2.65 | 13.95 | 13.46 | 179.97 | 5.268 | 0.028 | 13.375 |
| DGCR6 | 0.42 | 1.32 | 0.27 | 0.00 | 3.169 | 0.029 | 0.000 |
| SNORA51 | 0.20 | 1.05 | 0.37 | 0.63 | 5.198 | 0.029 | 1.711 |
| TRS-GCT | 0.00 | 3.62 | 0.26 | 21.67 | 36230.183 | 0.030 | 82.712 |
| TENT5C | 0.14 | 2.16 | 0.23 | 6.22 | 15.417 | 0.030 | 26.479 |

| | | | | | | | |
|---|---|---|---|---|---|---|---|
| MIR637 | 0.45 | 1.35 | 0.36 | 0.98 | 3.009 | 0.030 | 2.767 |
| RGS2 | 1.89 | 8.61 | 0.62 | 4.78 | 4.550 | 0.030 | 7.656 |
| BICDL1 | 0.86 | 2.80 | 4.26 | 3.47 | 3.269 | 0.031 | 0.815 |
| A1BG-AS | 0.37 | 2.01 | 1.51 | 0.17 | 5.503 | 0.031 | 0.111 |
| MAFK | 3.23 | 11.60 | 5.75 | 5.73 | 3.589 | 0.032 | 0.997 |
| ING1 | 2.53 | 9.55 | 1.77 | 8.60 | 3.768 | 0.032 | 4.844 |
| H2BC21 | 1.37 | 10.31 | 72.55 | 57.80 | 7.523 | 0.033 | 0.797 |
| ZSWIM6 | 2.17 | 8.83 | 1.62 | 3.37 | 4.075 | 0.035 | 2.085 |
| H2BC20P | 2.40 | 15.73 | 4.12 | 14.41 | 6.556 | 0.036 | 3.500 |
| TIGD3 | 0.45 | 2.32 | 0.35 | 0.49 | 5.178 | 0.036 | 1.406 |
| H2BS1 | 0.12 | 1.69 | 1.12 | 1.21 | 14.669 | 0.037 | 1.078 |
| H2BC19P | 2.35 | 18.21 | 3.87 | 15.89 | 7.764 | 0.037 | 4.106 |
| APOLD1 | 0.48 | 2.36 | 0.25 | 0.21 | 4.900 | 0.037 | 0.823 |
| PAQR6 | 0.44 | 1.49 | 2.20 | 5.50 | 3.372 | 0.037 | 2.506 |
| LINC018 | 0.00 | 1.06 | 0.00 | 0.15 | 10619.320 | 0.037 | 1499.307 |
| ID2 | 0.57 | 6.42 | 2.07 | 24.00 | 11.318 | 0.038 | 11.613 |
| CCDC17 | 0.37 | 1.45 | 0.39 | 4.65 | 3.923 | 0.039 | 11.975 |
| H3C10 | 1.41 | 5.92 | 2.79 | 3.92 | 4.211 | 0.039 | 1.403 |
| MIDN | 18.99 | 78.58 | 49.52 | 159.65 | 4.138 | 0.039 | 3.224 |
| H2BC5 | 2.34 | 40.70 | 58.53 | 77.84 | 17.357 | 0.040 | 1.330 |
| H4C15 | 0.93 | 9.43 | 2.60 | 3.98 | 10.136 | 0.040 | 1.531 |
| BAMBI | 5.20 | 16.25 | 6.63 | 12.10 | 3.126 | 0.040 | 1.825 |
| MIR3153 | 0.92 | 4.44 | 0.00 | 0.59 | 4.844 | 0.040 | 5918.128 |
| RND1 | 0.41 | 2.92 | 2.30 | 2.92 | 7.029 | 0.041 | 1.270 |
| LOC1053 | 0.14 | 1.81 | 9.41 | 21.65 | 13.013 | 0.041 | 2.300 |
| METRNL | 1.01 | 4.76 | 14.43 | 5.31 | 4.721 | 0.042 | 0.368 |
| TUBB2B | 1.91 | 5.87 | 10.79 | 16.19 | 3.074 | 0.042 | 1.500 |
| SNORA78 | 1.38 | 5.42 | 0.33 | 0.82 | 3.920 | 0.043 | 2.446 |
| MIR6758 | 3.55 | 13.34 | 4.56 | 6.79 | 3.762 | 0.044 | 1.488 |
| MIR22HG | 1.40 | 4.95 | 8.25 | 6.91 | 3.525 | 0.045 | 0.838 |
| NATD1 | 0.75 | 2.86 | 5.92 | 4.52 | 3.791 | 0.046 | 0.764 |
| DHRS2 | 0.61 | 3.33 | 0.72 | 1.38 | 5.460 | 0.046 | 1.907 |
| ADARB1 | 1.10 | 3.64 | 0.99 | 0.74 | 3.297 | 0.048 | 0.741 |
| FGF18 | 0.09 | 1.08 | 0.04 | 2.28 | 11.452 | 0.048 | 59.293 |
| RPL7AP4 | 0.36 | 1.21 | 1.08 | 1.75 | 3.366 | 0.048 | 1.626 |
| LRP4-AS | 0.17 | 1.36 | 0.08 | 0.34 | 8.169 | 0.049 | 4.481 |
| VGF | 0.40 | 2.01 | 0.27 | 12.30 | 5.012 | 0.051 | 44.962 |
| H2AC18 | 5.74 | 36.29 | 148.84 | 149.11 | 6.322 | 0.051 | 1.002 |
| H2AC19 | 5.93 | 36.55 | 194.77 | 149.25 | 6.165 | 0.052 | 0.766 |
| H2BC6 | 0.12 | 1.24 | 0.96 | 25.66 | 10.296 | 0.052 | 26.704 |
| PEG10 | 34.73 | 220.31 | 7.33 | 28.93 | 6.343 | 0.054 | 3.944 |
| CSKMT | 3.96 | 16.15 | 1.78 | 8.34 | 4.076 | 0.054 | 4.684 |
| EPHB3 | 4.76 | 14.96 | 36.90 | 24.45 | 3.140 | 0.055 | 0.662 |
| PREX1 | 0.05 | 1.67 | 0.00 | 0.10 | 35.256 | 0.055 | 1010.119 |
| H4C14 | 3.69 | 20.23 | 3.23 | 6.91 | 5.487 | 0.056 | 2.140 |
| SLC6A8 | 24.76 | 75.26 | 18.28 | 52.35 | 3.039 | 0.056 | 2.865 |
| CAPN5 | 0.41 | 2.73 | 2.33 | 3.03 | 6.660 | 0.063 | 1.298 |

| | | | | | | | |
|---|---|---|---|---|---|---|---|
| KCNE5 | 0.24 | 1.16 | 0.09 | 0.08 | 4.835 | 0.065 | 0.937 |
| LOC1087 | 0.06 | 1.62 | 3.24 | 15.60 | 25.924 | 0.067 | 4.815 |
| NPHP3 | 0.56 | 2.23 | 0.00 | 0.18 | 3.968 | 0.068 | 61.706 |
| LINC011 | 0.60 | 3.18 | 1.82 | 4.48 | 5.261 | 0.071 | 2.464 |
| PFDN6 | 3.09 | 11.28 | 7.02 | 6.51 | 3.652 | 0.071 | 0.927 |
| H3C2 | 2.88 | 9.24 | 0.75 | 1.47 | 3.211 | 0.071 | 1.967 |
| H3C14 | 1.00 | 4.37 | 1.32 | 6.14 | 4.387 | 0.075 | 4.652 |
| H2BC8 | 3.76 | 18.78 | 135.57 | 229.29 | 4.991 | 0.076 | 1.691 |
| ITPR1 | 0.63 | 1.90 | 0.35 | 0.46 | 3.020 | 0.076 | 1.290 |
| RRAD | 0.14 | 1.04 | 0.25 | 1.42 | 7.418 | 0.076 | 5.771 |
| H3C15 | 0.98 | 4.31 | 1.60 | 6.07 | 4.392 | 0.076 | 3.797 |
| FAM117A | 0.63 | 1.93 | 0.54 | 3.01 | 3.055 | 0.077 | 5.554 |
| DGAT1 | 4.89 | 14.77 | 3.89 | 8.28 | 3.019 | 0.077 | 2.130 |
| MIR548A | 0.00 | 1.02 | 0.41 | 0.24 | 10223.427 | 0.078 | 0.587 |
| LOC1005 | 0.00 | 1.73 | 0.17 | 24.15 | 17312.363 | 0.081 | 140.989 |
| KLHL38 | 0.23 | 1.08 | 0.05 | 0.11 | 4.666 | 0.081 | 2.298 |
| MXD1 | 2.53 | 7.76 | 16.68 | 13.72 | 3.071 | 0.084 | 0.823 |
| AVPI1 | 30.14 | 90.66 | 18.18 | 32.03 | 3.008 | 0.093 | 1.762 |
| TMEM47 | 0.27 | 1.10 | 0.13 | 0.35 | 4.112 | 0.095 | 2.597 |
| H3C4 | 1.26 | 14.21 | 11.93 | 20.73 | 11.238 | 0.095 | 1.738 |
| CCN1 | 32.25 | 97.76 | 0.85 | 12.33 | 3.031 | 0.098 | 14.470 |
| HMGN2P3 | 10.59 | 32.12 | 1.97 | 2.49 | 3.034 | 0.101 | 1.265 |
| BFSP1 | 0.12 | 1.12 | 0.24 | 0.19 | 9.152 | 0.101 | 0.817 |
| KCNJ14 | 0.15 | 2.60 | 0.27 | 0.93 | 17.347 | 0.102 | 3.447 |
| SNORD45 | 1.42 | 4.51 | 0.67 | 0.80 | 3.170 | 0.106 | 1.208 |
| LOC1079 | 0.38 | 2.40 | 0.31 | 2.99 | 6.304 | 0.107 | 9.563 |
| NFKBIB | 2.09 | 7.78 | 9.54 | 3.08 | 3.731 | 0.108 | 0.323 |
| SNORA29 | 0.40 | 1.77 | 0.49 | 0.63 | 4.390 | 0.109 | 1.285 |
| FAM229A | 0.55 | 1.97 | 0.94 | 2.78 | 3.604 | 0.121 | 2.959 |
| RPS10P1 | 0.18 | 2.45 | 11.18 | 25.54 | 13.572 | 0.121 | 2.284 |
| CCN2 | 3.04 | 80.89 | 0.29 | 2.33 | 26.574 | 0.122 | 8.050 |
| CASP8AP | 0.07 | 1.33 | 0.27 | 0.20 | 19.731 | 0.124 | 0.746 |
| AGAP4 | 0.09 | 2.34 | 0.39 | 0.19 | 25.135 | 0.127 | 0.494 |
| DPYSL3 | 0.05 | 1.07 | 0.10 | 0.08 | 21.490 | 0.132 | 0.793 |
| GAN | 0.45 | 1.51 | 1.43 | 0.73 | 3.336 | 0.140 | 0.513 |
| LOC1027 | 0.02 | 2.74 | 1.00 | 1.94 | 142.109 | 0.141 | 1.946 |
| LINC000 | 1.66 | 12.11 | 0.03 | 0.23 | 7.294 | 0.142 | 8.096 |
| H3-2 | 0.47 | 1.54 | 0.17 | 0.31 | 3.262 | 0.142 | 1.842 |
| SNORD45 | 0.49 | 2.66 | 0.25 | 0.94 | 5.382 | 0.144 | 3.765 |
| EEF1DP2 | 0.00 | 1.17 | 0.46 | 0.33 | 11676.467 | 0.149 | 0.712 |
| MIR425 | 0.13 | 1.64 | 0.00 | 0.32 | 12.558 | 0.154 | 3189.974 |
| CCDC200 | 0.00 | 1.30 | 0.08 | 0.50 | 12964.611 | 0.155 | 6.660 |
| PIP4K2B | 0.00 | 2.14 | 0.00 | 0.00 | 21445.040 | 0.184 | 1.000 |
| CPM | 0.28 | 1.76 | 1.22 | 2.59 | 6.364 | 0.185 | 2.129 |
| B3GALNT | 0.00 | 2.37 | 0.00 | 0.12 | 23709.273 | 0.185 | 1181.978 |
| HSPE1-M | 0.00 | 1.23 | 0.00 | 0.47 | 12345.180 | 0.186 | 4747.030 |
| CCDC84 | 0.00 | 1.97 | 0.00 | 0.00 | 19726.048 | 0.190 | 1.000 |

| | | | | | | | |
|---|---|---|---|---|---|---|---|
| TMEM106 | 0.35 | 1.05 | 0.16 | 0.57 | 3.023 | 0.191 | 3.658 |

**Supplemental Table 2D. Genes significantly induced by XMU dependent upon YAP/TAZ.** Genes were sorted based on XMU fold change expression of 3-fold over the siC DMSO treated condition (with p<0.2) and whose expression in siYAP/TAZ XMU treated cells was less than 50% of that in siC XMU treated cells.

| Gene Name | FPKM siC | FPKM siC + XMU | FPKM siYap/siTaz | FPKM siYap/siTaz + XMU | XMU Fold Change in siC cells | p value | XMU Fold change in siYap/siTaz cells | Ratio of FPKM in XMU treated siC cells to siYap/siTaz cells |
|---|---|---|---|---|---|---|---|---|
| PIP4K2B | 0.00 | 2.14 | 0.00 | 0.00 | 21445.040 | 0.184 | 1.000 | 0.000 |
| CCDC84 | 0.00 | 1.97 | 0.00 | 0.00 | 19726.048 | 0.190 | 1.000 | 0.000 |
| DGCR6 | 0.42 | 1.32 | 0.27 | 0.00 | 3.169 | 0.029 | 0.000 | 0.000 |
| LMO2 | 1.52 | 8.75 | 0.36 | 0.15 | 5.761 | 0.001 | 0.409 | 0.017 |
| LINC000 | 1.66 | 12.11 | 0.03 | 0.23 | 7.294 | 0.142 | 8.096 | 0.019 |
| CCN2 | 3.04 | 80.89 | 0.29 | 2.33 | 26.574 | 0.122 | 8.050 | 0.029 |
| ASNS | 30.18 | 186.48 | 3.50 | 8.47 | 6.178 | 0.007 | 2.423 | 0.045 |
| B3GALNT | 0.00 | 2.37 | 0.00 | 0.12 | 23709.273 | 0.185 | 1181.978 | 0.050 |
| ARHGEF2 | 0.28 | 1.09 | 0.03 | 0.06 | 3.903 | 0.008 | 1.840 | 0.056 |
| PREX1 | 0.05 | 1.67 | 0.00 | 0.10 | 35.256 | 0.055 | 1010.119 | 0.060 |
| KCNE5 | 0.24 | 1.16 | 0.09 | 0.08 | 4.835 | 0.065 | 0.937 | 0.070 |
| DPYSL3 | 0.05 | 1.07 | 0.10 | 0.08 | 21.490 | 0.132 | 0.793 | 0.073 |
| HMGN2P3 | 10.59 | 32.12 | 1.97 | 2.49 | 3.034 | 0.101 | 1.265 | 0.078 |
| NPHP3 | 0.56 | 2.23 | 0.00 | 0.18 | 3.968 | 0.068 | 61.706 | 0.079 |
| AGAP4 | 0.09 | 2.34 | 0.39 | 0.19 | 25.135 | 0.127 | 0.494 | 0.082 |
| A1BG-AS | 0.37 | 2.01 | 1.51 | 0.17 | 5.503 | 0.031 | 0.111 | 0.083 |
| APOLD1 | 0.48 | 2.36 | 0.25 | 0.21 | 4.900 | 0.037 | 0.823 | 0.088 |
| KLHL38 | 0.23 | 1.08 | 0.05 | 0.11 | 4.666 | 0.081 | 2.298 | 0.105 |
| TAF9 | 37.49 | 188.54 | 24.16 | 23.65 | 5.029 | 0.011 | 0.979 | 0.125 |
| CCN1 | 32.25 | 97.76 | 0.85 | 12.33 | 3.031 | 0.098 | 14.470 | 0.126 |
| PEG10 | 34.73 | 220.31 | 7.33 | 28.93 | 6.343 | 0.054 | 3.944 | 0.131 |
| MIR3153 | 0.92 | 4.44 | 0.00 | 0.59 | 4.844 | 0.040 | 5918.128 | 0.133 |
| LINC018 | 0.00 | 1.06 | 0.00 | 0.15 | 10619.320 | 0.037 | 1499.307 | 0.141 |
| SNORA78 | 1.38 | 5.42 | 0.33 | 0.82 | 3.920 | 0.043 | 2.446 | 0.151 |
| CASP8AP | 0.07 | 1.33 | 0.27 | 0.20 | 19.731 | 0.124 | 0.746 | 0.153 |
| SNORD14 | 8.74 | 27.14 | 2.93 | 4.26 | 3.105 | 0.001 | 1.454 | 0.157 |
| H3C2 | 2.88 | 9.24 | 0.75 | 1.47 | 3.211 | 0.071 | 1.967 | 0.159 |
| TMEM52 | 0.72 | 2.42 | 0.30 | 0.39 | 3.371 | 0.027 | 1.282 | 0.161 |
| BFSP1 | 0.12 | 1.12 | 0.24 | 0.19 | 9.152 | 0.101 | 0.817 | 0.172 |
| SNORD45 | 1.42 | 4.51 | 0.67 | 0.80 | 3.170 | 0.106 | 1.208 | 0.179 |
| ALKBH5 | 39.59 | 136.18 | 22.33 | 24.96 | 3.440 | 0.001 | 1.118 | 0.183 |
| NOS1AP | 0.30 | 1.33 | 0.15 | 0.26 | 4.414 | 0.015 | 1.774 | 0.193 |
| MIR425 | 0.13 | 1.64 | 0.00 | 0.32 | 12.558 | 0.154 | 3189.974 | 0.195 |
| H3-2 | 0.47 | 1.54 | 0.17 | 0.31 | 3.262 | 0.142 | 1.842 | 0.199 |
| ADARB1 | 1.10 | 3.64 | 0.99 | 0.74 | 3.297 | 0.048 | 0.741 | 0.203 |
| TIGD3 | 0.45 | 2.32 | 0.35 | 0.49 | 5.178 | 0.036 | 1.406 | 0.214 |
| PKIA | 0.35 | 1.38 | 0.11 | 0.30 | 3.898 | 0.009 | 2.793 | 0.218 |
| H2AX | 106.17 | 398.51 | 36.61 | 92.10 | 3.754 | 0.004 | 2.516 | 0.231 |
| MIR548A | 0.00 | 1.02 | 0.41 | 0.24 | 10223.427 | 0.078 | 0.587 | 0.235 |

| | | | | | | | | |
|---|---|---|---|---|---|---|---|---|
| ITPR1 | 0.63 | 1.90 | 0.35 | 0.46 | 3.020 | 0.076 | 1.290 | 0.241 |
| LRP4-AS | 0.17 | 1.36 | 0.08 | 0.34 | 8.169 | 0.049 | 4.481 | 0.253 |
| SNORD14 | 12.88 | 38.94 | 7.03 | 9.99 | 3.024 | 0.004 | 1.422 | 0.257 |
| LFNG | 0.94 | 3.59 | 0.77 | 0.93 | 3.839 | 0.013 | 1.199 | 0.258 |
| RBM15 | 7.58 | 23.72 | 2.18 | 6.20 | 3.127 | 0.002 | 2.848 | 0.261 |
| TMEM91 | 0.25 | 1.00 | 0.30 | 0.27 | 4.053 | 0.007 | 0.887 | 0.266 |
| ZNF887P | 0.48 | 2.58 | 0.32 | 0.69 | 5.315 | 0.000 | 2.176 | 0.269 |
| HES6 | 3.41 | 14.84 | 1.98 | 4.08 | 4.353 | 0.016 | 2.062 | 0.275 |
| KHDRBS3 | 0.51 | 2.99 | 0.09 | 0.82 | 5.898 | 0.025 | 9.048 | 0.275 |
| PKD1P3-[illegible] | 0.56 | 1.70 | 0.41 | 0.47 | 3.032 | 0.001 | 1.140 | 0.278 |
| EEF1DP2 | 0.00 | 1.17 | 0.46 | 0.33 | 11676.467 | 0.149 | 0.712 | 0.280 |
| SORBS1 | 0.43 | 1.98 | 0.09 | 0.61 | 4.651 | 0.007 | 6.561 | 0.305 |
| ZRSR2P1 | 0.52 | 4.17 | 1.66 | 1.30 | 8.009 | 0.002 | 0.780 | 0.311 |
| TMEM47 | 0.27 | 1.10 | 0.13 | 0.35 | 4.112 | 0.095 | 2.597 | 0.316 |
| LOC1027 | 0.20 | 1.47 | 0.38 | 0.48 | 7.450 | 0.008 | 1.271 | 0.327 |
| TESK2 | 1.26 | 5.69 | 1.35 | 1.87 | 4.526 | 0.011 | 1.380 | 0.328 |
| MEX3B | 0.27 | 1.65 | 0.21 | 0.55 | 6.211 | 0.026 | 2.584 | 0.331 |
| TRIM62 | 2.42 | 7.84 | 5.73 | 2.63 | 3.248 | 0.022 | 0.458 | 0.335 |
| OCEL1 | 4.45 | 14.64 | 6.14 | 4.95 | 3.289 | 0.002 | 0.807 | 0.338 |
| H4C14 | 3.69 | 20.23 | 3.23 | 6.91 | 5.487 | 0.056 | 2.140 | 0.342 |
| ARL6IP1 | 126.31 | 432.75 | 72.51 | 149.68 | 3.426 | 0.001 | 2.064 | 0.346 |
| MAP2 | 0.49 | 1.87 | 0.51 | 0.66 | 3.860 | 0.018 | 1.286 | 0.350 |
| SNORD45[illegible] | 0.49 | 2.66 | 0.25 | 0.94 | 5.382 | 0.144 | 3.765 | 0.353 |
| AVPI1 | 30.14 | 90.66 | 18.18 | 32.03 | 3.008 | 0.093 | 1.762 | 0.353 |
| KCNJ14 | 0.15 | 2.60 | 0.27 | 0.93 | 17.347 | 0.102 | 3.447 | 0.357 |
| SNORA29 | 0.40 | 1.77 | 0.49 | 0.63 | 4.390 | 0.109 | 1.285 | 0.357 |
| TRIM36 | 1.26 | 5.73 | 1.30 | 2.14 | 4.565 | 0.004 | 1.646 | 0.373 |
| ZSWIM6 | 2.17 | 8.83 | 1.62 | 3.37 | 4.075 | 0.035 | 2.085 | 0.382 |
| HSPE1-M[illegible] | 0.00 | 1.23 | 0.00 | 0.47 | 12345.180 | 0.186 | 4747.030 | 0.385 |
| CCDC200 | 0.00 | 1.30 | 0.08 | 0.50 | 12964.611 | 0.155 | 6.660 | 0.389 |
| NFKBIB | 2.09 | 7.78 | 9.54 | 3.08 | 3.731 | 0.108 | 0.323 | 0.396 |
| SNORD14[illegible] | 5.69 | 24.08 | 3.43 | 9.73 | 4.229 | 0.010 | 2.832 | 0.404 |
| TUBB4B | 795.59 | 3015.83 | 522.85 | 1221.87 | 3.791 | 0.002 | 2.337 | 0.405 |
| DHRS2 | 0.61 | 3.33 | 0.72 | 1.38 | 5.460 | 0.046 | 1.907 | 0.415 |
| H4C15 | 0.93 | 9.43 | 2.60 | 3.98 | 10.136 | 0.040 | 1.531 | 0.422 |
| SKI | 6.09 | 25.73 | 6.96 | 11.81 | 4.222 | 0.022 | 1.697 | 0.459 |
| PPM1L | 0.47 | 1.66 | 0.53 | 0.77 | 3.561 | 0.007 | 1.432 | 0.460 |
| RBM14 | 19.85 | 88.34 | 15.93 | 41.60 | 4.451 | 0.016 | 2.612 | 0.471 |
| CRY2 | 2.51 | 12.71 | 3.59 | 6.08 | 5.059 | 0.003 | 1.694 | 0.478 |
| NFYA | 4.67 | 14.30 | 3.61 | 6.87 | 3.061 | 0.021 | 1.905 | 0.480 |
| IRF7 | 4.81 | 30.19 | 10.48 | 14.54 | 6.277 | 0.018 | 1.387 | 0.482 |
| GAN | 0.45 | 1.51 | 1.43 | 0.73 | 3.336 | 0.140 | 0.513 | 0.485 |
| MAFK | 3.23 | 11.60 | 5.75 | 5.73 | 3.589 | 0.032 | 0.997 | 0.494 |
